# How a Formate Dehydrogenase Responds to Oxygen: Unexpected O_2_ Insensitivity of an Enzyme Harboring Tungstopterin, Selenocysteine, and [4Fe-4S] Clusters

**DOI:** 10.1101/2022.01.18.476765

**Authors:** Joel E. Graham, Dimitri Niks, Grant M. Zane, Qin Gui, Kellie Hom, Russ Hille, Judy D. Wall, C. S. Raman

## Abstract

The reversible two-electron interconversion of formate and CO_2_ is catalyzed by both non-metallo and metallo-formate dehydrogenases (FDHs). The latter group comprises molybdenum-or tungsten-containing enzymes with the metal coordinated by two equivalents of a pyranopterin cofactor, a cysteinyl or selenocysteinyl ligand supplied by the polypeptide, and a catalytically essential terminal sulfido ligand. In addition, these biocatalysts incorporate one or more [4Fe-4S] clusters for facilitating long-distance electron transfer. But an interesting dichotomy arises when attempting to understand how the metallo-FDHs react with O_2_. Whereas existing scholarship portrays these enzymes as being unable to perform in air due to extreme O_2_ lability of their metal centers, studies dating as far back as the 1930s emphasize that some of these systems exhibit formate oxidase (FOX) activity, coupling formate oxidation to O_2_ reduction. Therefore, to reconcile these conflicting views, we explored context-dependent functional linkages between metallo-FDHs and their cognate electron acceptors within the same organism vis-à-vis catalysis under atmospheric conditions. Here, we report the discovery and characterization of an O_2_-insensitive FDH2 from the sulfate-reducing bacterium *Desulfovibiro vulgaris* Hildenborough that ligates tungsten, selenocysteine, and four [4Fe-4S] clusters. Notably, we advance a robust expression platform for its recombinant production, eliminating both the requirement of nitrate or azide during purification and reductive activation with thiols and/or formate prior to catalysis. Because the distinctive spectral signatures of formate-reduced DvH-FDH2 remain invariant under anaerobic and aerobic conditions, we benchmarked the enzyme activity in air, identifying CO_2_ as the bona fide product of catalysis. Full reaction progress curve analysis uncovers a high catalytic efficiency when probed with an artificial electron acceptor pair. Furthermore, we show that DvH-FDH2 enables hydrogen peroxide production sans superoxide release to achieve O_2_ insensitivity. Direct electron transfer to cytochrome *c* in air also reveals that electron bifurcation is operational in this system. Taken together, our work unambiguously proves for the first time the coexistence of redox bifurcated FDH and FOX activities within a metallo-FDH scaffold. These findings have important implications for engineering O_2_-tolerant FDHs and bio-inspired artificial metallocatalysts, as well as for the development of authentic formate/air biofuel cells, modulation of catalytic bias, assessing the limits of reversible catalysis, understanding directional electron transfer, and discerning formate bioenergetics of gut microbiota.

## INTRODUCTION

The simplest carboxylic acid (formic acid), and its conjugate base (formate) are normal products of metabolic activity in living organisms, including bacteria and humans.^1^ However, bacterial aerobic respiration of formate derived from human gut microbiota drives inflammatory dysbiosis.^2^ Although formic acid is primarily used as a food preservative (E236) or as silage additive for maintaining the nutritive value of animal feed,^3^ it is a highly sought-after electron-mediator and feedstock in (electro)microbial bioproduction,^4^ as well as a low carbon-footprint molecule that serves as a chemically robust hydrogen storage medium.^5^ In addition to being a carbon and energy source for the (an)aerobic growth of disparate bacteria,^6^ archaea,^7^ and syntrophic consortia,^8^ formate can be generated abiotically from CO_2_ and renewable electricity.^5^

Formate oxidation and CO_2_ reduction are interconvertible processes that are carried out by prokaryotic formate dehydrogenases (FDHs) (Reaction 1).^9, 10^

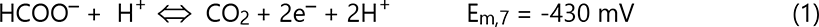

There are two phylogenetically distinct FDH families that can be distinguished by their transition metal ion requirement for enzyme activity.^11^ Metallo-FDHs are thought to be highly sensitive to O_2_,^12, 13^ necessitating catalytic measurements under anaerobic conditions. However, available data in the primary literature are more confusing than definitive. For example, DvH-FDH3 has been reproducibly shown to be O_2_ sensitive^14, 15^ while its ortholog from *D. desulfuricans* ATCC 27774 (Dd) can be purified in air.^16, 17^ Similarly, *D. gigas* (Dg) FDH1 is readily isolated and stored under atmospheric conditions^18, 19^ but its counterpart from DvH requires the presence of 10 mM sodium nitrate to prevent O_2_ inactivation.^20^ Regardless of the purification protocols, the resulting enzymes are ‘dead on arrival’ in that they must be resurrected by lengthy incubations with high concentrations of thiols (10 – 50 mM dithiothreitol^20^ for DvH-FDH1 and 130 mM β-mercaptoethanol^16, 17, 19^ for Dd-FDH3 and Dg-FDH1) and/or formate^17, 21^ prior to catalytic measurements under anaerobic conditions (a representative example can be found in Figure S5a of Oliveira et al^20^). A satisfactory molecular explanation for these phenomenological observations has not been forthcoming for over three decades.^18^ The situation is equally unclear with FDHs isolated from organisms other than sulfate-reducing bacteria (SRB). *Escherichia coli* Fdh-H has been purified and characterized in the presence of sodium azide to minimize O_2_ inactivation.^22, 23^ 10 mM sodium nitrate^24, 25^ or azide^26, 27^ has been added as stabilizers during the isolation of other bacterial metallo-FDHs as well. Very little is known about how these small molecules protect the enzyme from O_2_. Because stability in air does not enable aerobic catalysis, the inhibitors are either removed prior to measurements under anaerobic conditions^20, 25^ or allowed to remain while the activity is probed anaerobically^23^ or in air.^28^ To our knowledge, no FDH has been shown to reversibly interconvert formate and CO_2_ in air. Claims to the contrary have been considered as experimental artifacts.^25^ Moreover, mechanistic details regarding how O_2_ reacts with these metalloenzymes are not available.

Largely overlooked, however, is the fact that the present-day claims of FDH O_2_ sensitivity fail to recognize or rationalize the findings reported between the late 1920s and early-1990s vis-à-vis existence of metallo-FDHs capable of oxidizing formate with oxygen (Reaction 2).

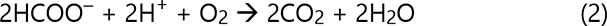

Starting with the first purification of a bacterial FDH by Stickland in 1929, O_2_ uptake served as a proxy for measuring enzyme activity.^29^ Subsequently, Stephenson and Stickland isolated *Escherichia coli* hydrogenlyase (Fdh-H), revealing that it was not responsible for the FOX activity.^30^ A key insight regarding the latter came from Gale’s demonstration that formate dependent O_2_ consumption by *E. coli* was higher in aerobically grown cells.^31^ Pinsent’s groundbreaking study relied on Gale’s O_2_ utilization assay to convincingly show that not only *E. coli* FDH requires molybdenum and selenium for function, but more importantly, that it retained robust FOX activity.^32^ This has since been independently confirmed by several research laboratories.^33–37^ Pichinoty went on to show that FOX activity was broadly distributed across bacteria.^38^ Collectively, these findings paved the way for Sawers to discover the third Fdh-O (O for oxidase; the remaining two being Fdh-N (nitrate)^39^ and Fdh-H (hydrogen)^40^) in *E. coli*.^9, 41, 42^ Although others have confirmed the presence of Fdh-O,^43, 44^ isolation and characterization of a metallo-FDH with FOX activity has remained intractable.^45^ And the possibility that coexistence of dehydrogenase and oxidase activities would render a metallo-FDH insensitive to O_2_ has not been entertained thus far. To tackle this challenge, one of our laboratories (C.S.R.) advanced and tested the hypothesis that FDHs capable of transferring electrons to natural high potential acceptors are likely to be O_2_-insensitive by virtue of their FOX activity, for such physiological reactions are poised to occur under aerobic conditions. Despite the paucity of information regarding redox partners (two well characterized systems exhibit low reduction potentials.^18, 46^), our central hypothesis was inspired by Yagi’s 1969 observation that an FDH from *D. vulgaris* Miyazaki (DvM) preferentially transfers electrons to a high-potential cytochrome *c*_553_.^47, 48^ Because the genetically tractable DvH^49, 50^ is closely related to DvM^51^ and encodes a 73% identical cytochrome *c*_553_ (E_m,7_ = +62 mV),^52^ we probed the O_2_ sensitivity of periplasmic FDHs from this SRB shown to thrive in microaerobic niches.^53–, 55^ We focused on the poorly characterized DvH-FDH2 (locus tag DVU2482-2481)^56, 57^ and cytochrome *c*_553_-reducing DvH-FDH3 (locus tag DVU2812-2809)^14, 15^ instead of the well-studied DvH-FDH1 (locus tag DVU0587-0588)^20^, which couples anaerobic formate oxidation to sulfate reduction by initiating electron transfer to a low-potential cytochrome *c*_3_ (E_m,7_ = −350 mV).^50^ Here, we describe our discovery and characterization of an O_2_-insensitive FDH that retains both dehydrogenase and oxidase activities.

## RESULTS AND DISCUSSION

### Robust Expression Platform for Facile Production of Highly Pure O_2_-Insensitive Metallo-FDHs

There are three distinct *fdh* loci in the DvH genome^57^ (Figure 1A). Only FDH1 encoded by the first locus is essential for growth when sulfate and formate serve as electron acceptor and donor, respectively.^50^ The cellular function of FDH2 and FDH3 are not well defined. Oliveira *et al*^20^ expressed FDH1 in a Δ*fdh1* deletion strain using the vectors and strategies developed in one of our laboratories (J.D.W.). We took a different approach. We reasoned that construction of a markerless FDH-free strain could be beneficial on three fronts: (a) Facilitate biochemical investigations of a native or foreign FDH without potential interference from host counterparts, (b) Benchmark whole cell biocatalysts, and (c) Uncover how synergy between enzyme catalysis and bioenergetics modulates organismal dynamics. To that end, we generated a DvH strain (JW2127; see Methods; Tables S1 and S2) that is devoid of all three *fdh* loci. Although JW2127 is unable to grow on formate-acetate-sulfate, it maintains wild-type-like growth profile on lactate-sulfate medium (Figure 1B). We also constructed deletion strains harboring different combinations of *fdh* genes for functional analyses, including JW2111 (Δ*fdh3*) and JW2121 ( Δ*fdh1* and Δ*fdh3*; see Tables S1 and S2). The latter two served as controls in this study (Figure S1). Subsequently, we used JW2127 for the homologous expression of FDH2. Introduction of a Strep-tag II at the C-terminus of the large subunit facilitated one-step affinity purification. Whereas Oliveira *et al*^20^ used DvH cells derived from 300 L fermentation to purify FDH1, we have streamlined our workflow to produce 1.8 mg of highly-pure heterodimeric FDH2 from a gram of wet cell paste (Figures 2A and 2B). Thus, our 10 L culture (biomass yield of ∼ 8 g) generates sufficient protein to tackle even the most demanding experiments. And our method can be readily scaled up.

**Figure 1.**
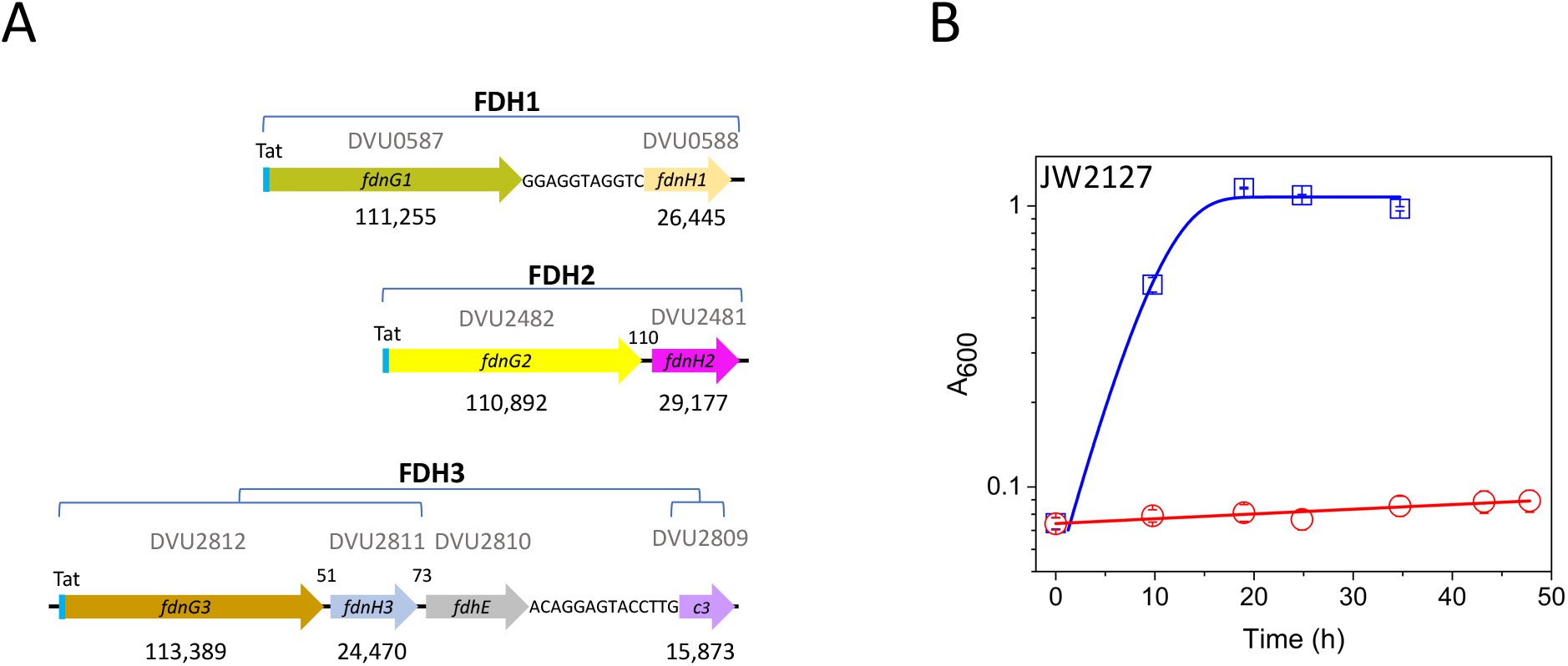
Structure and function of FDH operons in DvH: (A) Condensed map of the three fdh loci. fdnG2 (yellow; large subunit) and fdnH2 (magenta; small subunit) encode FDH2 investigated in this work and are part of a five gene (not shown) operon. Short intergenic regions are illustrated at the nucleotide level while the length of their long counterparts is identified by two-or three-digit numbers. Periplasmic FDH localization is made possible by the twin-arginine translocation (Tat) signal peptide (cyan). Theoretical molecular masses of the encoded polypeptide in daltons are listed below each gene. (B) Growth curves of JW2127: formate-acetate-sulfate (red), lactate-sulfate (blue). The lines going through the points represent fits to Weibull^58^ growth model. Error bars represent standard deviations from three independent measurements.

**Figure 2.**
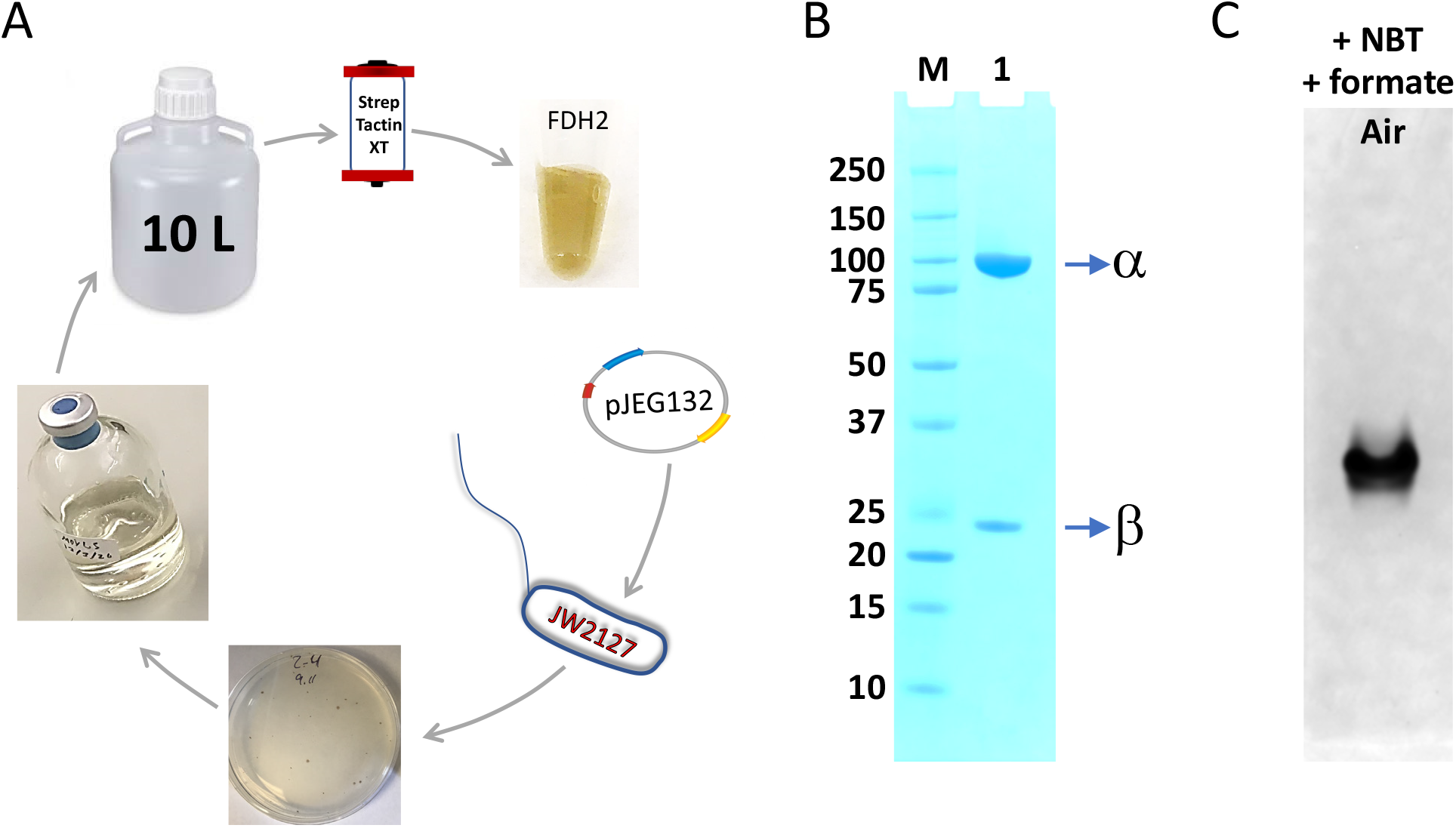
Isolation of DvH-FDH2. (A) Streamlined expression and purification workflow. Single colonies resulting from the transformation of fdh2 plasmid into strain JW2127 were used to start a pre-culture that served as the inoculum for a 10 L scaleup, cells from which were aerobically lysed and subjected to affinity purification, yielding Strep-tagII FDH2. (B) SDS-PAGE of purified protein (lane 1) and molecular weight markers (lane M). α and β represent the large and small subunits of FDH2, respectively. (C) Following non-denaturing PAGE, FDH2 activity (dark single band) is detectable in air via NBT staining.

Notably, there is a fundamental difference between prevailing strategies for metallo-FDH isolation and what we have advanced. Our purification workflow (Figure 2) and downstream handling steps (including storage) occur in air without involving nitrate, azide, thiols, or formate at any stage of the process.

### Aerobic In-Gel Catalysis of Recombinant DvH-FDH2

Literature precedents exist for anaerobic activity staining of FDHs in native polyacrylamide gels using 2,3,5-triphenyltetrazolium chloride^56, 59^ or phenazine methosulfate (PMS) / nitroblue tetrazolium chloride (NBT).^60–62^ However, this has not been achieved for any FDH in air. Because O_2_-insensitive group 5 [NiFe]-hydrogenases have been zymographically visualized using redox dyes,^63^ we asked whether a similar approach could work with DvH-FDH2. When native polyacrylamide gel strips containing recombinant DvH-FDH2 were incubated aerobically with NBT and formate, a single dark blue colored band appeared within two minutes (Figure 2C). In the absence of formate, this band was not observed. The same pattern was recapitulated in the spot assay where the blue color developed within 15 s (Figure S2). These observations demonstrate that electrons released from enzymatic aerobic formate oxidation are readily transferred to an artificial electron acceptor with positive reduction potential (E_m,7_ = +50 mV^24, 63^), resulting in the generation of insoluble reduced NBT-formazan precipitates. Furthermore, our results establish that both nitrate-assisted purification of FDH and/or reductive activation with high concentration of thiols are not essential for maintaining redox activity under anaerobic or atmospheric conditions.

### [4Fe-4S] Metalloclusters, Tungstopterin, and Selenocysteine Remain Unaffected by O_2_ During Catalytic Turnover

As correctly pointed out by Hagen^64^, the metal specificity profiles of SRB FDHs remain incompletely described. Moreover, the nature of redox centers in DvH-FDH2 is unknown.^56^ Because DvH-FDH1 and DvH-FDH2 exhibit 61% protein sequence identity (large subunit) and share all the metal coordination sites within the two subunits (Figure S3), we surmised that a similar complement of redox centers must exist in both systems. Since the DvH biomass was derived from a medium containing Mo (1.24 μM) and W (0.15 μM), we predicted a metal ratio of 1Mo/W:16Fe:1Se. Consistent with this, inductively coupled plasma mass spectrometry (ICP-MS) revealed that for every mole of ^182^W present, another 17± 1 moles of ^56^Fe and 0.7 ± 0.1 moles of ^78^Se were also found (Table S3). Despite the nine-fold excess of molybdate (excluding contributions from yeast extract) in the growth culture, we did not detect ^95^Mo in our FDH2 samples. These results underscore definitive tungsten selectivity of DvH-FDH2, distinguishing it from Mo-specific^14, 56^ DvH-FDH3 and the promiscuous DvH-FDH1, which is capable of incorporating both Mo and W.^20, 56^

### Electronic and Electron Paramagnetic Resonance (EPR) Spectral Signatures of DvH-FDH2 are Virtually Invariant in Air

The bulk of metallo-FDH electronic spectra in the primary literature have been measured under anaerobic conditions to avoid inactivation by molecular O_2_.^13, 16, 20, 24, 65^ Although aerobic spectra exist for an O_2_-tolerant Mo-Cys-FDH stabilized by 10 mM nitrate,^28^ their utility remains unclear, for the addition of formate did not afford a characteristic spectral change. Here, we offer the first functional validation of a W-Sec-FDH in air via electronic spectroscopy. Aerobically purified DvH-FDH2 is brown in color and shows a broad S → Fe^3+^ charge transfer transition at 412 nm (Figure 3A, blue trace), which is characteristic of [4Fe−4S]^2+^ clusters.^66^ Addition of formate leads to a substantial loss of this signal, indicating reduction to the [4Fe−4S]^+^ state (Figure 3A, green trace). Independently, reduction with dithionite yields a similar result (Figure 3A, orange trace). Employing anaerobic conditions makes no difference to the outcome (Figure 3C). The virtually identical lineshape and amplitude of the difference spectra (Figure 3B,D) illustrate that formate completely reduces (six reducing equivalents; four [4Fe−4S]^2+^clusters and one W center) the majority of catalytically competent FDH2 in solution. As dithionite would be expected to reduce both functional and non-functional metal centers, we conclude that >94% of DvH-FDH2 is functionally fit. We have also obtained the source DvH-FDH1 spectrum (Figure S4, pink trace, of Oliveira et al (2020)) and compared it with our as-isolated DvH-FDH2 counterpart acquired under anaerobic conditions (Figure S4). The A_400_/A_280_ ratio –an indicator of the extent of cluster loading^67^– estimated from these spectra are 0.18 (DvH-FDH2) and 0.17 (DvH-FDH1), affirming that the two orthologs exhibit comparable protein purity and cofactor integrity.

**Figure 3.**
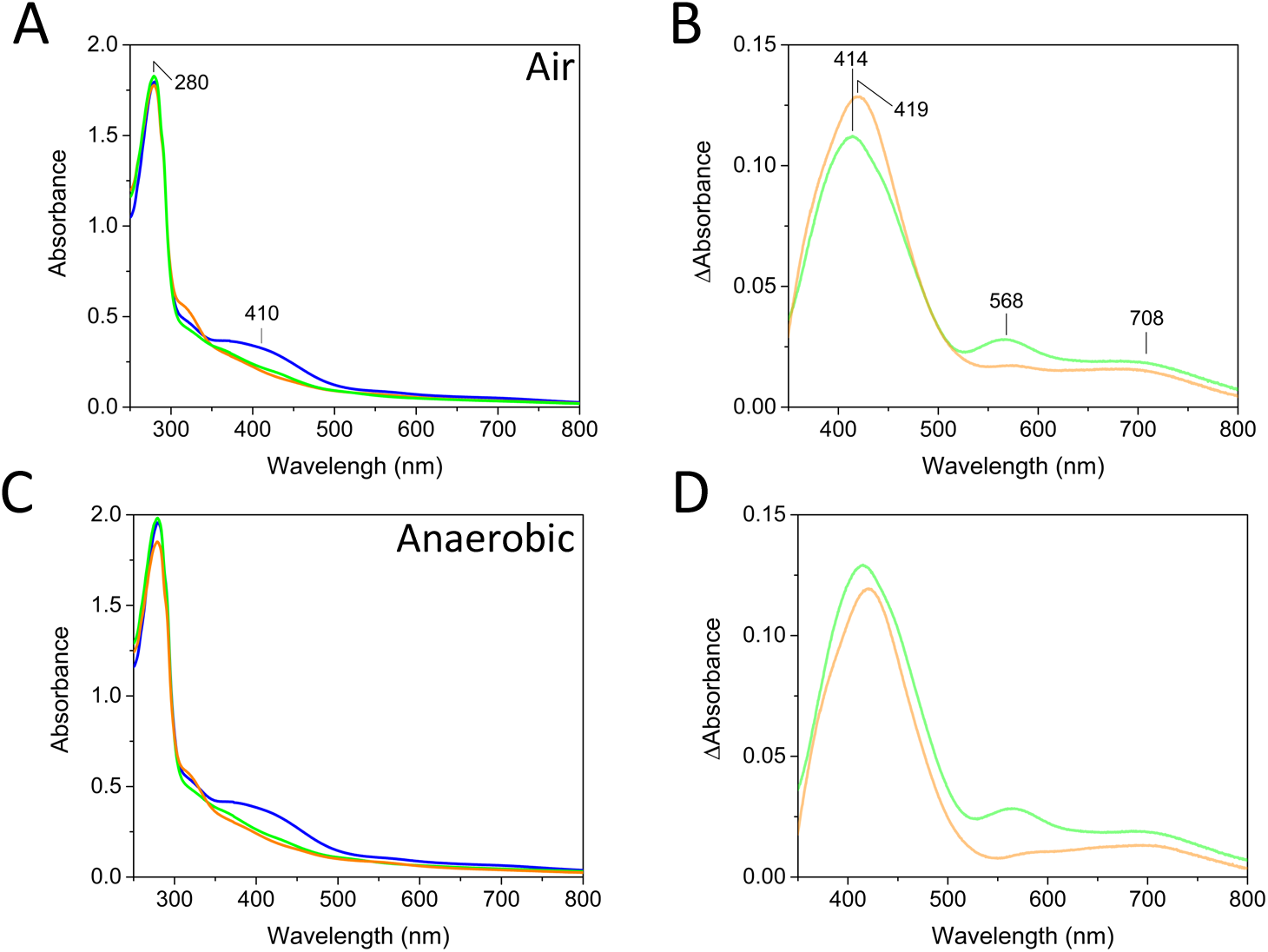
Electronic spectra of DvH-FDH2. As-isolated (blue), formate-reduced (green), and dithionite-reduced (orange) states are shown in panels A and C. Difference spectra are shown in panels B and D. As-isolated minus formate-reduced and as-isolated minus dithionite-reduced are in green and orange, respectively.

To evaluate the predictions made via electronic spectroscopy, we pursued EPR measurements. Figure 4 shows the EPR spectra seen with reduced DvH-FDH2 under a variety of conditions, as well as the spectra for the as-isolated protein (panels (i), (v)). At 15K, we observed predominantly two distinct EPR signals (Figure 4A(ii)-(iv)), the relative intensities of which are essentially independent of reductant (formate or dithionite) and environment (anaerobic or air). The signals are consistent with the presence of reduced iron sulfur clusters. At 26K, one of the signals is significantly broadened (Figure 4A(vi)-(viii)) while by 40K both signals have disappeared (data not shown). This behavior is consistent with fast relaxing [4Fe-4S] clusters. There is also some indication of additional signals (Figure 4A, red arrows), which are described in more detail below.

**Figure 4:**
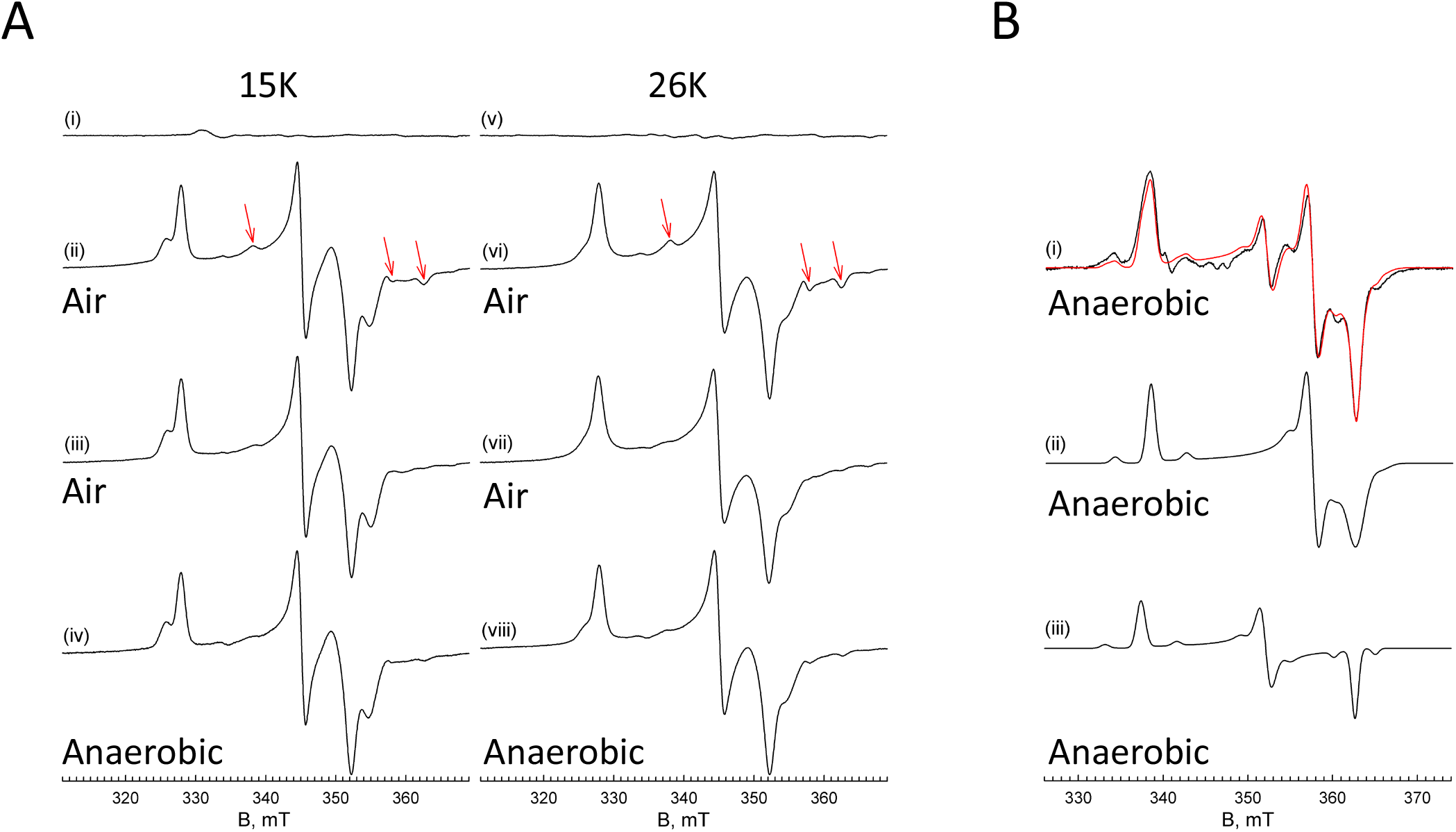
EPR spectra of DvH-Fdh2. (A). (i), (v) as isolated (45 μM). (ii), (vi) dithionite-reduced (45 μM). Red arrows indicate approximate location of W^V^ *g* tensors. (iii), (vii) formate-reduced aerobic (45 μM). (iv), (viii) formate-reduced anaerobic (27 μM). (i)-(iv) were collected with 10 Gauss modulation amplitude and 0.2 mW power at 15K. (v)-(viii) were collected with 10 Gauss modulation amplitude and 4 mW power at 26K. (B) Simulation of the W center of dithionite-reduced sample. (i) W^V^ EPR spectrum (black trace) and simulation (red trace) of 150 μM Fdh2 collected with 8 Gauss modulation amplitude and 10 mW power at 108K. (ii), (iii) Scaled individual contributions to the simulation of the composite spectrum in (i). Simulation includes hyperfine splittings originating from the 14.3% naturally occurring ^183^W (*I* = ½). Simulation parameters are presented in Table S4.

The simulated spectrum for the formate-reduced DvH-FDH2 prepared under aerobic conditions and collected at 15K from Figure 4(iii) is shown in Figure S5, and the simulation parameters given in Table S4. The g-values obtained are again consistent with iron-sulfur clusters and the relative contribution of each cluster is approximately 1:0.75, indicating almost a complete reduction of both centers. At higher microwave power, no indication of additional signals was observed. The absence of additional signals in the four-cluster FDH2 seen here is reminiscent of *D. gigas* W-FDH1 results,^68^ where the two observed signals represent pairs of Fe/S clusters with similar g-values. Alternatively, the remaining two clusters present in DvH-FDH2 may have reduction potentials that are too low to be reduced by formate or dithionite to any appreciable degree.^69^

When 150 μM enzyme is incubated with dithionite under anaerobic conditions for an extended amount of time (∼12 hours or more) and the spectrum is collected at 108K, an additional pair of signals are obtained (Figure 4B(i)); there is no evidence of the Fe/S signals described in Figures 4A and S5 at this temperature. The new signals persist from 15K all the way to 108K without considerable line broadening, consistent with their arising from slowly relaxing W(V) species. Their simulations are superimposed on the experimental spectra. Figure 4B(ii) and (iii) present the component spectra scaled to their contribution to the composite simulation in Figure 4B(i). The simulation parameters are presented in Table S4 and include the well-resolved tungsten *I*=1/2 hyperfine splittings originating from the 14.3% natural abundance ^183^W isotope. The presence of the *I*=1/2 hyperfine splitting is further evidence that these signals arise from the tungsten center rather than additional Fe/S clusters. The simulations indicate that the two species are in an approximate ratio of 1:0.54 and the *g*-values (*g*_1-3_ = 1.982, 1.876, 1.849 and 1.904, 1.849, 1.914, respectively) are in good agreement with those seen from other W-containing enzymes. Somewhat surprisingly, the large anisotropy of the W(V) *g*-values more closely resembles the “low potential” signal for the *P. furiosus* aldehyde ferredoxin oxidoreductase (AOR), which is a member of a different family of tungsten-containing enzyme than the FDHs.^70^ The presence of multiple W(V) signals in a single sample has been seen with a number of W-containing enzymes and may be due to the presence of inactive species in addition to the catalytically competent one, which is a rather common feature of W-containing enzymes.^64^

### Full Progress Curves Reveal High Catalytic Efficiency Under Atmospheric Conditions and Lack of Enzyme Inactivation or Product Inhibition

To the best of our knowledge, solution enzyme kinetics investigations of metallo-FDHs have not directly probed formate depletion or CO_2_ production. Instead, low-potential artificial electron acceptors, most commonly benzyl viologen (BV; E_m,7_ = −360 mV^24^) and methyl viologen (MV; E_m,7_ = −446 mV^24^) for the forward and reverse reactions, respectively, have been routinely used as surrogates to report on catalytic robustness. Although cautions have been raised against trusting kinetic parameters derived from the use of these “*inefficient and slow redox mediators*”^71, 72^, they continue to be favored. Mo-Cys-FDHs offer an alternative by making it possible to track NAD^+^ reduction or NADH oxidation.^24, 28^ Unfortunately, this strategy can be extended only to select metallo-FDHs and it is prone to yield false-positive results when interrogating aerobic CO_2_ reduction with aerotolerant enzymes.^25^ To further complicate matters, FDHs from SRB are in a class of their own (Table S5). Unlike other bacterial FDHs, these retain little to no activity after purification, requiring lengthy reductive activation with high concentration of thiols and/or formate.^17, 20^ Finally, it is impossible to assess the validity or robustness of the published results when experimental data remains inaccessible – we are not aware of a report on metallo-FDH enzymology that has disclosed a complete set of raw absorbance versus time data used to extract kinetics parameters.

To resolve these uncertainties, we explored rigorous and reproducible solution enzyme kinetics approaches capable of yielding results with functional information content. First, we assessed catalytic efficiency with two chemically distinct artificial electron acceptors, one each from the low-(BV) and high-potential (PES/DCPIP) E_m,7_ = +217 mV^24^) categories. Only the latter afforded the ability to acquire kinetics data both in air and under argon. Second, we identified conditions under which full progress curves could be measured. Such an approach is only possible for stable enzymes that catalyze a single-substrate irreversible reaction in the absence of enzyme inactivation or product inhibition (see below).^73–76^ And third, we have simultaneously analyzed several full progress curves using dynamic simulation-based global fitting^75^ to extract k_cat_ and k_cat_/K_m_. This strategy overcomes the limitations of classical steady-state analysis, such as the use of only the first few seconds of data, unreliable initial velocity values, and overparameterization.^77^ To benchmark our models (Schemes 1 and S1), we obtained source BV enzyme kinetics data that formed the basis of Figure S3 and Figure 1C of Maia *et al*^17^ (*Desulfovibrio desulfuricans* FDH3) and Oliveira *et al*^20^ (DvH-FDH1), respectively. Global fitting of the steady-state progress curves from the former work allowed us to recapitulate the published values (Figure S6). Although Oliveira *et al*^20^ only reported results from the initial velocity data, we were able to extract five full progress curves from their source data and perform *de novo* analysis (Figure S7A-C). In addition to finding k_cat_ values in the reported range, our method redefines the K_m_ of DvH-FDH1 to be 4.6 ± 0.3 μM rather than 17 μM (Figure S7D-F). These observations illustrate that our catalytic models are poised to extract reliable kinetic parameters from DvH-FDH2 progress curves.

**Scheme 1.**
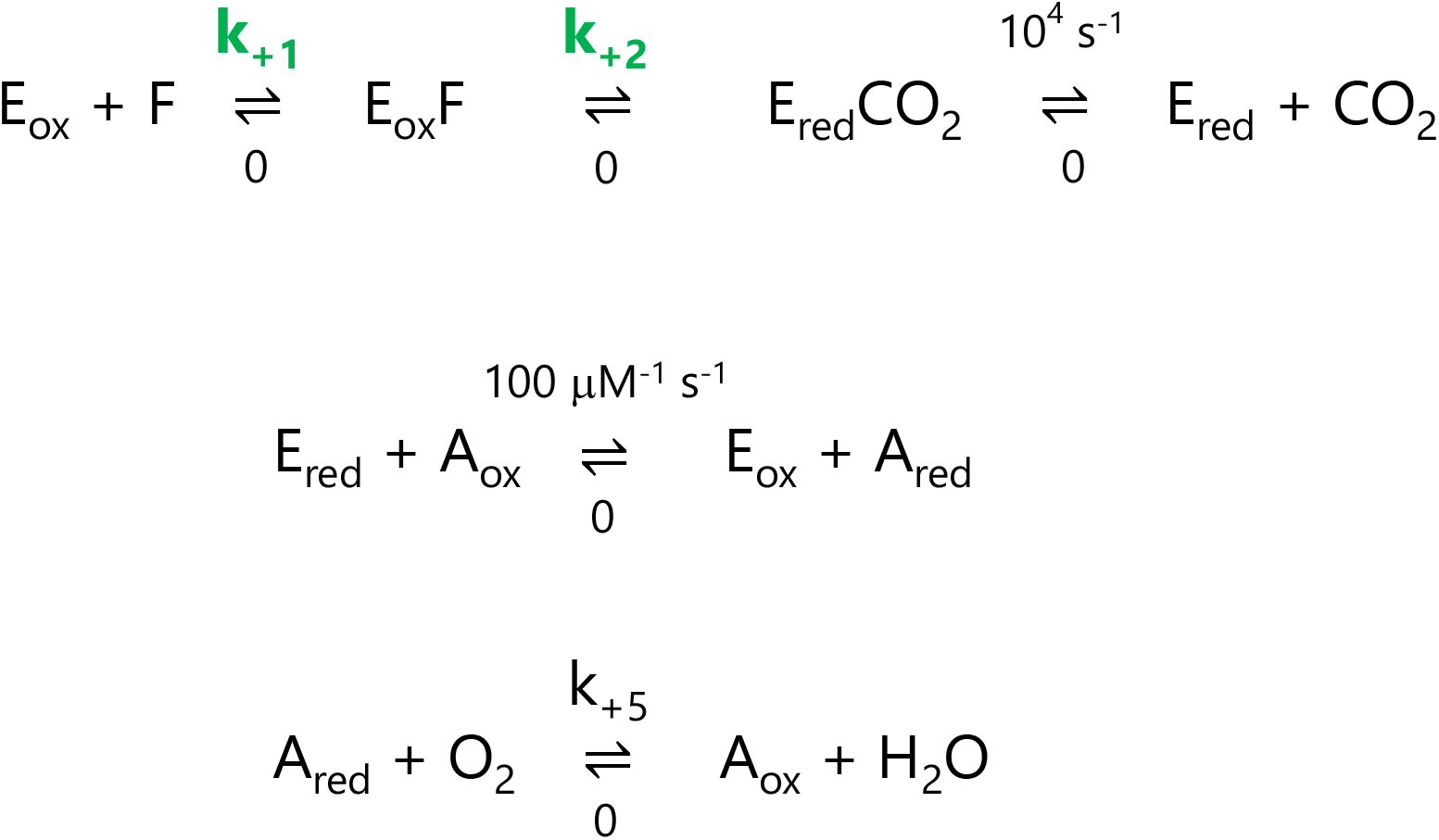
Minimal catalytic model used solely for estimating k_cat_ / K_m_ (**k+1**) and k_cat_ (**k+2**) from full progress curves via dynamic simulation-based global fitting. E_ox_ and E_red_ represent oxidized and reduced enzyme, respectively. A_ox_ and A_red_ denote oxidized and reduced forms of the electron acceptor, respectively. F, formate. The rate of product release (k_+3_) is set to a high value, facilitating the reaction chemistry to define k_cat_. Reaction of E_red_ with A_ox_ (k_+4_) is assumed to be diffusion limited. Rationale for setting reverse rate constants to zero has been explained by Johnson.^77^ The last equation is relevant only under aerobic conditions. Also, see Scheme S1.

Because the original characterization of native DvH-FDH2 –by the same laboratory that has reported extensively on DvH-FDH1– was done using 2 mM BV (see Table S5),^56^ we attempted to reproduce the published results with aerobically purified recombinant DvH-FDH2. However, our enzyme was added to the reaction mix without any pre-activation with thiols or formate. Although DvH-FDH2 displays redox activity in air (Figure 2C), abiotic reaction of reduced BV^+^ with O_2_ made us employ strict anaerobic conditions. Since our measurements were not made inside an anaerobic chamber, we ensured anoxic conditions by adding 1 unit/mL of glucose oxidase (GO). Catalase was also included to eliminate H_2_O_2_ that may arise from GO activity. Global fitting of full progress curves (1,2,4, and 6 μM formate) yielded values comparable to those reported by da Silva *et al*^56^ (Figure 5A-C, Table 1). Furthermore, our progress curves revealed that two molecules of BV^+^ are generated for every formate molecule oxidized by DvH-FDH2. Despite elimination of the reductive activation step, the enzymatic parameters derived from our standard steady-state kinetics analysis were virtually identical to the published values (Table S6), suggesting that the pre-activation step introduced by da Silva *et al* (2011) had no effect on the outcome. In our hands, DvH-FDH2 exhibits catalytic parameters that are roughly an order of magnitude smaller than their DvH-FDH1 counterparts (Table S6 and Figure S7F). A closer inspection of product stoichiometry at high formate concentrations suggested that BV concentration could be limiting (Figure S8). Therefore, we repeated the experiments at 20 mM BV. This restored 2BV^+^:1F stoichiometry across the board but catalytic parameters did not change appreciably (Figure S9, Table S6). We have also confirmed that addition of GO and catalase do not interfere with the results of activity measurements (Figure S10).

**Figure 5.**
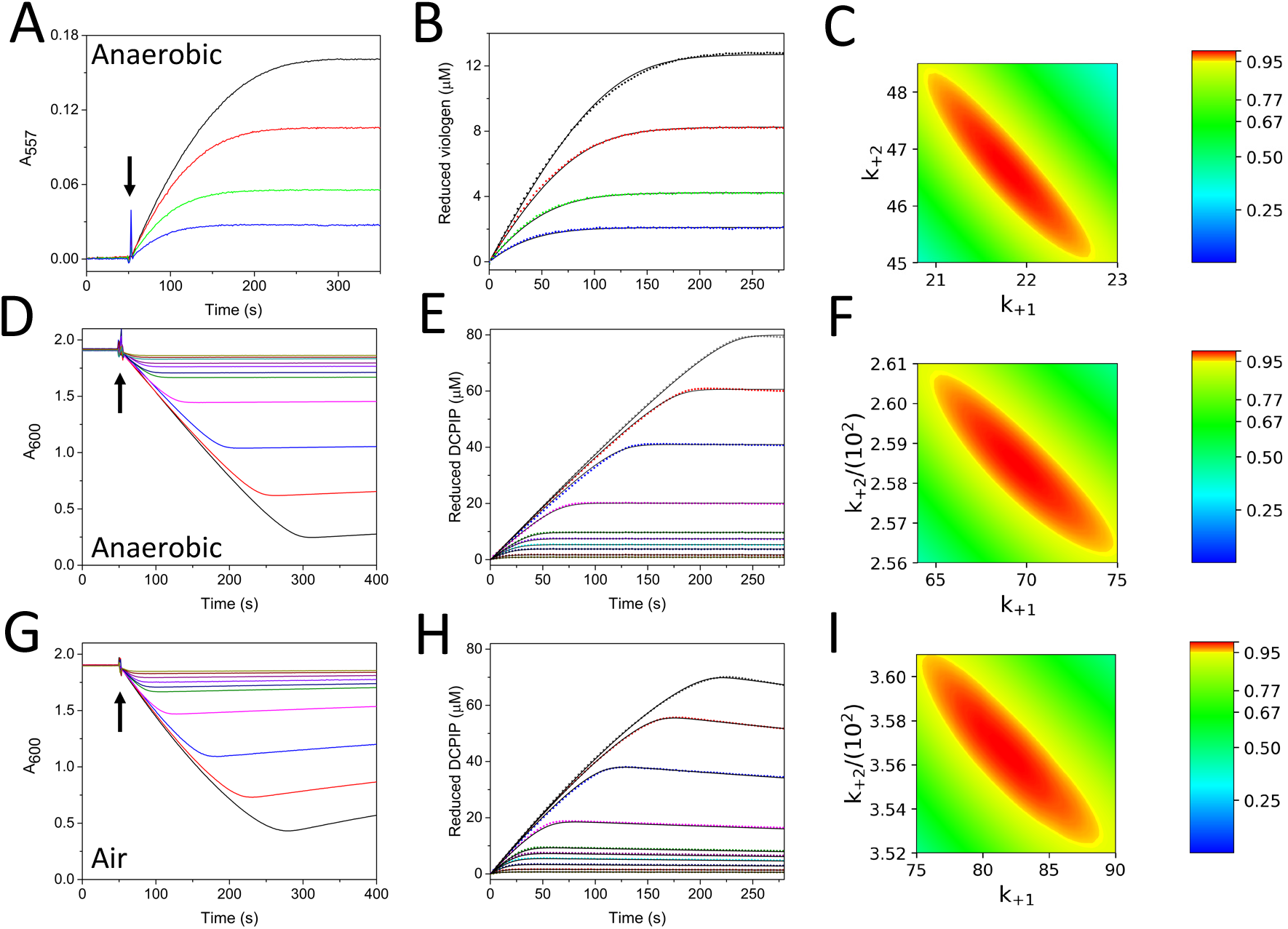
Full progress curve probing of DvH-FDH2 catalysis. Raw experimental traces are shown in A (BV), D (PES/DCPIP), and G (PES/DCPIP) panels. Arrows represent the point at which the experiments were started by either the addition of formate (panel A) or FDH2 (panels D and G). Data normalized for electron acceptor concentration (B, E, and H panels) were globally fit (solid lines) using Kintek Explorer. Whereas panel B was fit to model shown in Scheme S1, panels E and H were fit to the counterpart in Scheme 1. Confidence contour analysis (panels C, F, and I) illustrates that both k_+1_ (k_cat_/K_m_) and k_+2_ (k_cat_) are well constrained by the kinetic data. Upper and lower bounds of each rate constant is reflected by the axes labels. When plotted as a function of one another, red ovals signify the extent of variability in k_+1_ and k_+2_ while still being constrained by the model. Consequently, both parameters display a defined boundary (red), χ^2^ of which is 0.95 (side bar). Table 1 lists rate constants, as well as best fit parameters derived from this analysis.

**Table 1.** Parameters gleaned from full progress curve analysis#

| Electron acceptor | $k_{\text{cat}}$ ($\text{s}^{-1}$) | $K_m$ ($\mu\text{M}$)<sup>‡</sup> | $k_{\text{cat}}/K_m$ ($\mu\text{M}^{-1} \text{s}^{-1}$) | Product stoichiometry<sup>§</sup> | Enzyme inactivation | Product inhibition |
| --- | --- | --- | --- | --- | --- | --- |
| 2 mM BV (anaerobic) | $47 \pm 2$ | $2.1 \pm 0.1$ | $22 \pm 1.1$ | 2BV<sup>+</sup>:1F | No | No |
| DCPIP (anaerobic) | $258 \pm 3$ | $3.7 \pm 0.3$ | $69 \pm 6$ | 1DCPIP:1F | No | No |
| DCPIP (air) | $354 \pm 5$ | $4.5 \pm 0.4$ | $79 \pm 6$ | 1DCPIP:1F | No | No |

<sup>#</sup>Errors for $k_{\text{cat}}$ and $k_{\text{cat}}/K_m$ via confidence contour analysis implemented in FitSpace Explorer
<sup>‡</sup>Standard error values for $K_m$ (calculated from $k_{\text{cat}}$ and $k_{\text{cat}}/K_m$ ) estimated according to Johnson (2019)
<sup>§</sup>F, formate; BV<sup>+</sup>, reduced benzylviologen

Collectively, the observations above suggest that BV is not a good electron acceptor for DvH-FDH2. To test this hypothesis, we independently pursued activity assays with PES/DCPIP. Consistent with literature precedents on dehydrogenases,^78^ phenazine ethosulfate (PES; E_m,7_ = +65 mV^79^) was required for transferring electrons from FDH to DCPIP (Figure S11A). By varying the concentrations of DCPIP and PES, we were able to identify optimal conditions that would support activity measurements both in air (Figure S11B-F and Figure S12) argon (Figure S13). At DCPIP concentrations below 100 μM, global fitting of anaerobic full progress curves (Scheme 1 and Figure 5D-F) resulted in roughly five-fold higher turnover number (TN) than what we obtained with 2 mM BV (Table 1). K_m_ values did not show a significant difference, however. The same pattern was reproduced when the measurements were made in air (Figure 5G-I). A key difference between the two conditions is that the slow reoxidation (k_+5_ = 3 ± 0.5 x 10^-6^ μM^-1^ s^-1^; see last equation in Scheme 1) of reduced DCPIP by O_2_ caused the post reaction region to slope slightly upward (Figure 5G, Figure S11D, and Figure S12B). However, this should not be confused with alterations to the shape of progress curves stemming from product inhibition,^75^ which we did not observe when DCPIP (Figure 5G) or BV (Figure 5A) served as electron acceptors. In fact, a product stoichiometry of one reduced DCPIP for every formate oxidized was reproducibly found in our measurements (Table 1). Since addition of fresh substrate at the end of a progress curve cleanly reproduced the original trace (Figure S12F), we can further ascertain that the enzyme was stable and fully active during catalysis in air. Taken together, PES/DCPIP-dependent catalytic parameters obtained from global fits are in excellent agreement with those from our initial velocity calculations (Table S6). And the TN with PES/DCPIP remains virtually the same under anaerobic and aerobic conditions. Notably, the catalytic efficiency of DvH-FDH2 in air is in the range of 7 x 10^7^ M^-1^ s^-1^ (Table 1), which is comparable to that reported for DvH-FDH1^20^ (∼ 8 x 10^7^ M^-1^ s^-1^) when BV serves as the electron acceptor under anaerobic conditions. Moreover, our PES/DCPIP-based TN and k_cat_/K_m_ for formate oxidation are roughly an order of magnitude and 500-fold higher, respectively, than what has been reported for the aerotolerant Mo-Cys-FDH from *Rhodobacter capsulatus* using the natural (NAD^+^) electron acceptor.^28^ Finally, we have found that the inhibition profiles of DvH-FDH2 in the presence of azide or nitrate are not significantly impacted by O_2_ (Figures S14 and S15). Whereas azide blocks the enzyme with an IC_50_ of about 0.8 mM, nitrate is far less effective.

### Catalytic Redundancy or Gain of a New Enzyme Function? Exploiting the Peck-Gest Paradigm to Seek Insights into How FDHs May Have Evolved to Achieve Aerobic Catalysis

Our full progress curve analysis establishes that both k_cat_ and k_cat_/K_m_ are severely underestimated when BV is used as the electron acceptor. It also reveals a preference for the latter viz., whereas DvH-FDH2 favors high-potential acceptors, such as PES/DCPIP or NBT, DvH-FDH1 is highly active with BV^20^. Although DCPIP data is not available for the latter enzyme, a close ortholog (Dg-FDH1) only shows 10% of BV activity with the high-potential acceptor.^18^ Such linkages take on special significance when multiple FDHs encoded by the same organism are compared. In 1957, Peck and Gest^80^ discovered two types of FDH in *Escherichia coli* solely based on their preference for artificial electron acceptors – one was linked to PMS/DCPIP and its expression was confined to O_2_/nitrate-respiring cells while the other was BV-linked and unique to non-respiring cells (reviewed by Stewart^39^). It is now clear that Fdh-N is DCPIP-linked,^61^ and Fdh-H is BV-linked. The third poorly characterized variant of *E. coli*, Fdh-O, is also DCPIP-linked.^42^ Extending the Peck-Gest paradigm to DvH –only the second microbe for which all three FDHs have now been characterized– we would predict that the BV-linked FDH1 is involved in anaerobic respiration and that the PES/DCPIP-linked FDH2 plays a role in aerobic respiration. It has already been established that FDH1 is essential for anaerobic sulfate respiration when formate serves as the electron donor.^50^ Biological function of FDH2 remains to be elucidated. Our study proves that catalytic parameters derived from viologen-based measurements lack functional information content to make predictions about how well a given FDH would perform under aerobic conditions. Instead, high catalytic performance on BV only guarantees activity under anaerobic conditions. Confirmation bias has boosted reliance on viologen-based kinetics and stymied efforts to uncover O_2_-immune FDHs that can reversibly function in air. This is best exemplified by DvH-FDH2, which exhibits the lowest TN with BV (Table S5) and yet is the most O_2_-insensitive of all metallo-FDHs characterized to date from any bacterium. Therefore, biological context must factor critically into future search efforts aimed at discovering air insensitive FDHs.

### CO_2_ is the Product of Aerobic Formate Oxidation by DvH-FDH2

Although several metallo-FDHs have been investigated, there is just one report in the literature describing the product resulting from enzymatic oxidation of formate under anaerobic conditions.^23^ In all remaining works, product formation is implied based on the reduction of a natural (NAD^+^) or artificial electron acceptor, which is often BV. Although we have used two different artificial electron acceptors in this study, we made sure to leave no stone unturned where product analysis in air is concerned. At pH 7.5, combining DvH-FDH2 with isotopically labeled ^13^C-formate readily yields a discernible H^13^CO_3_**^−^** resonance at 162.93 ppm (Figure 6A, middle and Figure S16) via ^13^C NMR spectroscopy. Addition of PES to the mix substantially enhances the resonance intensity, indicating that the number of turnovers has increased in the presence of an artificial electron acceptor (Figure 6A, top; Figure S17). This hints at the likelihood that O_2_ may not be a good electron acceptor for DvH-FDH2 despite its abundance under our experimental conditions (257 μM at 25 °C). Similarly, at pH 6, both H^13^CO_3_**^−^** (162.88 ppm) and ^13^CO_2_ (127.29 ppm) resonances appear, and their intensities amplify when PES is included (Figure 6B, middle and top, respectively; Figures S18 and S19). As a positive control, we have confirmed that NaH^13^CO3, in isolation at pH 6, generates H^13^CO_3_**^−^** and ^13^CO_2_ resonances at positions identical to those found with enzymatic formate oxidation (Figure S20). These chemical shifts agree well with those reported in the literature.^82 13^C-formate was independently validated by both ^1^H and ^13^C NMR spectra. Whereas the latter generates a single resonance at 173.65 ppm (Figures 6A and 6B, bottom; Figures S21 and S22), *J*-coupling (∼ 195 Hz) between the ^1^H and ^13^C atoms splits ^1^H spectrum of ^13^C-formate into two resonances (8.66 ppm and 8.27 ppm) (Figure S23). Despite the poor sensitivity of ^13^C NMR spectroscopy^83^ and not optimizing data collection for 5x *T_1_* (relaxation time), we succeeded in demonstrating that CO_2_ is the true product of aerobic catalysis.

**Figure 6.**
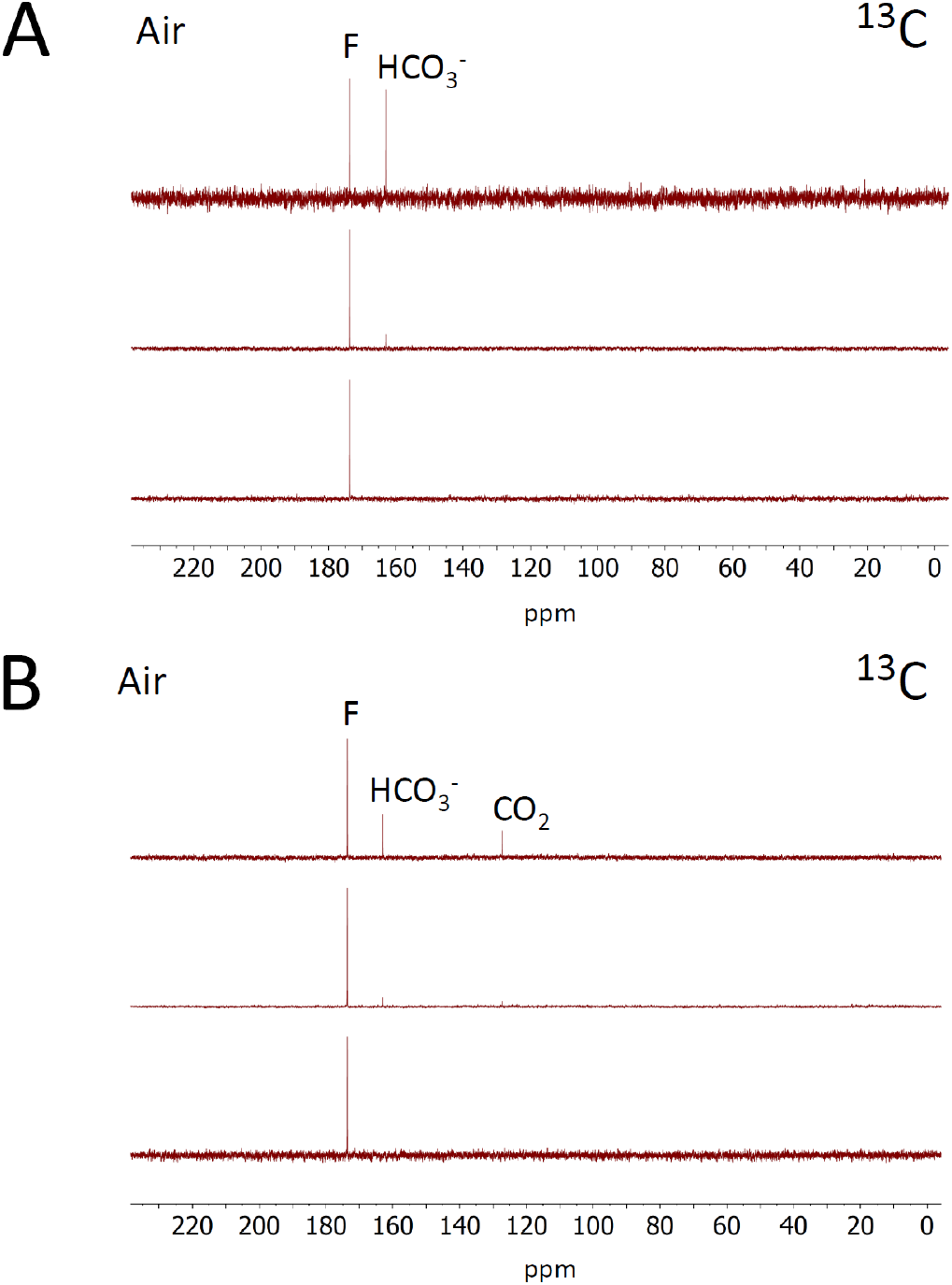
Product generated by DvH-FDH2 catalyzed reaction. ^13^C NMR spectra at pH 7.5 (A) and pH 6 (B). ^13^C-formate (bottom), ^13^C-formate + enzyme (middle), ^13^C-formate + enzyme + PES (top).

### FOX Activity Generates H_2_O_2_, Enabling Oxygen Insensitivity of DvH-FDH2

To understand how DvH-FDH2 deals with O_2_, we used a Clark-type O_2_ electrode to ask whether formate oxidation under atmospheric conditions is coupled to O_2_ reduction. Addition of enzyme to formate-containing aerobic buffer led to robust O_2_ consumption (Figure 7A). Once the signal plateaued, catalase was added. This led to O_2_ evolution followed by O_2_ uptake until a plateau was reached, suggesting the production of H_2_O_2_ via 2e^−^ reduction of O_2_ (reaction 3).

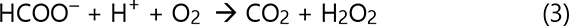

**Figure 7.**
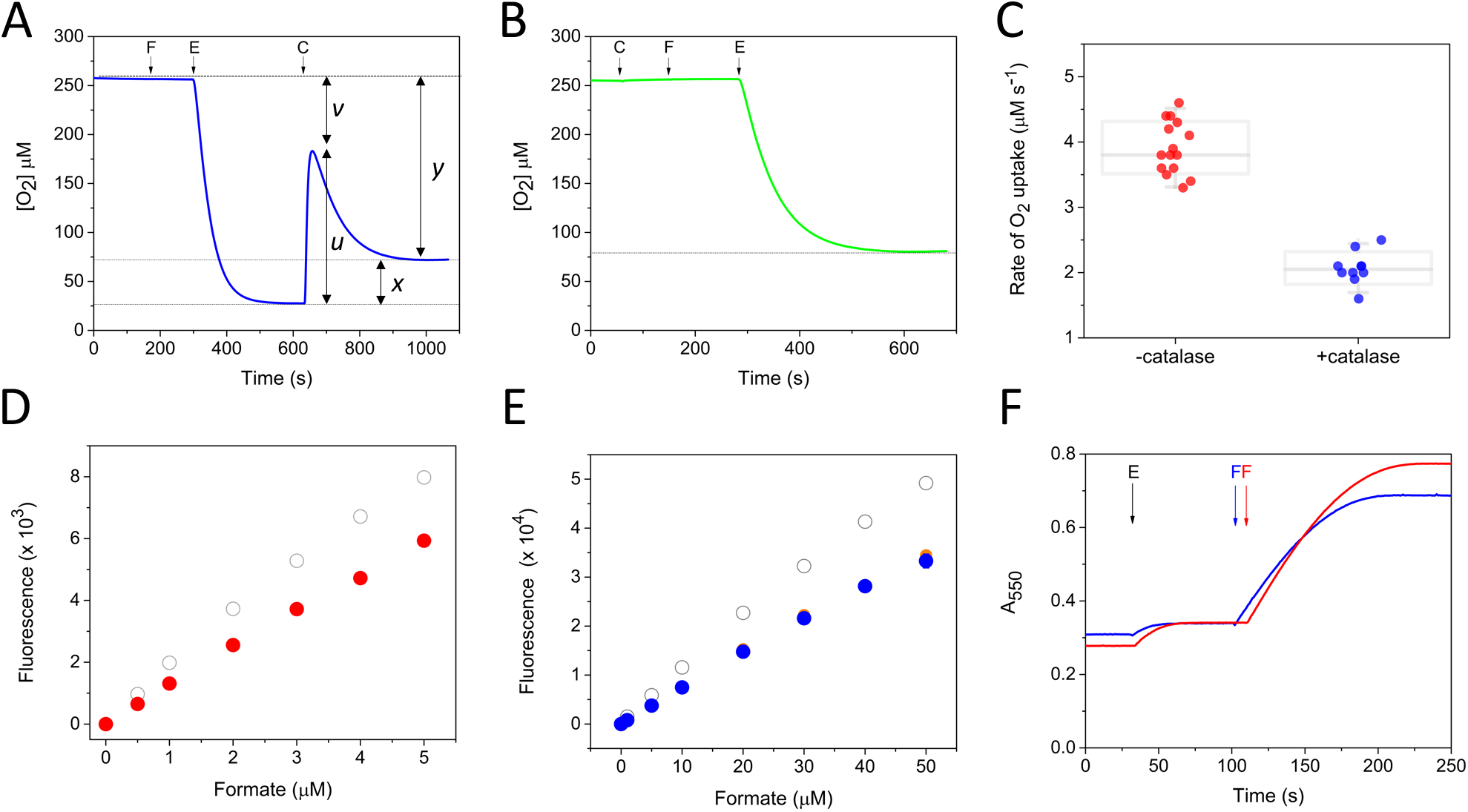
Mechanistic basis of DvH-FDH2 O_2_ insensitivity. (A) Oximetry reveals coupling of formate oxidation to O_2_ reduction. (B) Oximetry in the presence of catalase. (C) Comparison of O_2_ uptake rates in the absence (n = 15) and presence (n = 10) of catalase. Both dot and box plots are shown with the latter deemphasized due to medium sample size.^81^ (D) Enzymatic H_2_O_2_ generation monitored via Amplex Red assay (filled red circles) (n=3). (E) Enzymatic H_2_O_2_ production evaluated using the CBA assay (filled blue circles) (n=3). Filled orange circles represent data obtained at 10 nM FDH2. In both D and E panels, open grey circles represent H_2_O_2_ standard curve determined in the absence of formate or FDH2. (F) Reduction of equine cytochrome c under aerobic (blue) and anaerobic (red) conditions (n = 3). Points of addition of formate (F), enzyme (E = FDH2), and catalase (C), are identified by down arrows. u, v, x, and y are defined in the text.

We attempted a calculation of the electron flux that led to H_2_O_2_ formation.^84, 85^ The *x*/*y* value in Figure 7A would imply that roughly one quarter of the electrons from formate were ending up in H_2_O_2_. However, this is likely to be an underestimate because formate was in large excess and was continuing to be oxidized post dismutation of H_2_O_2_ by catalase, manifesting as the second O_2_ uptake step. Instead, if we considered the *u*/*v* value, ∼ 80% of the electron flux goes towards peroxide generation. To resolve this uncertainty, we pursued the kinetics approach developed by Lu and colleagues.^86^ By comparing the initial velocities of O_2_ uptake in the absence (Figure 7A) and presence (Figure 7B) of catalase, we found that it was 50% lower in the latter (Figure 7C). And the rates did not vary significantly between pH 6.0 and 8. This outcome suggested that H_2_O_2_ was the major product of O_2_ reduction during aerobic formate oxidation. Appropriate controls were built into our experimental design for evaluating alternate endpoints. H_2_O_2_ addition in the absence of exogenous catalase showed that DvH-FDH2 lacks catalase activity (Figure S24A). To rule out the possibility of abiotic O_2_ consumption, DvH-FDH2 was heat denatured and subjected to oximetry. Neither O_2_ uptake nor H_2_O_2_ generation was found (Figure S24B). Moreover, inclusion of 1 mM EDTA minimized artifacts arising from transition metal contaminants. Even with these controls in place, we could not eliminate the possibility that atmospheric O_2_ was completely excluded during oximetry. Therefore, we pursued two orthogonal approaches to directly quantify H_2_O_2_. In the first method, horseradish peroxidase (HRP) catalyzed formation of fluorescent resorufin from H_2_O_2_ and amplex red was monitored. We observed that for every mole of formate oxidized, roughly 0.75 mole of H_2_O_2_ was produced during aerobic DvH-FDH2 catalysis (Figure 7D). Inclusion of catalase abolished the fluorescence signal and denatured enzyme failed to yield H_2_O_2_ (Figure S25). However, the inability of amplex red assay to quantify H_2_O_2_ beyond 5 μM made it impossible for us to investigate the consequences of O_2_ reduction at formate concentrations approaching 10 – 20K_m_. Furthermore, despite being considered the gold standard, this assay is prone to artifacts.^87, 88^ For example, interferences stem from interaction between redox enzymes and resorufin^89^. To overcome these limitations, we resorted to a method independent of HRP and amplex red. Here, we followed the direct reaction of non-fluorescent coumarin-7-boronic acid (CBA) with H_2_O_2_, leading to the production of fluorescent 7-hydroxycoumarin (COH).^90^ Although this reaction is slow^90^ (*k*_on_ 1.5 M^-1^ s^-1^), the assay is linear over a much broader range of H_2_O_2_. Therefore, we quantified H_2_O_2_ production during aerobic DvH-FDH2 catalysis, varying formate concentration between 1 - 10K_m_. It amounted to 64 ± 6% and did not show significant variation when higher enzyme concentrations were used. Catalase addition eliminated the signal completely, confirming that H_2_O_2_ is indeed the major product of O_2_ reduction by DvH-FDH2 (Figure S26).

Next, we tried to assess superoxide (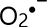) generation^91^ by DvH-FDH2. Because addition of SOD had negligible effect on both direct quantification (Figures S25 and S26) and oximetry (Figure S27), we probed the reduction of partially acetylated cytochrome *c*. The advantage of using the latter is that it is still reducible by 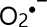 but not susceptible to interferences arising from oxidase or reductase activities when unmodified cytochrome *c* serves as the substrate.^92^ Significant reduction was not observed, implying that 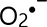 was not released by DvH-FDH2 (Figure S28).

Taken together, our results establish FOX activity of a highly pure metallo-FDH. To the best of our knowledge, this has never been demonstrated before. Based on IUPAC-IUB nomenclature, the term “*oxidase*” is reserved for enzymes, which utilize O_2_ as the electron acceptor. In our case, formate oxidation is coupled to 2e^−^ reduction of O_2_ by DvH-FDH2, resulting in 65 to 75% H_2_O_2_ production (reaction 3). Hence, we project that roughly 25 – 35% O_2_ is reduced to H_2_O by a 4e**^−^** process (reaction 2). From a mechanistic perspective, this is reminiscent of how O_2_-tolerance is achieved in some [NiFe]-hydrogenases (see below)^93^. However, given the high level of difficulty associated with detecting and quantifying H_2_O, only a handful of studies have been performed on this front using redox enzymes.^86, 89, 94, 95^

### Redox Bifurcation Facilitates the Coexistence of FOX and FDH Activities

When DvH-FDH2 couples formate oxidation to the reduction of electron acceptors other than O_2_, it functions as a dehydrogenase. Our kinetic (Figure 5) and product (Figure 6) analysis illustrate that FOX activity does not interfere with the latter function. Nevertheless, it is unknown if this pattern holds when a macromolecule serves as an electron acceptor. Because our central hypothesis rests on the assumption that FOX activity is likely to preserve electron transfer to natural high potential acceptors (see Introduction), we tested it directly. Since an eleven-heme cytochrome *c* (*uhc*) is located immediately adjacent to *fdh2* in DvH^57^, we asked whether DvH-FDH2 reduces native equine cytochrome *c* (E_m,7_ = +260 mV) in the presence of formate. Whereas stoichiometric reduction occurred under anaerobic conditions, only ∼80% underwent reduction in air (Figure 7F). However, the initial rates remained invariant (Figure S29). Doubling the formate concentration resulted in near stoichiometric reduction in air (Figure S30). Moreover, inclusion of SOD or catalase did not result in a noticeable difference (Figure S31). Under the conditions employed, dissolved O_2_ (257 μM) is at a much higher concentration than cytochrome *c*. Yet, electrons are readily delivered to the latter. These results prove for the first time that oxidase and dehydrogenase activities coexist within the DvH-FDH2 scaffold. We advance a non-energy-conserving electron bifurcation (EB)^96^ mechanism to rationalize how this is accomplished (Figure 8). Here, electrons resulting from the oxidation of a 2e^−^ donor (formate) traverse two independent thermodynamically favorable electron transfer paths (FOX and FDH) to reduce two different electron acceptors (O_2_ and cytochrome c). Although we do not know the identity of the cofactor engaged in EB, both the tungstopterin and the proximal [4Fe-4S] cluster of the large subunit are strong candidates. Apropos, arsenite oxidase is thought to achieve EB by cycling between the Mo(VI)-dioxo and Mo(IV)-oxo states.^97^ Whether tungsten can assume this role in DvH-FDH2 remains to be investigated. Based on our observation that superoxide was not generated during aerobic catalysis, we hypothesize that the bifurcating site likely favors normally ordered over inverted potentials.^98, 99^ Even though the interconversion of formate to CO_2_ isreversible, the bifurcation events in either direction (especially O_2_ reduction) are potentially irreversible.^100^ Finally, EB is thought to be the dominant source of reactive oxygen species in biological systems.^100^ We have shown that redox metalloenzymes can capitalize on it to achieve O_2_ insensitivity.

**Figure 8.**
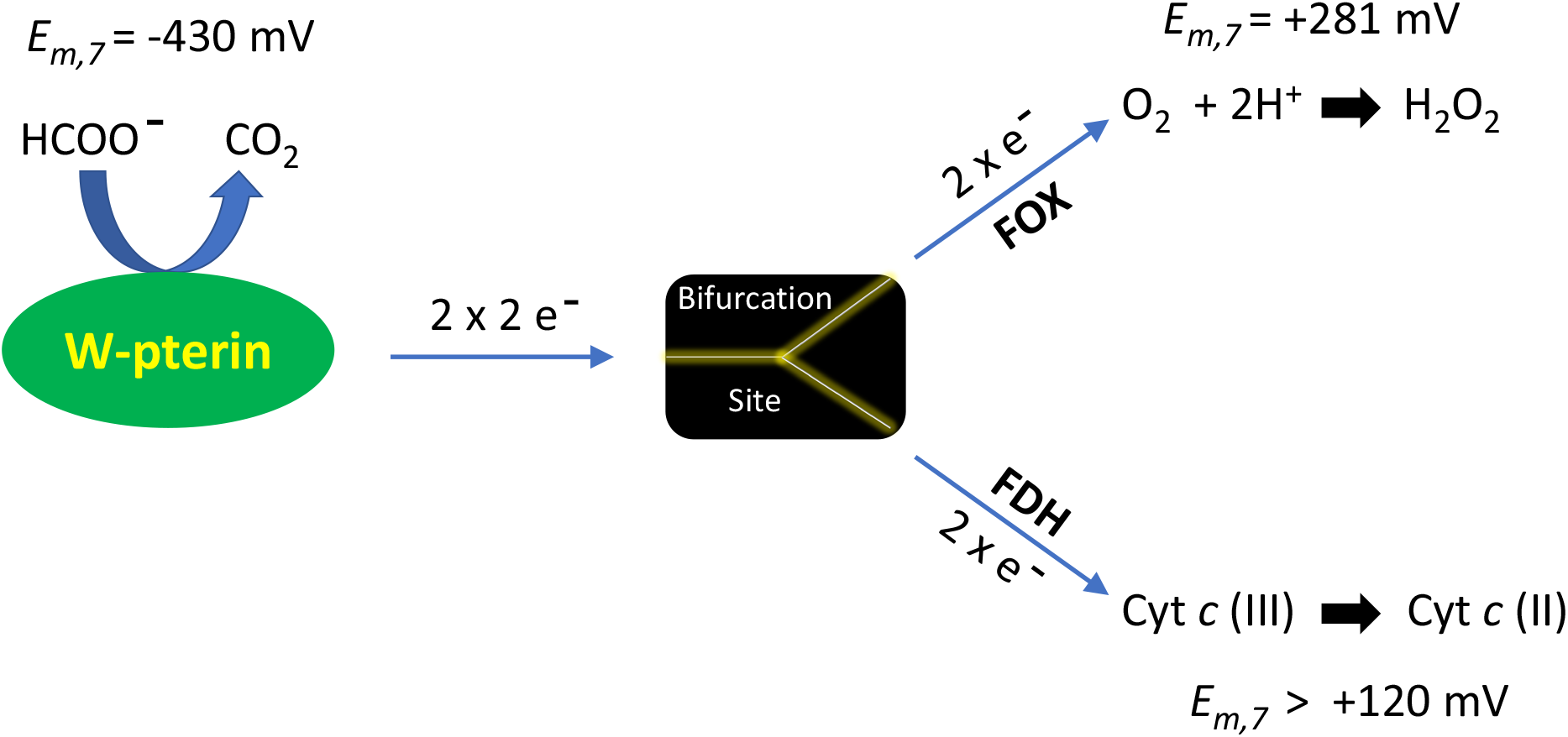
Working model of redox bifurcation by DvH-FDH2. W-pterin denotes the tungstopterin active site. The nature of bifurcation site (black box) is unknown. Since a multiheme cytochrome c (Cyt c) is likely the natural redox partner of DvH-FDH2, it is shown at the end of the bottom branch. However, its reduction potential is a predicted value based on our results with equine Cyt c (E_m,7_ = +260 mV).

### Molecular Basis of DvH-FDH2 O_2_ Insensitivity

In their phylogenetic analysis, Oliveira et al^20^ began with over 6000 FDH sequences and reduced it by an order of magnitude in an effort to understand how the variability impacts catalytic mechanism and O_2_ stability. Here, we have narrowed the sequence space to just two closely related paralogs – one of these (DvH-FDH1) is unable to achieve catalysis in the presence of O_2_ while the other (DvH-FDH2) thrives in air. To gain atomic insights, we constructed a highly accurate heterodimeric structure of the latter using AlphaFold2.1^101^ (Figure 9A). We did not utilize preexisting structure templates to model the structure. The resulting atomic coordinates (sans cofactors) include confidence metrics (predicted Local Distance Difference Test, pLDDT) at the single residue level wherein higher scores on a scale of 1 – 100 represent greater confidence. Figure S32A shows that the heterodimeric structure of DvH-FDH2 is modeled with high confidence; bulk of the polypeptide chain displays pLDDT scores >90. Similarly, the predicted aligned error (PAE; color saturation found at any x,y coordinate in Figure S32B) is a metric of how well a residue is positioned and oriented.

**Figure 9.**
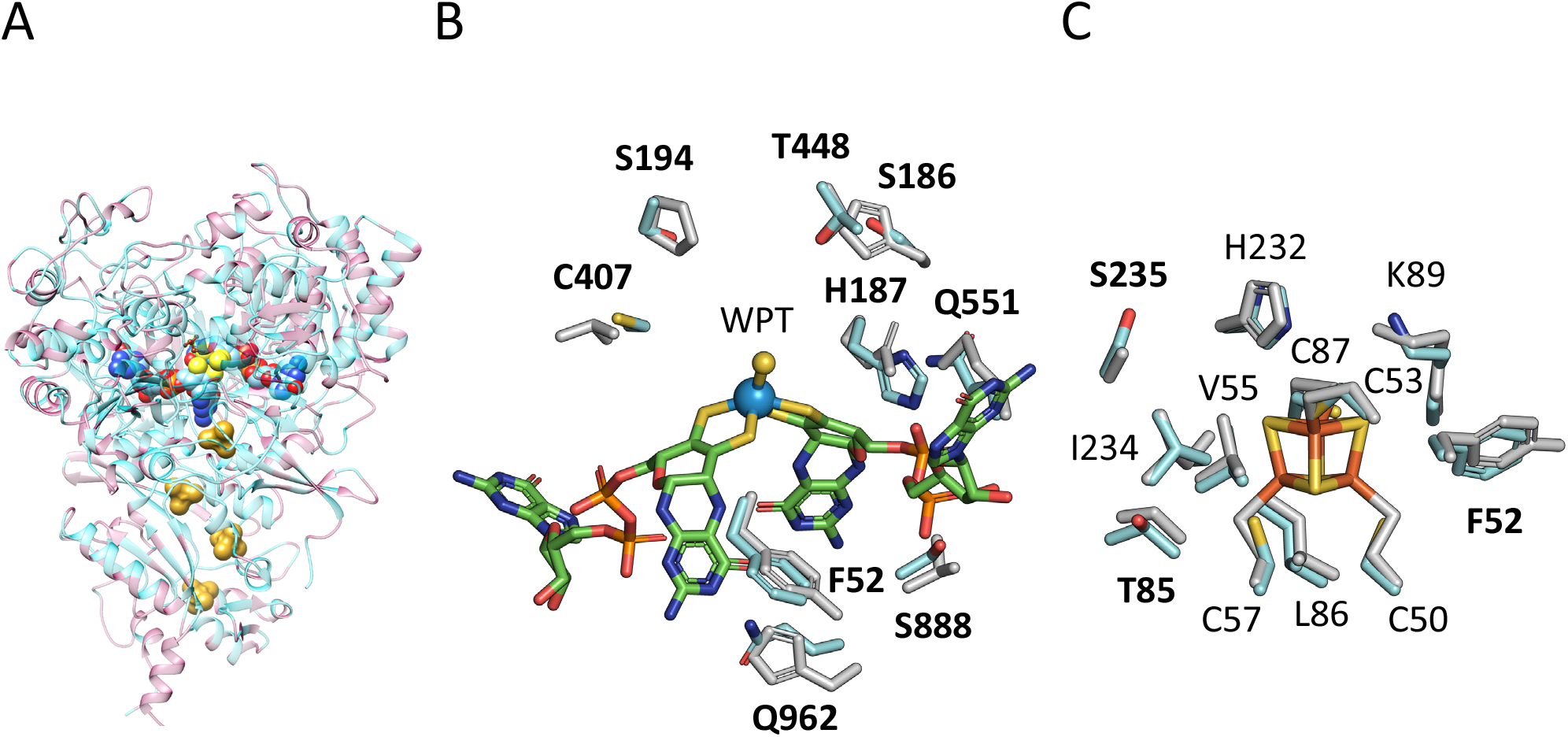
Molecular insights into the O_2_ insensitivity of DvH-FDH2. (A) AlphaFold2-based tertiary topology and quaternary structure. Shading in pink represents sites that are unique to FDH2. The four [4Fe-4S] clusters are depicted by a combination of yellow and orange spheres. The two pterin moieties coordinating a W (WPT) are shown at the top using blue and red spheres. (B) Variability within a 10 Å radius of W. Side chains belonging to DvH-FDH1 are shown in grey. W and sulfide are identified as blue and yellow spheres, respectively. For clarity, conserved sites are not shown (see Figure S36). (C) Environment of the large subunit [4Fe-4S] cluster (yellow-orange sticks). In both panels B and C bolded labels signify variations.

Here too, very low PAE values are seen for both subunits. Independently, we predicted the structures of the two subunits and compared them with the heterodimeric counterpart. Both approaches yielded very similar results. DALI^102^ confirms that the tertiary folds of large and small subunits are superimposable on their DvH-FDH1 counterparts with a root-mean-square-deviation (RMSD) of 1.4 Å (953 Ca atoms; Z-score 58.2; 61% identity) and 1.0 Å (213 Cα atoms; Z-score 35.9; 63% identity), respectively. Backbone RMSD variations are shown in Figures S33 and S34). Similarly, the RMSD between DvH-FDH1 and DvH-FDH2 heterodimers is 1.37 Å (952 Cα atoms). After completing these validations, we computed a residue-residue (RR) distance map to identify the regions of major variation between FDH2 and FDH1 (Figure S35).

Although the active site residues are largely conserved between FDH2 and FDH1, there are several differences in the vicinity of the tungsten center (compare Figure 9B and S36B). Specifically, two highly conserved residues^20^ have been substituted in FDH2. The introduction of S186 and H187 are noteworthy because they replace H187 and Q188, respectively, in FDH1. These changes are likely to influence Sec reactivity. Nonetheless, the relevance of Sec to O_2_ insensitivity is unclear. Whereas it would be expected to confer O_2_ tolerance,^103, 104^ both FDH1 and FDH2 incorporate it and only the latter is capable of aerobic catalysis. Consequently, additional investigations are required to ascertain how Sec contributes differentially to stability and catalysis in enzymatic systems.

[4Fe-4S] clusters are prone to oxidative damage^105, 106^ and enzymes harboring them would be expected to be inactivated by H_2_O_2_ generated during aerobic catalysis.^107^ However, this does not happen with DvH-FDH2. As a corollary, cellular experiments with *Campylobacter* group of bacteria have shown that they harbor an FDH capable of producing H_2_O_2_.^108–110^ In these organisms, H_2_O_2_ functions as a terminal electron acceptor in respiration.^84, 85, 111^ *E. coli* capitalizes on H_2_O_2_ in a similar fashion.^112^ It is unknown whether DvH exploits this strategy. Therefore, we examined the environment of the active site proximal [4Fe-4S] cluster to glean insights. Strikingly, there are three substitutions (Y52F, S85T, A236 → S235) within a 5 Å radius (Figure 9C). This is relevant because in some O_2_-tolerant [Ni-Fe]-hydrogenases, introduction of a polar residue near catalytic metal clusters has been shown to improve O_2_ resistance. Because the predicted structure of DvH-FDH2 requires us to dock the cofactors, we are unable to say more about their conformation or reactivities. Nevertheless, one or more of the following precedents may inform how O_2_ insensitivity and / or EB evolved in our system: (1) group 1d O_2_-tolerant [Ni-Fe]-hydrogenases, which also reduce O_2_ to H_2_O_2_ and H_2_O^89, 113^, generate an unusual [4Fe-3S] cluster; (2) a canonical [4Fe-4S] cluster in another [NiFe] hydrogenase undergoes redox-dependent structural changes,^114^ poising it in a protected state until the next catalytic cycle; (3) second coordination sphere effects; and (4) local or remote conformational fluctuations at the protein level that could offer protection from attack by oxidants. Ongoing structural and biochemical studies are likely to clarify the nature of innovations at work in DvH-FDH2.

### Impact

The findings reported here have broad utility in disparate fields of research. (1) The current state of the art in formate/air biofuel cells is limited to mediated electron transfer because O_2_-sensitive metallo-FDHs need protection from redox polymer films to operate.^115^ Consequently, a “true” formate/air BFC is yet to be fabricated. DvH-FDH2 should be able to work in the absence of protective matrices and power BFCs via direct electron transfer. (2) There is no known biocatalyst that accomplishes CO_2_ reduction in air.^116^ And it is not possible to ground truth this reaction in a chemical environment because low-potential artificial electron donors (e.g., methyl viologen) are futile under atmospheric conditions.^117^ Since the tungstopterin active site and other redox centers of DvH-FDH2 are O_2_ insensitive, there is a high probability of exploiting bioelectrocatalysis to run the reverse reaction aerobically. In so far as the O_2_-tolerant [NiFe] hydrogenase has been successfully shown to perform aerobic bioelectrocatalysis^118–120^ it should be possible to achieve the same for DvH-FDH2. (3) Insights gleaned from DvH-FDH2 should inform tunability of catalytic bias,^121^ limits to electrocatalytic reversibility,^122^ and design of biomimetic metallosynthetics.^123^ (4) Virtually nothing is known about the redox partners of metallo-FDHs in the context of aerobic respiration.^124^ The ability to aerially manipulate DvH-FDH2 will enable strategies for solving problems previously considered intractable in bioenergetics.

## METHODS

### Genetics

#### Construction of *fdh* Deletion Strains

The naming schemes for the *fdh* genes follow the convention established previously^56^. DvH deletion strains (Table S1) were constructed using methods already described^125, 126^. Briefly, for the deletion of each predicted operon, two plasmids were constructed: one to create a marker-exchange deletion and another to remove the marker. Both plasmids are suicide vectors and require at least one homologous recombination event to occur to provide the selectable phenotypes. A phenotypic screen was performed to determine if a double recombination event took place, thereby increasing the likelihood of choosing isolates that had the desired genotype. Each vector contained a cloned copy of at least 300 bp upstream and a similar DNA region downstream of the operon targeted for deletion that were captured in a vector backbone containing the pUC origin of replication and a gene conferring spectinomycin-resistance. The plasmids were constructed by the sequence and ligation independent cloning (SLIC) technique^127^ with amplicons obtained from PCR using the primers found in Table S2 (Integrated DNA Technologies, Coralville, IA) and the Herculase II DNA polymerase (Life Technologies, Grand Island, NY). For the marker-exchange plasmids, the two DNA regions up-and down-stream are separated by an artificial, two-gene operon consisting of *aph(3’)-IIa* (conferring antibiotic resistance to 50 µg kanamycin/mL in *E. coli* and 400 µg G418/mL in DvH) and the counter-selectable marker uracil phosphoribosyltransferase (*upp*, DVU1025) genes. The marker-exchange plasmids were introduced by electroporation into a strain containing a deletion of the *upp* gene and the operon to be deleted. The transformed DvH cells were allowed to recover overnight at 34 °C, as previously described^126^. The cells were then grown for 3-5 days on solidified MOYLS4 medium^128^ containing G418 to select for transformants. Single isolates were screened for sensitivity to 100 µg spectinomycin/mL (consistent with the double homologous recombination event), sensitivity to 40 µg 5-fluorouracil/mL (5FU^S^; to ensure the counter-selection of 5FU resistance (5FU^R^) would be effective) and maintenance of resistance to G418. A putative marker-exchange deletion isolate was then chosen and transformed with the marker-less deletion plasmid, as described above. The transformed cells were recovered, plated on medium containing 5FU and the three phenotypic markers again screened. For the marker-less deletion isolates, however, isolates were selected that were 5FU-resistant and G418-sensitive showing that the marker exchange cassette had been removed from the cell by double homologous recombination. Up to three isolates with the desired antibiotic-resistance phenotype were further analyzed by Southern blot. Once confirmed, one of these isolates was chosen as the marker-less deletion mutant.

For operon deletions, the upstream and downstream regions, respectively, consisted of 858 bp and 806 bp (*fdh1*; DVU0586-0588), 795 bp and 878 bp (*fdh2*; DVU2485-2481), and 976 bp and 970 bp (*fdh3*; DVU2809-2812). Parental strain JW710^126^ was used for the deletion of *fdh1* and *fdh3*. Confirmation by Southern blot was accomplished by digesting the genomic DNA of the parental and putative deletion strains with AgeI (NEB, Ipswich, MA), separating the DNA fragments by gel electrophoresis, and probing with the upstream region.

### Molecular Biology and Microbiology

#### Plasmid Construction

pMO9075 backbone was amplified via Phusion polymerase (New England Biolabs #E0553S) using the primers pMo9075 slic_F and pMo9075 slic_R, separated on 0.6% TAE (BioRad QBI 351-008-131) agarose gel (BioRad 161-3102) and purified via gel extraction (Qiagen #28704). Inserts were amplified with Phusion polymerase via standard reaction conditions. Primers 2482_pmo_F and 2481_pmo_R were used to amplify FDH2 for cloning into pMO9075. Primers 2482_pmo_F and 2482_strII_R were used to amplify DVU2482, introducing the upstream vector flank to DVU2482 and a StrepII tag to the 3’-end of DVU2482. Primers strII_2481_F and 2481_pmo_R were used to amplify DVU2481 with StrepII-tag overlap (while maintaining native intergenic spacer) and downstream vector flank. Amplicons were separated in 0.6 % TAE agarose gels and purified by gel extraction. Inserts were assembled with vector backbone via overlap assembly using Gibson cloning (New England Biolabs #M5510A). Assembly reactions were used to transform *E. coli* α-select chemically competent cells (Bioline BIO-85026) and colonies were selected on YT glucose plus 50 mg/mL spectinomycin HCl (Sigma-Aldrich S9007). For positive clones, 50 mL of transformant was grown in MDAG-11 formulated in house^129^ supplemented with spectinomycin, and the plasmid was purified using a Qiagen Plasmid Midi Kit (Qiagen 12943).

#### Bacterial growth

DvH strains were grown on MOYLS4 medium (see Protocol S1), which was adjusted to pH 7.2 with NaOH. Thioglycolate was added before bottling and after equilibration to dinitrogen (Airgas NI NF200 or research grade). For generating inocula, media were bottled anaerobically under dinitrogen (5 psi), 50 mL per 100 mL serum bottle (Duran Wheaton Kimble 223747) with butyl rubber stopper (Chemglass CLS-4209-14) and aluminum crimp seal (Wheaton 20-0000AS). For larger volumes, glass media bottles (Pyrex 1395500, 13951L; Fisher 06-414-1C/06-41401D) were sealed with no. 6 neoprene stoppers (RPI-259100-6) and capped with media bottle lid with a center bore to access the stopper. Bottles were autoclaved and vitamins were syringed in from a filter sterilized (RPI 256131) 1X stock just before inoculation.

#### Transformation

One of our laboratories (C.S.R.) formulated a protocol that is different from what has been reported earlier.^49^ DvH strains were grown in 50 mL MOYLS4 in 100 mL serum bottle at 37 C with nitrogen headspace to near stationary phase and chilled on ice. Cells were aerobically spun down in a 50 mL conical centrifuge tube (Corning 430828) at 7,500 x g for 5 min, then washed twice in 17 mL of ice cold 15 mM Tris pH 7.2, 10% glycerol supplemented to 1 mM with dithiothreitol. The final pellet was resuspended to 1 mL in the same buffer. A 100 μL aliquot of cells was aerobically mixed on ice with 7.5 μL from plasmid midi prep (∼2-3 μg plasmid) and electroporated at 1.5 kV (1 mm gap cuvette; MBP #5510) in an Eppendorf electroporator 2510. 1 mL of sterile anaerobic MOYLS4 was immediately added and the entire volume was transferred to a bottle of MOYLS4. The bottle was incubated at 37 C (Glascol, Micro-expression Vertiga shaker). Once the culture recovered and became densely turbid, transfers were made to fresh medium containing 100 μg/mL spectinomycin HCl. After two rounds of growth with spectinomycin selection, freezer stocks in 10% glycerol were made. For colony selection the same medium supplemented with 1.5% agar, 5 mM cysteine, 1 mM sodium sulfide, and 100 μg/mL spectinomycin was used and kept in gas tight jars with an AnaeroGen 3.5 L Gas generating system pack (Oxoid). Colonies were picked into selective medium using a sterile 1 mL syringe (Becton Dickinson 309659) fitted with an 18-gauge needle.

#### 10 L carboy growth of DvH-FDH2 producing strain CSR21271

For each carboy, the strain was transferred from 10% glycerol freezer stock in MOYLS4 medium; ∼0.5 mL of stock added to a 50 mL bottle of anaerobic MOYLS4 medium, supplemented with vitamins and 100 μg/mL spectinomycin hydrochloride. Transfers were made by nitrogen purged syringe with 23-gauge needles (Becton Dickison 305190). The culture was incubated overnight at 37 C or until mid-exponential phase of growth. 20 mL of the overnight culture was used to inoculate a 500 mL bottle of MOYLS4 medium, containing vitamins and 100 μg/mL Spectinomycin HCl. The 500 mL culture was incubated overnight at 37 C. 10 Liters of MOYLS4 medium in 2 L bottles, prewarmed, sterile, aerobic, with iron and EDTA withheld, was poured into a sterile 10 L polypropylene carboy (Thermo 2250-0020). The medium was completed by addition of filter sterilized vitamins, spectinomycin hydrochloride (1g dissolved in 15 mL water; 100 μg/mL final) and 4.8 mL of iron chloride (125 mM; Acros 423705000) / EDTA (250 mM; Fisher BP120-1) solution. The carboy was closed and purged with nitrogen via a butyl rubber stopper port (Chemglass CLS-4209-14) affixed to the lid (Figure S37A). 5 mL of 25% sterile neutral sodium sulfide (Alfa Aesar 36622) was then injected through the port and the carboy was mixed by shaking (Figure S37B). Subsequently, the carboy was incubated at 37 C until resazurin indicator turned colorless. The 500 mL culture (OD_550_ ∼0.6) was then transferred into the carboy via sterile rubber transfer line (VWR-62993-726), 18-gauge needles (Becton Dickinson 305196) and under nitrogen pressure (Figure S37C). The carboy was placed in an incubator (Sanyo MCO-17A1C) at 37 C and the optical density (OD) was monitored at 550 nm (Beckman DU-800 spectrophotometer) via 1 mL samples removed from the same port. Once OD_550_ nm plateaued, the carboy was chilled in the cold room overnight (Figure S37D) and then harvested by centrifugation at 8,000 x g (Beckman Avanti HP-26 XPI) in 1 L bottles. Cell pellets were transferred to 50 mL conical centrifuge tubes, respun (7,500 x g; Corning 430828), and then froze at −80 C.

### Biochemistry

#### Protein Expression and Purification

Strep-tag II DvH-FDH2 was purified from strain CSR21271 (see Table S1). Unless specified otherwise all the following steps were done at 4 C and under atmospheric conditions. Nitrate, azide, thiols, or formate were not used at any step of the purification or storage. Cells (∼ 18 g) were suspended in six volumes of 50 mM sodium phosphate (Fluka 71505, Sigma-Aldrich S0786), pH 7.4, containing 150 mM sodium chloride and 1 mL of 50 x Complete Proteinase inhibitor (Roche 45582400; 1 tablet in 1 mL of MilliQ water), by gentle pipetting in cold buffer. Cells were lysed using an Avestin C3 homogenizer and cell debris spun down at 4500 x *g* (Beckman Avanti HP-26 XPI) for 15 min. Membrane vesicles were removed by centrifugation at 285,000 x *g* 1 hr (Beckman Optima L100XP). The clarified lysate was then fractionated by ammonium sulfate precipitation with fractions pelleted at 10,000 x *g* for 10 min and the 40-70% saturating fraction was retained and exchanged via centrifugal concentrator (Amicon 30 kDA molecular weight cutoff) into 100 mM Tris-HCL buffer pH 8, containing 150 mM NaCl and 1 mM EDTA. The sample was loaded on to streptactin-XT superflow resin (IBA-LifeSciences) and the column was washed with 40 volumes of the same buffer. StrepII-tagged protein was eluted by several column volumes of 100 mM Tris-HCl buffer, pH 8, containing150 mM NaCl, 1mM EDTA, and 50 mM biotin (IBA-LifeSciences 2-1016-005). The protein was concentrated via centrifugal concentrator (Amicon 30 kDA MWCO) and exchanged into 20 mM Tris-HCl buffer pH 8.0, with or without 10% glycerol (Sigma-Aldrich 49770) and stored at −80 C for future use. The protein concentration was estimated by BCA assay (Thermo Fisher) versus a BSA standard.

#### Gel Electrophoresis

DvH-FDH2 was separated on a Nupage 4-12% Bis-Tris Gel (Thermo Fisher). The running buffer was 1x MES-SDS. The sample was loaded as 5 μL of 12 μM DvH-FDH2 in 62.5 mM Tris-HCl buffer, containing 1.5% SDS, 10% sucrose, 0.0075% bromphenol blue, pre-incubated at room temp (23 °C) for 30 minutes and then heated 5 minutes at 50 C. The protein was run alongside Precision plus Kaleidoscope prestained standards (Bio-Rad #1610375) for 100 minutes at 100 V (Invitrogen mini gel talk A25977). The gel was fixed in 40% methanol, 10% acetic acid, stained in 30% methanol, 10% acetic acid, and 0.05% Coomassie blue G-250, and destained in 8% acetic acid. Gels were scanned with a gel doc imager (Bio-Rad).

#### Chromogenic Visualization (In-Gel Assay)

DvH-FDH2 was separated on a standard Tris buffered 5% polyacrylamide gel, 2.6% crosslinker gel supplemented with 0.05% triton X-100 (Fisher BP151-100). The running buffer was 25 mM Tris, 192 mM glycine and 0.05% triton X-100 (v/v). Every other lane was loaded with 7.5 μL of 3 μM FDH2 in 20% sucrose, 0.25 M Tris pH 6.9, 0.05% triton, 0.0125% bromphenol blue. Electrophoresis was conducted at 100 V for 209 minutes, 10 mA limit at 4 C. Lanes were cut into strips then assayed in 10 mL of 0.24 mg/mL NBT (Invitrogen N6495) in 20 mM Tris pH 8.0 with or without 10 mM formate (added as 100 μL of 1 M). Strips were incubated with shaking for 15 minutes and then washed with MilliQ water for 3 x 5 min, and then scanned on a Bio-Rad Gel Doc imager.

#### NBT strip assay

3 mm strips of chromatography paper (Whatman, 3MM CHR), cut with a paper cutter, were soaked with a solution consisting of 300 μM NBT (1 mL of 3 mM in water), 30 μM PMS (Sigma P9625) (1 mL of 300 μM in ethanol) in 8 mL Tris-buffered saline pH 7.5, with or without 10 mM formate (Aldrich 798630-500g) (100 μL of 1 M in water). Strips were then spotted with 5 μL of 12 μM enzyme or control solution (buffer only) and monitored for color development. PMS was added to the NBT solutions to 30 μM final from a 30 mM stock in water before soaking the strips. For the NBT only experiment, strips of Whatman chromatography paper (∼8.8 cm^2^) were soaked with 140 μL of 293 μM NBT in 20 mM Tris pH 8.0 with or without 10 mM sodium formate. FDH2 was spotted onto strips as a 5 μL drop of 12 μM enzyme in 20 mM Tris pH 8.0.

#### Metal analysis

Inductively coupled plasma mass spectrometry (ICP-MS; iCAP-RQ, Thermo Fisher Scientific, Bannockburn, IL) was used in the KED mode to assess the metal stoichiometry of DvH-FDH2. 25 μL of protein samples were placed in 15 mL conical tubes followed by the addition of 200 μL of Optima grade HNO_3_. Samples were digested for 20 minutes at room temperature followed by the addition of 9.775 mL of Millipore H_2_O to obtain a final acid matrix of 2% HNO_3_ (v/v). QCS27 high-purity standards (Delta Scientific Laboratory Products) were used as a multi-element standard, as well as tungsten individual standard. The following isotopes were analyzed: ^56,57^Fe, ^63,65^Cu, ^77,78,82^Se, ^95,96,98^Mo, and ^182–184^W. All sample were run with one survey run and three main peak jumping runs.

### Enzymology

#### Benzyl viologen assay

In this assay BV (colorless) is one-electron reduced to BV+ (purple)^130, 131^ by DvH-FDH2. Our workflow is described in Figure S38. FDH2 was assayed in an argon (Airgas AR-300) atmosphere with 2 mM or 20 mM Benzyl viologen (Alfa Aesar H66836) in 50 mM Tris pH 8.0 with 20 mM glucose and supplemented with glucose oxidase (Sigma-G7141-50ku) 1 U/mL, and catalase (Sigma C1345-G) 1 μg/mL, in a stoppered quartz cuvette (Helma 110-10-40) fitted with a Suba seal silicone septum (Sigma Z279730-25ea) and an 8 mm x 3 mm pivoted spin bar (VWR 37119-6183). Assays were performed under argon in a UV-2600i spectrophotometer (Shimadzu Life Sciences) equipped with a T2 Peltier/stirring unit (Quantum; T = 25 °C; stirring speed 650 rpm) and monitored at 557 nm. Measurements were started by the addition of 10 μL formate or buffer blank using a 10 μL Hamilton syringe (under an argon headspace) to a 2.5 mL reaction mix containing pre-equilibrated 1.6 nM FDH2.

#### Phenazine Ethosulfate (PES) / Dichlorophenolindophenol (DCPIP) Assay

Here, PES reacts directly with the enzyme, gets reduced and then transfers electrons to DCPIP non-enzymatically.^78^ Our workflows are described in Figures S39 and S40 We chose PES over PMS because of its long-term stability.^132^ Furthermore, we took the necessary precautions in handling and preparing the reagents.^133^ DvH-FDH2 was assayed in 1 mM PES (Sigma-Aldrich P4544-1G) / 93 μM DCPIP (98% pure; Acros 152870100) in 50 mM Tris pH 8.0. Reactions, 2.5 mL volume, were set up to the desired formate concentration and then started by addition of 10 μL of 400 nM FDH2 (1.6 nM final). Assays were performed using a UV-2600i spectrophotometer (Shimadzu Life Sciences) equipped with T2 Peltier/stirrer accessory (Quantum) and monitored at 600 nm. For aerobic measurements (Figure S39), reactions were performed in open top styrene disposable cuvettes (Brand 75907D) with 6 mm x 9 mm cuvette stirrer (Cowie 001.1609) spun at 400 rpm, and reactions started by addition of 10 μL of 400 nM FDH2 via micropipette. Some aerobic experiments were conducted using a Cary 300 UV-Vis spectrophotometer (Agilent) at a volume of 3.0 mL with a 9 mm x 8 mm spin bar (Sigma Z363545) cuvette stirrer. For the anaerobic conditions (Figure S40), reactions were set up under argon in screw cap quartz cuvettes (Starna 1-SOG-10_GL14-S) sealed with Suba-Seal 13 white rubber septa (Sigma Z167258). The stirrer was an 8 mm x 3 mm pivot bar (VWR 37119-6183), stirring at 650 rpm. Reactions were started by addition of FDH2 via 10 μL syringe (Hamilton 701N 80300) under an argon atmosphere.

#### Data analysis

For classical steady-state kinetics, initial velocities (guided by residual plots) were obtained using ICEKAT.^134^ KinTek Explorer^75, 77^ (version 10.1.6, KinTek Corporation) was used to perform global fitting of enzyme kinetics data to Schemes 1 and S1. This is based on numerical integration of rate equations. Confidence contour analysis was performed to assess whether the parameters were properly constrained by the data.

### Spectroscopy

#### Electronic

FDH2 spectra were collected at 23 °C in 50 mM Tris pH 8.0 using a screw cap 1 cm pathlength quartz cuvette (Starna; 1-SOG_10_GL14s). Under aerobic conditions, the spectrum of air equilibrated enzyme was collected first. Then, 10 mM formate was added, and the spectrum was measured again. The cuvette was then capped with silicone septa (Starna GL14/SI) and 10 μL of 2 mM of dithionite was added under argon before collecting a spectrum. For anaerobic measurements, FDH2 was gassed with argon in the sealed cuvette before addition or formate or dithionite. Reduced spectra were also measured using dithionite as the sole reductant (in the absence of formate).

#### Electron Paramagnetic Resonance (EPR)

All samples were prepared in 20 mM Tris-HCl, pH 7.6 containing 10% glycerol (v/v). Anaerobic samples were first purged with Ar and then transferred to septum-sealed, Ar-flushed EPR tubes and reduced with either 20 mM anaerobic sodium formate or ∼4 mM anaerobic sodium dithionite. Aerobic samples were reduced directly in open EPR tubes with either 20 mM sodium formate or ∼2 mM sodium dithionite. All samples were subsequently frozen in a dry ice/ethanol bath, then transferred to liquid nitrogen for storage. The anaerobic sample reduced with sodium dithionite was incubated for 12 h prior to freezing.

EPR spectra were recorded using a Brüker EMX spectrometer operating WinEPR version 4.33 acquisition software and equipped with a Bruker ER 4119HS high sensitivity X-band cavity and gaussmeter. Temperature was controlled with a Brüker variable temperature unit and a liquid nitrogen or liquid helium cryostat. For purposes of comparison, all spectra were calibrated to a microwave frequency of 9.385 GHz. Detailed instrument settings are included in the figure captions. Simulations were performed using the EasySpin 4.5.5 software package.^135^ Simulations included a “weight” term, which was used to estimate the relative contribution of each component to the composite spectrum.

#### Nuclear Magnetic Resonance

NMR data were recorded on an Agilent DD2 500 MHz spectrometer equipped with a 5 mm quadruple (^1^H,^13^C, ^15^N, ^31^P) PFG Penta Probe, which was maintained at 25 °C. ^13^C data were acquired with 70332 points with a spectral width of 30478 Hz, 242 ppm centered at 110 ppm, with proton-decoupling on throughout the experiment (1 s delay between transients and 1.15s of acquisition time) and the number of transients collected ranged from 64 to 1024. The fids were zero-filled and multiplied with a 3-Hz line-broadening function prior to Fourier-transformation; the final size of the spectrum was 65536 points. Proton data were recorded with 16384 points with a spectral width of 7530 Hz (15 ppm centered at 4.7 ppm) with pre-saturation (2 s) to suppress the water peak; 1 s delay between transients were used. Additional parameters are detailed in Figures S15 – S23. Reference spectra of ^13^C-formate (9.5 – 10.5 mM in 100 mM sodium phosphate buffer, pH 6 or 7.5) and ^13^C-sodium bicarbonate (4.8 mM in 100 mM sodium phosphate buffer, pH 6) were first collected using standard 5 mm thin-walled NMR tubes (Wilmad). 10% D_2_O was used to obtain internal signal lock. Subsequently, 1.3 μM DvH-FDH2 was added to the tube containing ^13^C-formate (pH 7.5), mixed, and spectra were recollected. Upon completion, 2 mM PES was added to the same tube, mixed, and remeasured. Independently, this process was repeated with ^13^C-formate at pH 6. NMR data were processed with MestReNova NMR suite version 14.2.1-27684.

### Quantification of O_2_ Reduction

#### Oximetry

A Clark-type O_2_ electrode (Oxygraph Plus System from Hansatech Instruments, UK) was used to monitor changes to the dissolved O_2_ concentration, which corresponds to 267 μM at 23 °C. O_2_ saturation under these conditions would be equivalent to 1.27 mM. A decrease in O_2_ level would indicate that O_2_ was being consumed during aerobic catalysis. Conversely, O_2_ evolution would be diagnostic of catalase activity. The electrode was calibrated each time before use with air saturated Milli-Q water and dithionite per manufacturer’s instructions. Freshly made reagent stocks and buffer solutions were used throughout. 1 mL reactions were performed at 23 °C in a closed cell using air-saturated 100 mM Tris-HCl, pH 8, containing 1 mM EDTA (Fisher BP120-1). The latter was added to limit adventitious metal ions from mediating O_2_ consumption. After obtaining a stable baseline with the buffer, 10 mM formate was added, and the baseline was allowed to stabilize. The reaction was started by the addition of 50 nM DvH-FDH2. Once the O_2_ consumption plateaued, 2 μM catalase (Sigma C1345-G) was added. Catalase breaks down H_2_O_2_ to water and dioxygen (2H_2_O_2_ → 2H_2_O + O_2_) To test the effect of additives, the order of addition was changed. For example, to test whether DvH-FDH2 had catalase activity, H_2_O_2_ was added to buffer first, followed by the enzyme. Similarly, to assess the effect of SOD on O2 uptake, SOD was the first component to be added. O_2_ consumption rates were calculated as described before.^136^ Initial velocities were determined from the gradients of [O_2_] versus time traces after subtracting O_2_ consumption under the same experimental conditions without FDH2.

### Quantification of Hydrogen Peroxide

#### Amplex Red method

5 mg of Amplex Red (AR; Invitrogen A12222) was dissolved in 0.9725 mL neet DMSO to yield a 20 mM solution. Horseradish peroxidase (HRP; Sigma P8250-5ku) was prepared at a concentration of 10 U/mL (45.5 μg/mL) in sodium phosphate pH 7.4. Prior to assay, a 2x working solution was prepared from 10.6 μL of 20 mM AR, 80 μL of HRP, 1.6 μL of 0.5 M Diethylenetriaminepentaacetic acid (DTPA; TCI D0504) and 3.9 mL of 50 mM sodium phosphate pH 7.4 and kept in the dark. Production of H_2_O_2_ was measured by preparing reaction mixtures in a Costar 3915 black flat bottom 96 well plate. Reactions consisted of 50 mM sodium phosphate pH 7.4, with desired amounts of sodium formate added from a 50 μM stock and initiated by addition of 5 μL of 32 nM DvH-FDH2 in the same buffer to a volume of 50 μL. This approach allowed H_2_O_2_ generation to commence prior to the introduction of AR/HRP mixture. H_2_O_2_ (Sigma-Aldrich H1009-100mL) standard curve was generated in the same buffer to a volume of 50 μL. Detection was initiated by addition of 50 μL of the 2x Amplex Red/HRP working solution and fluorescence was scanned in top read mode at medium sensitivity on a SpectraMax M2 (Molecular Devices) plate reader (excitation 530 nm and emission 590 nm) every four minutes for twelve minutes (23 °C). Independently, we assessed whether outcomes differed when order of addition was varied. To that end, in one set of assays, 5 μL of 32 nM FDH2 was added after AR/HRP. Here, 0.5 mM DTPA was used instead of 0.1 mM.

#### Coumarin boronic acid (CBA) assay

10 mg of coumarin-7-boronic acid (CBA) Cayman Chemicals 14051) was dissolved in 3.33 mL of DMSO. 101 μL of the CBA stock and 1.6 μL 0.5 M DTPA were added to 3.9 mL 50 mM sodium phosphate pH 7.4 to produce a 2x working solution. The remaining steps essential identical to those used in the Amplex Red assay except that the plate was shaken at 400 rpm in an incubator (23 °C) for 15 min prior to CBA addition. Fluorescence detection was initiated by addition of 50 μL of the CBA 2x working solution, and the plate was scanned in fluorescence mode (excitation 332 nm and emission 470 nm).

The following controls were common to both assays. (1) Heat denatured DvH-FDH2, (2) Addition of catalase (100 U/mL), (3) Addition of superoxide dismutase (10 U / mL), (4) Sans FDH2, (5) Fluorogenic substrate excluded, (6) Formate excluded, and (7) Buffer alone.

#### Superoxide Generation / Redox Bifurcation

Native (Sigma C2506) and partially acetylated (Sigma C4186) equine heart cytochrome c were used to assess superoxide production by DvH-FDH2. The integrity of oxidized cytochrome c was validated by establishing the presence of 695 nm transition. The reduction of 30 μM (native) or 60 μM (partially acetylated) cytochrome *c* by DvH-FDH2 was followed at 550 nm in a 1 cm pathlength cell (Shimadzu UV-2600i spectrophotometer). We used two-fold higher concentration of acetylated cytochrome c to offset its slightly weaker reactivity with superoxide.^92^ The reaction mix (total volume 2.5 mL) was stirred (Cowie 001.1609) at 300 rpm (Quantum T2/Peltier unit) and maintained at 25 °C. For aerobic experiments, open top styrene disposable cuvettes (Brand 75907D) were used. For anaerobic measurements under argon, screw cap quartz cuvettes (Starna 1-SOG-10_GL14-S) sealed with Suba-Seal 13 white rubber septa (Sigma Z167258) were used. To 50 mM Tris-HCl buffer, pH 8, containing cytochrome c, enzyme (1.6 nM final) was added first to obtain the background signal. Subsequently, 10 μM formate was added to start the reaction. Upon completion, ∼ 2 mM dithionite was spiked into the mix to estimate the amount of remaining oxidized protein. Controls devoid of formate, enzyme, and cytochrome c were also employed. The effect of superoxide dismutase (10 – 100 U / mL) or catalase (100 – 400 U / mL) was tested independently.

### Structural analysis

Protein alignments were constructed using MUSCLE or MAFFT. Structural alignments were performed using Chimera v1.16. Amino acid sequences of the large (DVU2482) and small (DVU2481) subunits of DvH-FDH2 were input together for running structure predictions using a modified version of AlphaFold2.1.^101^ Because this algorithm does not recognize Sec, a Cys was substituted and that Tat signal peptide (see Fig. 3A) was not included. A dedicated Google Colab notebook (AlphaFold.ipynb-Colaboratory (google.com)), which does not utilize homologous structures for making predictions was used with default settings. We also independently predicted the structures of DVU2482 and DVU2481 using a full version of AlphaFold2.1. The resulting heterodimeric structures were superposed on the DvH-FDH1 counterpart determined via X-ray crystallography (PDB ID: 6SDV^20^) to assess similarities and differences. Difference distance matrices were computed using Chimera v1.16. Structure visualizations and manipulations were done via Pymol.

## AUTHOR INFORMATION

### Corresponding Author

C. S. Raman *

### Author Contributions

C.S.R. conceived the work, designed the experiments, acquired funding, and directed the project. J.E.G. created new DvH strains (identified by the CSR prefix in Table S1), established a novel scaleup strategy and streamlined workflow for generating DvH biomass, optimized DvH transformation protocol, developed a DvH expression platform (cloning, homologous expression, and purification), characterized DvH-FDH2 via electronic spectroscopy, demonstrated in-gel activity, measured enzyme kinetics data, confirmed cytochrome *c* reduction, and validated formate oxidase activity under the supervision of C.S.R. Oximetric measurements were made by C.S.R. and J.E.G. D.N. performed EPR measurements and interpreted the results under the supervision of R.H. DvH deletion strains (with a JW prefix in Table S1) were generated by Q.G., and G.M.Z. measured their growth characteristics under the supervision of J.W. K.H. acquired NMR spectra. C.S.R. analyzed data from enzyme kinetics, oximetry, and NMR, ran the AlphaFold predictions and interpreted the structural differences, and wrote the manuscript. All authors have given approval to the final version of the manuscript.

### Notes

The authors declare the following competing financial interest(s): C.S.R. and J.E.G. are inventors on a patent application related to this work filed by the University of Maryland Baltimore. The authors declare no other competing interests.

### Associated Content

#### Supporting Information

Strains and plasmids (Table S1), Primers (Table S2), Growth curves of JW2111 and JW2121 (Figure S1), NBT strip assay (Figure S2), Sequence alignment of DvH-FD2 and DvH-FDH1 (Figure S3), ICP-MS quantification of metals in DvH-FDH2 (Table S3), Comparison of electronic spectra of DvH-FDH2 and DvH-FDH1 (Figure S4), Simulation of Fe/S centers of formate-reduced DvH-Fdh2 prepared under aerobic conditions (Figure S5), EPR parameters of DvH-FDH2 (Table S4), Steady-state kinetic parameters of SRB-FDHs compiled from the literature (Table S5), Minimal catalytic model used to assess BV kinetics (Scheme S1), Ground truthing published enzyme kinetics data of Dd-FDH3 (Figure S6) and DvH-FDH1 (Figure S7), Steady state kinetics of DvH-FDH2 (Table S6), 2 mM BV reduction by DvH-FDH2 (Figure S8), 20 mm BV reduction by DvH-FDH2 (Figure S9), GO and catalase do not affect the kinetics of BV reduction by DvH-FDH2 (Figure S10), Optimizing DCPIP and PES for aerobic enzyme kinetics (Figure S11), Reduction of PES/DCPIP by DvH-FDH2 in air (Figure S12), Anaerobic reduction of PES/DCPIP by DvH-FDH2 (Figure S13), Effect of azide on the reduction of PES/DCPIP (Figure S14), Effect of nitrate on the reduction of PES/DCPIP (Figure S15), ^13^C NMR spectra of ^13^C-formate + DvH-FDH2 (pH 7.5) (Figure S16), ^13^C NMR spectra of ^13^C-formate + DvH-FDH2 + PES (pH 7.5) (Figure S17), ^13^C NMR spectra of ^13^C-formate + DvH-FDH2 (pH 6) (Figure S18), ^13^C NMR spectra of ^13^C-formate + DvH-FDH2 + PES (pH 6) (Figure S19), ^13^C NMR spectra of ^13^C-NaHCO3 (pH 6) (Figure S20), ^13^C NMR spectra of ^13^C-formate (pH 6) (Figure S21), ^13^C NMR spectra of ^13^C-formate (pH 7.5) (Figure S22), ^1^H NMR spectra of ^13^C-formate (pH 6) (Figure S23), DvH-FDH2 lacks catalase activity (Figure S24), Assessing the effect of variables on amplex red assay (Figure S25), Evaluating how variables impact the CBA assay (Figure S26), O2 uptake by DvH-FDH2 is not impacted by SOD addition (Figure S27), Acetylated cytochrome c is not reduced during aerobic FDH2 catalysis (Figure S28), Initial rates of native cytochrome c reduction is unaffected under aerobic and anaerobic conditions (Figure S29), Spectra of native cytochrome c reduced by FDH2 (Figure S30), Lack of effect of SOD and catalase on the aerobic reduction of equine cytochrome *c* by DvH-FDH2 (Figure S31), Confidence metrics of AlphaFold2.1 prediction (Figure S32), Variation in backbone RMSD (Figures S33 and S34), Residue-residue distance map (Figure S35), Active site of DvH-FDH2 (Figure S36), MOYLS4 medium recipe (Protocol S1), DvH growth in 10 L carboy (Figure S37), BV assay workflow (Figure S38), Aerobic PES/DCPIP assay workflow (Figure S39), and Anaerobic PES/DCPIP assay workflow (Figure S40), and TOC Graphic (OutFoxing Oxygen via Redox Bifurcation)

This information is available free of charge on the ACS Publications website.

## ACKNOWLEDGMENTS

This work is supported by the U.S. Department of Energy (DOE), Office of Science, Basic Energy Sciences (BES) under award # DE-SC-0018047 (C.S.R.). EPR measurements conducted at the University of California Riverside is supported by the DOE, Office of Science, BES under award # DE-SC0010666 (R.H.). C.S.R. is grateful to the late Jane Gibson (née Pinsent) for graciously sharing the early history of FDH and for her constant encouragement to pursue “unusual bioenergetics of bacteria”. C.S.R. thanks David A. Grahame and Luisa Maia for helpful discussions regarding FDH enzymology, Jon Hosler for insights into oximetric data analysis, Clive Bagshaw for stimulating exchanges on fitting by simulation, Luisa Maia, Ana Rita Oliveira, and Inês Pereira for generously sharing source data from their respective publications, Keith MacRenaris (ThermoFisher Scientific) and Bert Woods (Agilent) for help with ICP-MS measurements, Angela Wilks for the use of Cary-300 spectrophotometer, and Norman Brach, Jonathan Soffer and Gilbert Vial (Shimadzu Corporation, Columbia, MD) for timely assistance with the UV-2600i system.

---

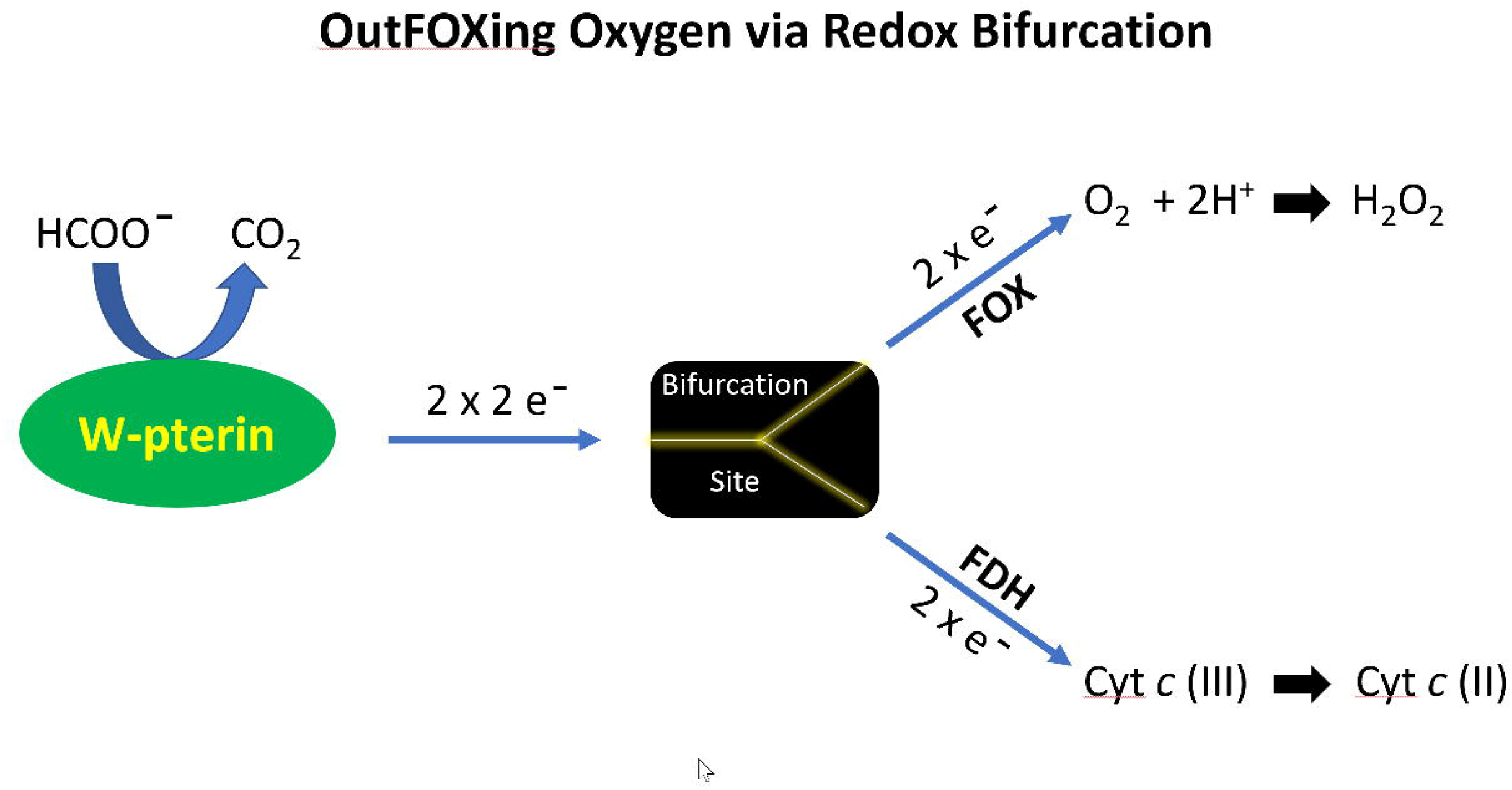

## REFERENCES

1. Pinske, C.; Sawers, R. G., Anaerobic Formate and Hydrogen Metabolism. EcoSal Plus 2016, 7.

2. Hughes, E. R.; Winter, M. G.; Duerkop, B. A.; Spiga, L.; Furtado de Carvalho, T.; Zhu, W.; Gillis, C. C.; Buttner, L.; Smoot, M. P.; Behrendt, C. L.; Cherry, S.; Santos, R. L.; Hooper, L. V.; Winter, S. E., Microbial Respiration and Formate Oxidation as Metabolic Signatures of Inflammation-Associated Dysbiosis. Cell Host Microbe 2017, 21, 208–219.

3. FEEDAP, Scientific opinion on the safety and efficacy of formic acid when used as a technological additive for all animal species. EFSA J. 2014, 12, 3827.

4. Yishai, O.; Lindner, S. N.; Gonzalez de la Cruz, J.; Tenenboim, H.; Bar-Even, A., The formate bio-economy. Curr Opin Chem Biol 2016, 35, 1–9.

5. Du, D.; Lan, R.; Humphreys, J.; Tao, S., Progress in inorganic cathode catalysts for electrochemical conversion of carbon dioxide into formate or formic acid. J. Appl. Electrochem. 2017, 47, 661–678.

6. Roden, E. E.; Jin, Q., Thermodynamics of microbial growth coupled to metabolism of glucose, ethanol, short-chain organic acids, and hydrogen. Appl Environ Microbiol 2011, 77, 1907–9.

7. Windman, T.; Zolotova, N.; Schwandner, F.; Shock, E. L., Formate as an energy source for microbial metabolism in chemosynthetic zones of hydrothermal ecosystems. Astrobiology 2007, 7, 873–90.

8. Crable, B. R.; Plugge, C. M.; McInerney, M. J.; Stams, A. J., Formate formation and formate conversion in biological fuels production. Enzyme Res 2011, 2011, 532536.

9. Sawers, G., The hydrogenases and formate dehydrogenases of Escherichia coli. Antonie Van Leeuwenhoek 1994, 66, 57–88.

10. Maia, L. B.; Moura, I.; Moura, J. J. G., Carbon Dioxide Utilisation - The Formate Route. In Enzymes for Solving Humankind’s Problems: Natural and Artificial Systems in Health, Agriculture, Environment, and Energy, Moura, J. J. G.; Moura, I.; Maia, L. B., Eds. Springer Nature: Switzerland, 2021; pp 29–80.

11. Niks, D.; Hille, R., Molybdenum- and tungsten-containing formate dehydrogenases and formylmethanofuran dehydrogenases: Structure, mechanism, and cofactor insertion. Protein Sci 2019, 28, 111–122.

12. Boyington, J. C.; Gladyshev, V. N.; Khangulov, S. V.; Stadtman, T. C.; Sun, P. D., Crystal structure of formate dehydrogenase H: catalysis involving Mo, molybdopterin, selenocysteine, and an Fe4S4 cluster. Science 1997, 275, 1305–8.

13. Niks, D.; Hille, R., Reductive activation of CO2 by formate dehydrogenases. Methods Enzymol 2018, 613, 277–295.

14. Sebban, C.; Blanchard, L.; Bruschi, M.; Guerlesquin, F., Purification and characterization of the formate dehydrogenase from Desulfovibrio vulgaris Hildenborough. FEMS Microbiol Lett 1995, 133, 143–9.

15. ElAntak, L.; Dolla, A.; Durand, M. C.; Bianco, P.; Guerlesquin, F., Role of the tetrahemic subunit in Desulfovibrio vulgaris hildenborough formate dehydrogenase. Biochemistry 2005, 44, 14828–34.

16. Costa, C.; Teixeira, M.; LeGall, J.; Moura, J. J. G.; Moura, I., Formate dehydrogenase from Desulfovibrio desulfuricans ATCC 27774: isolation and spectroscopic characterization of the active sites (heme, iron-sulfur centers and molybdenum). JBIC Journal of Biological Inorganic Chemistry 1997, 2, 198–208.

17. Maia, L. B.; Fonseca, L.; Moura, I.; Moura, J. J., Reduction of Carbon Dioxide by a Molybdenum-Containing Formate Dehydrogenase: A Kinetic and Mechanistic Study. J Am Chem Soc 2016, 138, 8834–46.

18. Riederer-Henderson, M. A.; Peck, H. D., In Vitro Requirements for Formate Dehydrogenase Activity from Desulfovibrio. Can J Microbiol 1986, 32, 425–429.

19. Almendra, M. J.; Brondino, C. D.; Gavel, O.; Pereira, A. S.; Tavares, P.; Bursakov, S.; Duarte, R.; Caldeira, J.; Moura, J. J.; Moura, I., Purification and characterization of a tungsten-containing formate dehydrogenase from Desulfovibrio gigas. Biochemistry 1999, 38, 16366–72.

20. Oliveira, A. R.; Mota, C.; Mourato, C.; Domingos, R. M.; Santos, M. F. A.; Gesto, D.; Guigliarelli, B.; Santos-Silva, T.; Romão, M. J.; Cardoso Pereira, I. A., Toward the Mechanistic Understanding of Enzymatic CO2 Reduction. ACS Catalysis 2020, 10, 3844–3856.

21. Rivas, M. G.; Gonzalez, P. J.; Brondino, C. D.; Moura, J. J.; Moura, I., EPR characterization of the molybdenum(V) forms of formate dehydrogenase from Desulfovibrio desulfuricans ATCC 27774 upon formate reduction. J Inorg Biochem 2007, 101, 1617–22.

22. Axley, M. J.; Grahame, D. A.; Stadtman, T. C., Escherichia coli formate-hydrogen lyase. Purification and properties of the selenium-dependent formate dehydrogenase component. J Biol Chem 1990, 265, 18213–8.

23. Khangulov, S. V.; Gladyshev, V. N.; Dismukes, G. C.; Stadtman, T. C., Selenium-containing formate dehydrogenase H from Escherichia coli: a molybdopterin enzyme that catalyzes formate oxidation without oxygen transfer. Biochemistry 1998, 37, 3518–28.

24. Friedebold, J.; Bowien, B., Physiological and biochemical characterization of the soluble formate dehydrogenase, a molybdoenzyme from Alcaligenes eutrophus. J Bacteriol 1993, 175, 4719–28.

25. Yu, X.; Niks, D.; Mulchandani, A.; Hille, R., Efficient reduction of CO2 by the molybdenum-containing formate dehydrogenase from Cupriavidus necator (Ralstonia eutropha). J Biol Chem 2017, 292, 16872–16879.

26. Scherer, P. A.; Thauer, R. K., Purification and properties of reduced ferredoxin: CO2 oxidoreductase from Clostridium pasteurianum, a molybdenum iron-sulfur-protein. Eur J Biochem 1978, 85, 125–35.

27. Kroger, A.; Winkler, E.; Innerhofer, A.; Hackenberg, H.; Schagger, H., The formate dehydrogenase involved in electron transport from formate to fumarate in Vibrio succinogenes. Eur J Biochem 1979, 94, 465–75.

28. Hartmann, T.; Leimkuhler, S., The oxygen-tolerant and NAD+-dependent formate dehydrogenase from Rhodobacter capsulatus is able to catalyze the reduction of CO2 to formate. FEBS J 2013, 280, 6083–96.

29. Stickland, L. H., The bacterial decomposition of formic acid. Biochem J 1929, 23, 1187–98.

30. Stephenson, M.; Stickland, L. H., Hydrogenlyases: Bacterial enzymes liberating molecular hydrogen. Biochem J 1932, 26, 712–24.

31. Gale, E. F., Formic dehydrogenase of Bacterium coli: its inactivation by oxygen and its protection in the bacterial cell. Biochem J 1939, 33, 1012–27.

32. Pinsent, J., The need for selenite and molybdate in the formation of formic dehydrogenase by members of the coli-aerogenes group of bacteria. Biochem J 1954, 57, 10–6.

33. Itagaki, E.; Fujita, T.; Sato, R., Solubilization and properties of formate dehydrogenase and cytochrome b1 from Escherichia coli. J. Biochem. 1962, 52, 131–141.

34. Linnane, A. W.; Wrigley, C. W., Fragmentation of the Electron Transport Chain of Escherichia Coli. Preparation of a Soluble Formate Dehydrogenase-Cytochrome B1 Complex. Biochim Biophys Acta 1963, 77, 408-18.

35. Gray, C. T.; Gest, H., Biological Formation of Molecular Hydrogen. Science 1965, 148, 186–92.

36. Ruiz-Herrera, J.; Alvarez, A., A physiological study of formate dehydrogenase, formate oxidase, and hydrogenlyase from Escherichia coli K-12. Antonie Van Leeuwenhoek 1972, 38, 479–491.

37. Ljungdahl, L. G., Formate Dehydrogenases: Role of Molybdenum, Tungsten, and Selenium. In Molybdenum and Molybdenum-Containing Enzymes, Coughlan, M. P., Ed. Pergamon: New York, 1980; pp 463–486.

38. Pichinoty, F., Recherche des activités formiate-oxydase, hydrogène-lyase, hydrogénase et formiate-déshydrogénase chez quelques enterobacteriaceae. Ann Inst Pasteur (Paris) 1969, 117, 3–15.

39. Stewart, V., Nitrate respiration in relation to facultative metabolism in enterobacteria. Microbiol Rev 1988, 52, 190–232.

40. Finney, A. J.; Sargent, F., Formate hydrogenlyase: A group 4 [NiFe]-hydrogenase in tandem with a formate dehydrogenase. Adv Microb Physiol 2019, 74, 465–486.

41. Sawers, G.; Heider, J.; Zehelein, E.; Bock, A., Expression and operon structure of the sel genes of Escherichia coli and identification of a third selenium-containing formate dehydrogenase isoenzyme. J Bacteriol 1991, 173, 4983–93.

42. Soboh, B.; Pinske, C.; Kuhns, M.; Waclawek, M.; Ihling, C.; Trchounian, K.; Trchounian, A.; Sinz, A.; Sawers, G., The respiratory molybdo-selenoprotein formate dehydrogenases of Escherichia coli have hydrogen: benzyl viologen oxidoreductase activity. BMC Microbiol 2011, 11, 173.

43. Abaibou, H.; Pommier, J.; Benoit, S.; Giordano, G.; Mandrand-Berthelot, M. A., Expression and characterization of the Escherichia coli fdo locus and a possible physiological role for aerobic formate dehydrogenase. J Bacteriol 1995, 177, 7141–9.

44. Benoit, S.; Abaibou, H.; Mandrand-Berthelot, M. A., Topological analysis of the aerobic membrane-bound formate dehydrogenase of Escherichia coli. J Bacteriol 1998, 180, 6625–34.

45. Unden, G.; Steinmetz, P. A.; Degreif-Dunnwald, P., The Aerobic and Anaerobic Respiratory Chain of Escherichia coli and Salmonella enterica: Enzymes and Energetics. EcoSal Plus 2014, 6.

46. Richardson, D.; Sawers, G., Structural biology. PMF through the redox loop. Science 2002, 295, 1842–3.

47. Yagi, T., Formate: cytochrome oxidoreductase of Desulfovibrio vulgaris. J Biochem 1969, 66, 473–8.

48. Yagi, T., Monoheme cytochromes. Methods Enzymol 1994, 243, 104–18.

49. Keller, K. L.; Wall, J. D.; Chhabra, S., Methods for engineering sulfate reducing bacteria of the genus Desulfovibrio. Methods Enzymol 2011, 497, 503–17.

50. Rabus, R.; Venceslau, S. S.; Wohlbrand, L.; Voordouw, G.; Wall, J. D.; Pereira, I. A., A Post-Genomic View of the Ecophysiology, Catabolism and Biotechnological Relevance of Sulphate-Reducing Prokaryotes. Adv Microb Physiol 2015, 66, 55–321.

51. Price, M. N.; Wetmore, K. M.; Waters, R. J.; Callaghan, M.; Ray, J.; Liu, H.; Kuehl, J. V.; Melnyk, R. A.; Lamson, J. S.; Suh, Y.; Carlson, H. K.; Esquivel, Z.; Sadeeshkumar, H.; Chakraborty, R.; Zane, G. M.; Rubin, B. E.; Wall, J. D.; Visel, A.; Bristow, J.; Blow, M. J.; Arkin, A. P.; Deutschbauer, A. M., Mutant phenotypes for thousands of bacterial genes of unknown function. Nature 2018, 557, 503–509.

52. Verhagen, M. F.; Wolbert, R. B.; Hagen, W. R., Cytochrome c553 from Desulfovibrio vulgaris (Hildenborough). Electrochemical properties and electron transfer with hydrogenase. Eur J Biochem 1994, 221, 821–9.

53. Lefevre, C. T.; Howse, P. A.; Schmidt, M. L.; Sabaty, M.; Menguy, N.; Luther, G. W., 3rd; Bazylinski, D. A., Growth of magnetotactic sulfate-reducing bacteria in oxygen concentration gradient medium. Environ Microbiol Rep 2016, 8, 1003–1015.

54. Schoeffler, M.; Gaudin, A. L.; Ramel, F.; Valette, O.; Denis, Y.; Hania, W. B.; Hirschler-Rea, A.; Dolla, A., Growth of an anaerobic sulfate-reducing bacterium sustained by oxygen respiratory energy conservation after O2-driven experimental evolution. Environ Microbiol 2019, 21, 360–373.

55. Muyzer, G.; Stams, A. J., The ecology and biotechnology of sulphate-reducing bacteria. Nat Rev Microbiol 2008, 6, 441–54.

56. da Silva, S. M.; Pimentel, C.; Valente, F. M.; Rodrigues-Pousada, C.; Pereira, I. A., Tungsten and molybdenum regulation of formate dehydrogenase expression in Desulfovibrio vulgaris Hildenborough. J Bacteriol 2011, 193, 2909–16.

57. Heidelberg, J. F.; Seshadri, R.; Haveman, S. A.; Hemme, C. L.; Paulsen, I. T.; Kolonay, J. F.; Eisen, J. A.; Ward, N.; Methe, B.; Brinkac, L. M.; Daugherty, S. C.; Deboy, R. T.; Dodson, R. J.; Durkin, A. S.; Madupu, R.; Nelson, W. C.; Sullivan, S. A.; Fouts, D.; Haft, D. H.; Selengut, J.; Peterson, J. D.; Davidsen, T. M.; Zafar, N.; Zhou, L.; Radune, D.; Dimitrov, G.; Hance, M.; Tran, K.; Khouri, H.; Gill, J.; Utterback, T. R.; Feldblyum, T. V.; Wall, J. D.; Voordouw, G.; Fraser, C. M., The genome sequence of the anaerobic, sulfate-reducing bacterium Desulfovibrio vulgaris Hildenborough. Nat Biotechnol 2004, 22, 554–9.

58. Lopez, S.; Prieto, M.; Dijkstra, J.; Dhanoa, M. S.; France, J., Statistical evaluation of mathematical models for microbial growth. Int J Food Microbiol 2004, 96, 289–300.

59. de Bok, F. A.; Roze, E. H.; Stams, A. J., Hydrogenases and formate dehydrogenases of Syntrophobacter fumaroxidans. Antonie Van Leeuwenhoek 2002, 81, 283–91.

60. Hartwig, S.; Pinske, C.; Sawers, R. G., Chromogenic assessment of the three molybdo-selenoprotein formate dehydrogenases in Escherichia coli. Biochem Biophys Rep 2015, 1, 62–67.

61. Enoch, H. G.; Lester, R. L., The purification and properties of formate dehydrogenase and nitrate reductase from Escherichia coli. J Biol Chem 1975, 250, 6693–705.

62. Pinske, C.; Jaroschinsky, M.; Sargent, F.; Sawers, G., Zymographic differentiation of [NiFe]-hydrogenases 1, 2 and 3 of Escherichia coli K-12. BMC Microbiol 2012, 12, 134.

63. Schafer, C.; Friedrich, B.; Lenz, O., Novel, oxygen-insensitive group 5 [NiFe]-hydrogenase in Ralstonia eutropha. Appl Environ Microbiol 2013, 79, 5137–45.

64. Hagen, W. R., Tungsten-Containing Enzymes. In Molybdenum and Tungsten Enzymes: Bioichemistry, Hille, R., Schulzke, C., Kirk, M.L., Ed. The Royal Society of Chemistry: Cambridge, 2017; pp 313–342.

65. Gladyshev, V. N.; Boyington, J. C.; Khangulov, S. V.; Grahame, D. A.; Stadtman, T. C.; Sun, P. D., Characterization of crystalline formate dehydrogenase H from Escherichia coli. Stabilization, EPR spectroscopy, and preliminary crystallographic analysis. J Biol Chem 1996, 271, 8095–100.

66. Orme-Johnson, W. H., Iron-sulfur proteins: structure and function. Annu Rev Biochem 1973, 42, 159–204.

67. Sweeney, W. V.; Rabinowitz, J. C., Proteins containing 4Fe-4S clusters: an overview. Annu Rev Biochem 1980, 49, 139–61.

68. Raaijmakers, H.; Teixeira, S.; Dias, J. M.; Almendra, M. J.; Brondino, C. D.; Moura, I.; Moura, J. J.; Romao, M. J., Tungsten-containing formate dehydrogenase from Desulfovibrio gigas: metal identification and preliminary structural data by multi-wavelength crystallography. J Biol Inorg Chem 2001, 6, 398–404.

69. de Bok, F. A.; Hagedoorn, P. L.; Silva, P. J.; Hagen, W. R.; Schiltz, E.; Fritsche, K.; Stams, A. J., Two W-containing formate dehydrogenases (CO2-reductases) involved in syntrophic propionate oxidation by Syntrophobacter fumaroxidans. Eur J Biochem 2003, 270, 2476–85.

70. Koehler, B. P.; Mukund, S.; Conover, R. C.; Dhawan, I. K.; Roy, R.; Adams, M. W.; Johnson, M. K., Spectroscopic characterization of the tungsten and iron centers in aldehyde ferredoxin oxidoreductases from two hyperthermophilic archaea. J Am Chem Soc 1996, 118, 12391–12405.

71. Bassegoda, A.; Madden, C.; Wakerley, D. W.; Reisner, E.; Hirst, J., Reversible interconversion of CO2 and formate by a molybdenum-containing formate dehydrogenase. J Am Chem Soc 2014, 136, 15473–6.

72. Silveira, C. M.; Besson, S.; Moura, I.; Moura, J. J.; Almeida, M. G., Measuring the cytochrome C nitrite reductase activity-practical considerations on the enzyme assays. Bioinorg Chem Appl 2010.

73. Duggleby, R. G.; Clarke, R. B., Experimental designs for estimating the parameters of the Michaelis-Menten equation from progress curves of enzyme-catalyzed reactions. Biochim Biophys Acta 1991, 1080, 231–6.

74. Schweins, T.; Geyer, M.; Scheffzek, K.; Warshel, A.; Kalbitzer, H. R.; Wittinghofer, A., Substrate-assisted catalysis as a mechanism for GTP hydrolysis of p21ras and other GTP-binding proteins. Nat Struct Biol 1995, 2, 36–44.

75. Johnson, K. A., Fitting enzyme kinetic data with KinTek Global Kinetic Explorer. Methods Enzymol 2009, 467, 601–626.

76. Stroberg, W.; Schnell, S., On the estimation errors of KM and V from time-course experiments using the Michaelis-Menten equation. Biophys Chem 2016, 219, 17–27.

77. Johnson, K. A., New standards for collecting and fitting steady state kinetic data. Beilstein J Org Chem 2019, 15, 16–29.

78. Jahn, B.; Jonasson, N. S. W.; Hu, H.; Singer, H.; Pol, A.; Good, N. M.; den Camp, H.; Martinez-Gomez, N. C.; Daumann, L. J., Understanding the chemistry of the artificial electron acceptors PES, PMS, DCPIP and Wurster’s Blue in methanol dehydrogenase assays. J Biol Inorg Chem 2020, 25, 199–212.

79. Prince, R. C.; Linkletter, S. J.; Dutton, P. L., The thermodynamic properties of some commonly used oxidation-reduction mediators, inhibitors and dyes, as determined by polarography. Biochim Biophys Acta 1981, 635, 132–48.

80. Peck, H. D., Jr.; Gest, H., Formic dehydrogenase and the hydrogenlyase enzyme complex in coli-aerogenes bacteria. J Bacteriol 1957, 73, 706–21.

81. Weissgerber, T. L.; Winham, S. J.; Heinzen, E. P.; Milin-Lazovic, J. S.; Garcia-Valencia, O.; Bukumiric, Z.; Savic, M. D.; Garovic, V. D.; Milic, N. M., Reveal, Don’t Conceal: Transforming Data Visualization to Improve Transparency. Circulation 2019, 140, 1506–1518.

82. Xiang, D.; Magana, D.; Dyer, R. B., CO2 reduction catalyzed by mercaptopteridine on glassy carbon. J Am Chem Soc 2014, 136, 14007–10.

83. Merritt, M. E.; Harrison, C.; Storey, C.; Jeffrey, F. M.; Sherry, A. D.; Malloy, C. R., Hyperpolarized 13C allows a direct measure of flux through a single enzyme-catalyzed step by NMR. Proc Natl Acad Sci U S A 2007, 104, 19773–7.

84. Niekus, H. G.; Stouthamer, A. H., Formate oxidase in glutaraldehyde-treated Campylobacter sputorum subspecies bubulus. FEMS Microbiol Lett 1981, 11, 83–87.

85. Goodhew, C. F.; elKurdi, A. B.; Pettigrew, G. W., The microaerophilic respiration of Campylobacter mucosalis. Biochim Biophys Acta 1988, 933, 114–23.

86. Miner, K. D.; Mukherjee, A.; Gao, Y. G.; Null, E. L.; Petrik, I. D.; Zhao, X.; Yeung, N.; Robinson, H.; Lu, Y., A designed functional metalloenzyme that reduces O2 to H2O with over one thousand turnovers. Angew Chem Int Ed Engl 2012, 51, 5589–92.

87. Kakeshpour, T.; Bax, A., NMR characterization of H2O2 hydrogen exchange. J Magn Reson 2021, 333, 107092.

88. Kalyanaraman, B.; Darley-Usmar, V.; Davies, K. J.; Dennery, P. A.; Forman, H. J.; Grisham, M. B.; Mann, G. E.; Moore, K.; Roberts, L. J., 2nd; Ischiropoulos, H., Measuring reactive oxygen and nitrogen species with fluorescent probes: challenges and limitations. Free Radic Biol Med 2012, 52, 1–6.

89. Wulff, P.; Day, C. C.; Sargent, F.; Armstrong, F. A., How oxygen reacts with oxygen-tolerant respiratory [NiFe]-hydrogenases. Proc Natl Acad Sci U S A 2014, 111, 6606–11.

90. Zielonka, J.; Sikora, A.; Joseph, J.; Kalyanaraman, B., Peroxynitrite is the major species formed from different flux ratios of co-generated nitric oxide and superoxide: direct reaction with boronate-based fluorescent probe. J Biol Chem 2010, 285, 14210–6.

91. Tarpey, M. M.; Fridovich, I., Methods of detection of vascular reactive species: nitric oxide, superoxide, hydrogen peroxide, and peroxynitrite. Circ Res 2001, 89, 224–36.

92. Azzi, A.; Montecucco, C.; Richter, C., The use of acetylated ferricytochrome c for the detection of superoxide radicals produced in biological membranes. Biochem Biophys Res Commun 1975, 65, 597–603.

93. Lu, Y.; Koo, J., O2 sensitivity and H2 production activity of hydrogenases-A review. Biotechnol Bioeng 2019, 116, 3124–3135.

94. Shaw, R. W.; Rife, J. E.; O’Leary, M. H.; Beinert, H., Oxidation of reduced cytochrome c oxidase with 18O2. A search for mu-oxo-bridged metal species in the oxidized enzyme. J Biol Chem 1981, 256, 1105–7.

95. Lauterbach, L.; Lenz, O., Catalytic production of hydrogen peroxide and water by oxygen-tolerant [NiFe]-hydrogenase during H2 cycling in the presence of O2. J Am Chem Soc 2013, 135, 17897–905.

96. Peters, J. W.; Beratan, D. N.; Bothner, B.; Dyer, R. B.; Harwood, C. S.; Heiden, Z. M.; Hille, R.; Jones, A. K.; King, P. W.; Lu, Y.; Lubner, C. E.; Minteer, S. D.; Mulder, D. W.; Raugei, S.; Schut, G. J.; Seefeldt, L. C.; Tokmina-Lukaszewska, M.; Zadvornyy, O. A.; Zhang, P.; Adams, M. W., A new era for electron bifurcation. Curr Opin Chem Biol 2018, 47, 32–38.

97. Hoke, K. R.; Cobb, N.; Armstrong, F. A.; Hille, R., Electrochemical studies of arsenite oxidase: an unusual example of a highly cooperative two-electron molybdenum center. Biochemistry 2004, 43, 1667–74.

98. Yuly, J. L.; Zhang, P.; Ru, X.; Terai, K.; Singh, N.; Beratan, D. N., Efficient and reversible electron bifurcation with either normal or inverted potentials at the bifurcating cofactor. Chem 2021, 7, 1–17.

99. Nitschke, W.; Russell, M. J., Redox bifurcations: mechanisms and importance to life now, and at its origin: a widespread means of energy conversion in biology unfolds. Bioessays 2012, 34, 106–9.

100. Chowdhury, N. P.; Kahnt, J.; Buckel, W., Reduction of ferredoxin or oxygen by flavin-based electron bifurcation in Megasphaera elsdenii. FEBS J 2015, 282, 3149–60.

101. Jumper, J.; Evans, R.; Pritzel, A.; Green, T.; Figurnov, M.; Ronneberger, O.; Tunyasuvunakool, K.; Bates, R.; Zidek, A.; Potapenko, A.; Bridgland, A.; Meyer, C.; Kohl, S. A. A.; Ballard, A. J.; Cowie, A.; Romera-Paredes, B.; Nikolov, S.; Jain, R.; Adler, J.; Back, T.; Petersen, S.; Reiman, D.; Clancy, E.; Zielinski, M.; Steinegger, M.; Pacholska, M.; Berghammer, T.; Bodenstein, S.; Silver, D.; Vinyals, O.; Senior, A. W.; Kavukcuoglu, K.; Kohli, P.; Hassabis, D., Highly accurate protein structure prediction with AlphaFold. Nature 2021, 596, 583–589.

102. Holm, L., Using Dali for Protein Structure Comparison. Methods Mol Biol 2020, 2112, 29–42.

103. Reich, H. J.; Hondal, R. J., Why Nature Chose Selenium. ACS Chem Biol 2016, 11, 821–41.

104. Evans, R. M.; Krahn, N.; Murphy, B. J.; Lee, H.; Armstrong, F. A.; Soll, D., Selective cysteine-to-selenocysteine changes in a [NiFe]-hydrogenase confirm a special position for catalysis and oxygen tolerance. Proc Natl Acad Sci U S A 2021, 118.

105. Nicolet, Y.; Fontecilla-Camps, J. C., Iron-sulfur clusters and molecular oxygen:function, adaptation, degradation, and repair. In Iron-Sulfur Clusters in Chemistry and Biology, Rouault, T. A., Ed. 2014; pp 359–385.

106. Lu, Z.; Imlay, J. A., When anaerobes encounter oxygen: mechanisms of oxygen toxicity, tolerance and defence. Nat Rev Microbiol 2021.

107. Sen, A.; Imlay, J. A., How Microbes Defend Themselves From Incoming Hydrogen Peroxide. Front Immunol 2021, 12, 667343.

108. Niekus, H. G.; Van Doorn, E.; De Vries, W.; Stouthamer, A. H., Aerobic growth of Campylobacter sputorum subspecies bubulus with formate. J. Gen. Microbiol. 1980, 118, 419–428.

109. Ohta, H.; Gottschal, J. C., Formate oxidation by Wolinella recta ATCC 33238 with oxygen as electron acceptor. FEMS Microbiol Lett 1988, 50, 163–188.

110. Hoffman, P. S.; Goodman, T. G., Respiratory physiology and energy conservation efficiency of Campylobacter jejuni. J Bacteriol 1982, 150, 319–26.

111. Taylor, A. J.; Kelly, D. J., The function, biogenesis and regulation of the electron transport chains in Campylobacter jejuni: New insights into the bioenergetics of a major food-borne pathogen. Adv Microb Physiol 2019, 74, 239–329.

112. Khademian, M.; Imlay, J. A., Escherichia coli cytochrome c peroxidase is a respiratory oxidase that enables the use of hydrogen peroxide as a terminal electron acceptor. Proc Natl Acad Sci U S A 2017, 114, E6922–E6931.

113. Nishikawa, K.; Ogata, H.; Higuchi, Y., Structural basis of the function of [NiFe]-hydrogenases. Chem Lett 2020, 49, 164–173.

114. Noor, N. D. M.; Matsuura, H.; Nishikawa, K.; Tai, H.; Hirota, S.; Kim, J.; Kang, J.; Tateno, M.; Yoon, K. S.; Ogo, S.; Kubota, S.; Shomura, Y.; Higuchi, Y., Redox-dependent conformational changes of a proximal [4Fe-4S] cluster in Hyb-type [NiFe]-hydrogenase to protect the active site from O2. Chem Commun (Camb) 2018, 54, 12385–12388.

115. Sakai, K.; Kitazumi, Y.; Shirai, O.; Takagi, K.; Kano, K., High-Power Formate/Dioxygen Biofuel Cell Based on Mediated Electron Transfer Type Bioelectrocatalysis. ACS Catalysis 2017, 7, 5668–5673.

116. Meneghello, M.; Leger, C.; Fourmond, V., Electrochemical Studies of CO2-Reducing Metalloenzymes. Chemistry 2021, 27, 17542–17553.

117. Ruth, J. C.; Spormann, A. M., Enzyme electrochemistry for industrial applications - A perspective on future area of focus. ACS Catal 2021, 11, 5951–5967.

118. Cracknell, J. A.; Vincent, K. A.; Ludwig, M.; Lenz, O.; Friedrich, B.; Armstrong, F. A., Enzymatic oxidation of H2 in atmospheric O2: the electrochemistry of energy generation from trace H2 by aerobic microorganisms. J Am Chem Soc 2008, 130, 424–5.

119. Vincent, K. A.; Cracknell, J. A.; Lenz, O.; Zebger, I.; Friedrich, B.; Armstrong, F. A., Electrocatalytic hydrogen oxidation by an enzyme at high carbon monoxide or oxygen levels. Proc Natl Acad Sci U S A 2005, 102, 16951–4.

120. Vincent, K. A.; Cracknell, J. A.; Clark, J. R.; Ludwig, M.; Lenz, O.; Friedrich, B.; Armstrong, F. A., Electricity from low-level H2 in still air--an ultimate test for an oxygen tolerant hydrogenase. Chem Commun (Camb) 2006, 5033–5.

121. Mulder, D. W.; Peters, J. W.; Raugei, S., Catalytic bias in oxidation-reduction catalysis. Chem Commun (Camb) 2021, 57, 713–720.

122. Fourmond, V.; Leger, C.; Plumere, N., Reversible catalysis. Nat Rev Chem 2021, 5, 348–360.

123. Shafaat, H. S.; Yang, J. Y., Uniting biological and chemcal strategies for selective CO2 reduction. Nat Catal 2021, 4, 928–939.

124. Magalon, A.; Alberge, F., Distribution and dynamics of OXPHOS complexes in the bacterial cytoplasmic membrane. Biochim Biophys Acta 2016, 1857, 198–213.

125. Hillesland, K. L.; Lim, S.; Flowers, J. J.; Turkarslan, S.; Pinel, N.; Zane, G. M.; Elliott, N.; Qin, Y.; Wu, L.; Baliga, N. S.; Zhou, J.; Wall, J. D.; Stahl, D. A., Erosion of functional independence early in the evolution of a microbial mutualism. Proc Natl Acad Sci U S A 2014, 111, 14822–7.

126. Keller, K. L.; Bender, K. S.; Wall, J. D., Development of a markerless genetic exchange system for Desulfovibrio vulgaris Hildenborough and its use in generating a strain with increased transformation efficiency. Appl Environ Microbiol 2009, 75, 7682–91.

127. Li, M. Z.; Elledge, S. J., Harnessing homologous recombination in vitro to generate recombinant DNA via SLIC. Nat Methods 2007, 4, 251–6.

128. Zane, G. M.; Yen, H. C.; Wall, J. D., Effect of the deletion of qmoABC and the promoter-distal gene encoding a hypothetical protein on sulfate reduction in Desulfovibrio vulgaris Hildenborough. Appl Environ Microbiol 2010, 76, 5500–9.

129. Studier, F. W., Protein production by auto-induction in high density shaking cultures. Protein Expr Purif 2005, 41, 207–34.

130. Jones, R. W.; Garland, P. B., Sites and specificity of the reaction of bipyridylium compounds with anaerobic respiratory enzymes of Escherichia coli. Effects of permeability barriers imposed by the cytoplasmic membrane. Biochem J 1977, 164, 199–211.

131. Axley, M. J.; Grahame, D. A., Kinetics for formate dehydrogenase of Escherichia coli formate-hydrogenlyase. J Biol Chem 1991, 266, 13731–6.

132. Ghosh, R.; Quayle, J. R., Phenazine ethosulfate as a preferred electron acceptor to phenazine methosulfate in dye-linked enzyme assays. Anal Biochem 1979, 99, 112–7.

133. Jahn, B.; Pol, A.; Lumpe, H.; Barends, T. R. M.; Dietl, A.; Hogendoorn, C.; Op den Camp, H. J. M.; Daumann, L. J., Similar but Not the Same: First Kinetic and Structural Analyses of a Methanol Dehydrogenase Containing a Europium Ion in the Active Site. Chembiochem 2018.

134. Olp, M. D.; Kalous, K. S.; Smith, B. C., ICEKAT: an interactive online tool for calculating initial rates from continuous enzyme kinetic traces. BMC Bioinformatics 2020, 21, 186.

135. Stoll, S.; Schweiger, A., EasySpin, a comprehensive software package for spectral simulation and analysis in EPR. J Magn Reson 2006, 178, 42–55.

136. Brautigan, D. L.; Ferguson-Miller, S.; Margoliash, E., Mitochondrial cytochrome c: preparation and activity of native and chemically modified cytochromes c. Methods Enzymol 1978, 53, 128–64.

