## Supplementary Figures and Tables for "How a Formate Dehydrogenase Responds to Oxygen: Unexpected O_2_ Insensitivity of an Enzyme Harboring Tungstopterin, Selenocysteine, and [4Fe-4S] Clusters"

### Supporting Information

Joel E. Graham,‡ Dimitri Niks,¶ Grant M. Zane,† Qin Gui,† Kellie Hom,‡ Russ Hille,¶ Judy D. Wall,† and C. S. Raman‡*

‡Department of Pharmaceutical Sciences, University of Maryland, Baltimore, MD 21201, USA; ¶Department of Biochemistry, University of California, Riverside, CA 92521; †Department of Biochemistry, University of Missouri, Columbia, MO 65211, USA.

Table S1: Strains and plasmids used in this study.[§]

| Strain or plasmid | Genotype and relevant features | Source |
|---|---|---|
| *Desulfovibrio vulgaris* strains | | |
| *Desulfovibrio vulgaris* Hildenborough | Wild-type strain, ATCC 29579 | ATCC |
| JW710 | *Desulfovibrio vulgaris* Hildenborough Δ*upp* 5FU$^r$ | [1] |
| JW2103 | JW710 Δ*fdh2* | This study |
| JW2109 | JW710 Δ*fdh3 aph(3')-IIa:upp* G418$^r$ 5FU$^s$ | This study |
| JW2111 | JW710 Δ*fdh3* | This study |
| JW2115 | JW710 Δ*fdh1* | This study |
| JW2117 | JW710 Δ*fdh2* Δ*fdh3* | This study |
| JW2120 | JW710 Δ*fdh3* Δ*fdh1 aph(3')-IIa:upp* G418$^r$ 5FU$^s$ | This study |
| JW2121 | JW710 Δ*fdh3* Δ*fdh1* | This study |
| JW2123 | JW710 Δ*fdh1* Δ*fdh2* | This study |
| JW2126 | JW710 Δ*fdh3* Δ*fdh1* Δ*fdh2 aph(3')-IIa:upp* G418$^r$ 5FU$^s$ | This study |
| JW2127 | JW710 Δ*fdh3* Δ*fdh1* Δ*fdh2* | This study |
| CSR21210 | JW2121 transformed with pJEG127 | This study |
| CSR21271 | JW2127 transformed with pJEG132 | This study |
| *Escherichia coli* strains | | |
| α-select | *deoR endA1 recA1 relA1 gyrA96 hsdr17*($r_k^-$ $m_k^+$) *supE44 thi-1* Δ(*lacZYA-argFU169*) ϕ80δ*lacZΔM15* F- λ- | Bioline |
| **Plasmids** | | |

| | | |
|---|---|---|
| pCR8/GW/TOPO | plasmid used to amplify pUC-Sp$^r$ fragment, Sp$^r$ | Life Technologies |
| pMO746 | plasmid containing *aph(3')-II:upp* 2-gene operon, Ap$^r$, Km$^r$ | [2] |
| pMO2100 | plasmid containing upstream and downstream regions of *fdh-2* on either side of *aph(3')-II:upp* (used to construct marker-exchange deletion), Sp$^r$, Km$^r$ | This study |
| pMO2102 | plasmid containing upstream and downstream regions of *fdh-2* (used to construct marker-less deletion), Sp$^r$ | This study |
| pMO2108 | plasmid containing upstream and downstream regions of *fdh-3* on either side of *aph(3')-II:upp* (used to construct marker-exchange deletion), Sp$^r$, Km$^r$ | This study |
| pMO2110 | plasmid containing upstream and downstream regions of *fdh-3* (used to construct marker-less deletion), Sp$^r$ | This study |
| pMO2112 | plasmid containing upstream and downstream regions of *fdh-1* on either side of *aph(3')-II:upp* (used to construct marker-exchange deletion), Sp$^r$, Km$^r$ | This study |
| pMO2114 | plasmid containing upstream and downstream regions of *fdh-1* (used to construct marker-less deletion), Sp$^r$ | This study |
| pMO9075 | Plasmid used for genetic complementation and/or gene expression in *Desulfovibrio* strains. It contains Km$^r$ gene-*aph*(*3'*)-II promoter, pGB1, Sp$^r$, and RBS. | [1,2] |
| pJEG127 | pMO9075 containing the DVU2482-2481 insert | This study |
| pJEG132 | pMO9075 containing the DVU2482strII-2481 insert | This study |

[§]Δ*fdh1*, Δ*fdh2*, and Δ*fdh3* represent deletion of DVU0586-0588, DVU2481-2485, and DVU2809-2812 operons in DvH, respectively.

**Table S2: Primers used in this study.**

| Primer name | Primer sequence$^{\%}$ | Application |
|---|---|---|
| SpecRpUC-F | CCAGCCAGGACAGAAATGCCTCG | Amplification of pUC-Sp$^r$ fragement |
| SpecRpUC-R | ATGTGAGCAAAAGGCCAGCAAAAGGC | Amplification of pUC-Sp$^r$ fragement |
| Kan gene Prom Nterm | CCGGAATTGCCAGCTGGGGCGC | Amplification of *aph(3')-IIa* and *upp* fragement |
| *upp* gene Cterm | CTTACTTGGTGCCGAATATCTTGTCGC | Amplification of *aph(3')-IIa* and *upp* fragement |
| DVU2481-5-upF | GCCTTTTGCTGGCCTTTTGCTCACAT<br>CACTCTTGCGCGAGGAAAGC | Amplification of upstream region of DVU2481-5 |
| DVU2481-5-upR | GCGACAAGATATTCGGCACCAAGTAAG<br>GGGAAGGCATTAACCGATACTTG | Amplification of upstream region of DVU2481-5, specific for marker-exchange plasmid |
| DVU2481-5-dnF | GCGCCCCAGCTGGCAATTCCGG<br>CCGACTGGATACGCAACACC | Amplification of downstream region of DVU2481-5, specific for marker-exchange plasmid |
| DVU2481-5-dnR | CGAGGCATTTCTGTCCTGGCTGG<br>CCTGTTCGGACTCTCGATGTTC | Amplification of downstream region of DVU2481-5 |
| DVU2809-12-upF | GCCTTTTGCTGGCCTTTTGCTCACAT<br>CAGAACCTCATCGCCATGC | Amplification of upstream region of DVU2809-12 |
| DVU2809-12-upR | GCGACAAGATATTCGGCACCAAGTAAG<br>TCCTCTCCTTGTTGATGCCCTG | Amplification of upstream region of DVU2809-12, specific for marker-exchange plasmid |
| DVU2809-12-dnF | GCGCCCCAGCTGGCAATTCCGG<br>GGGAATGTCGTCTCACGCAG | Amplification of downstream region of DVU2809-12, specific for marker-exchange plasmid |
| DVU2809-12-dnR | CGAGGCATTTCTGTCCTGGCTGG<br>GTTTCCGGCAAGGTCAAGG | Amplification of downstream region of DVU2809-12 |
| DVU0586-88-upF | GCCTTTTGCTGGCCTTTTGCTCACAT<br>TGGGCGTACAGTTCGGTATC | Amplification of upstream region of DVU0586-8 |

| | | |
|---|---|---|
| DVU0586-88-upR | GCGACAAGATATTCGGCACCAAGTAAG<br>GTGACAAAGCAACGCATCTTGTG | Amplification of upstream region of DVU0586-8, specific for marker-exchange plasmid |
| DVU0586-88-dnF | GCGCCCCAGCTGGCAATTCCGG<br>TCTGCCGAAGAAAGATGCCTG | Amplification of downstream region of DVU0586-8, specific for marker-exchange plasmid |
| DVU0586-88-dnR | CGAGGCATTTCTGTCCTGGCTGG<br>AGACCGTCCATCTCGTCTGC | Amplification of downstream region of DVU0586-8 |
| DVU2481-85-MLD-upR | GGGAAGGCATTAACCGATACTTG | Amplification of upstream region of DVU2481-85, specific for marker-less deletion plasmid |
| DVU2481-85-MLD-dnF | CAAGTATCGGTTAATGCCTTCCC<br>CCGACTGGATACGCAACACC | Amplification of downstream region of DVU2481-85, specific for marker-less deletion plasmid |
| DVU2809-12-MLD-upR | TCCTCTCCTTGTTGATGCCCTG | Amplification of upstream region of DVU2809-12, specific for marker-less deletion plasmid |
| DVU2809-12-MLD-dnF | CAGGGCATCAACAAGGAGAGGA<br>GGGAATGTCGTCTCACGCAG | Amplification of downstream region of DVU2809-12, specific for marker-less deletion plasmid |
| DVU0586-88-MLD-upR | GTGACAAAGCAACGCATCTTGTG | Amplification of upstream region of DVU0586-88, specific for marker-less deletion plasmid |
| DVU0586-88-MLD-dnF | CACAAGATGCGTTGCTTTGTCAC<br>TCTGCCGAAGAAAGATGCCTG | Amplification of downstream region of DVU0586-88, specific for marker-less deletion plasmid |
| pMO9075slic_F | CAAGGATCTGATGGCGCAGGG | Amplification of pMO9075 backbone |
| pMO9075slic_R | ATGGTACCTCCTGGGACTGCATTGCAGGGCTTCCCAACCT | Amplification of pMO0975 backbone |
| 2481_pmo_R | GATCGTGATCCCCTGCGCCATCAGATCCTTGTCAGGCGAAAGGACGCAGGCGCAACAA | Amplification of DvH-FDH2 from genomic DNA |

| | | |
|---|---|---|
| 2482_pmo_F | GCAGTCCCAGGAGGTACCATATGCGAATGCCTCGCAGAACGTTC | Amplification of DvH-FDH2 from genomic DNA |
| 2482_strII_R | TCATTTTTCGAACTGCGGGTGGCTCCAAGCGCTGGCCTTGCGCAGGTTGACCATGAA | Amplification of DvH-FDH2-strII |
| strII_2481_F | TGGAGCCACCCGCAGTTCGAAAAATGATGGCGCGCCATCAGAAGACTTGAT | Amplification of DvH-FDH2-strII |

[%]Underlined regions represent overhangs necessary for assembling the fragments by SLIC.

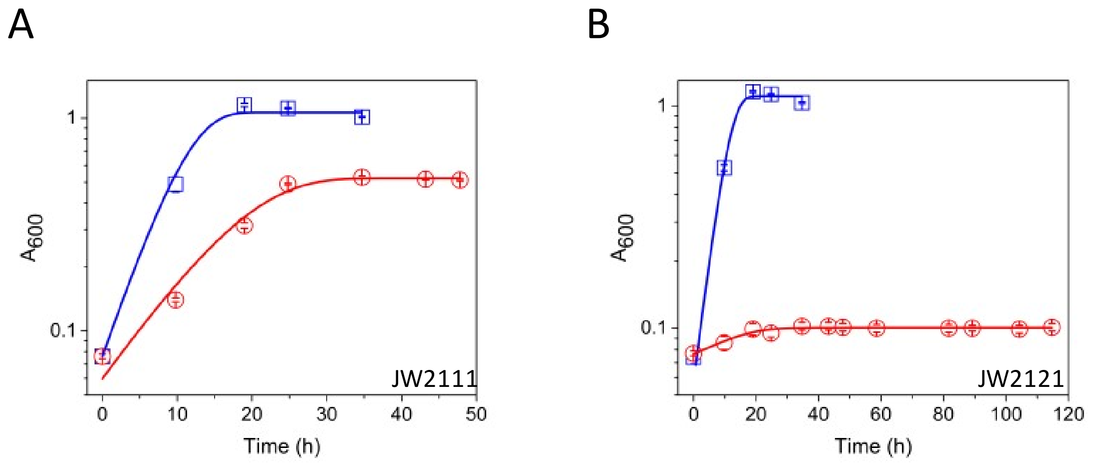

Figure S1. Growth curves of JW2111 (A) and JW2121 (B) strains of DvH. Blue and red traces represent growth on MOYLS4 and MOY-formate-acetate-S4 (MOYFAS4) media, respectively. FDH1 is the sole FDH encoded by JW2111 and is able to support growth on MOYFAS4. However, JW2121, which lacks both FDH1 and FDH3 does not. In the MOYFAS4 medium, 60 mM formate and 10 mM acetate are present to support growth but lactate is excluded.

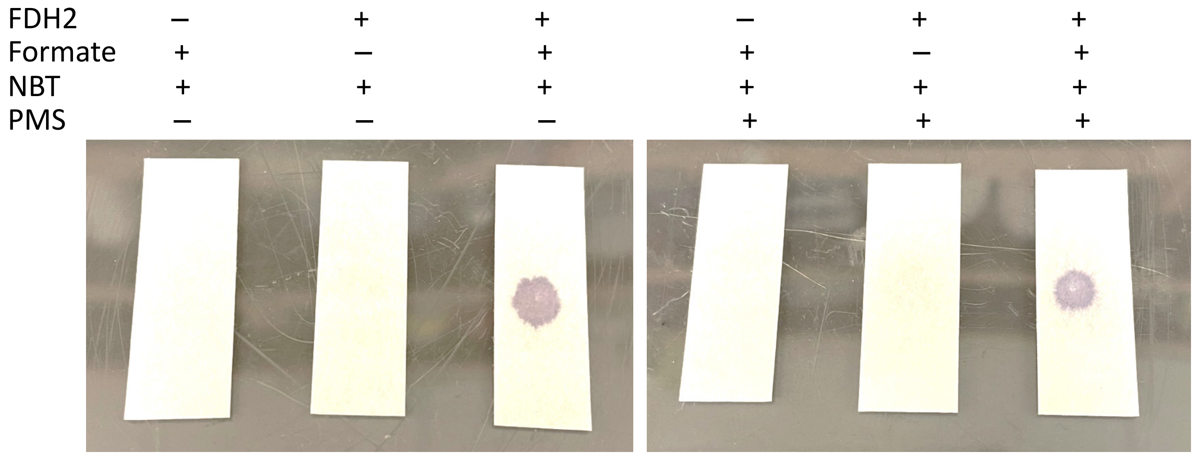

Figure S2. NBT-strip assay reveals aerobic formate oxidation by DvH-FDH2. The ability of the latter to transfer electrons to NBT is retained both in the presence and absence of PMS. A blue spot develops within 15 s only when enzyme, formate, and NBT are mixed. Although inclusion of PMS accelerates spot formation, it is not essential for the assay to succeed.

A

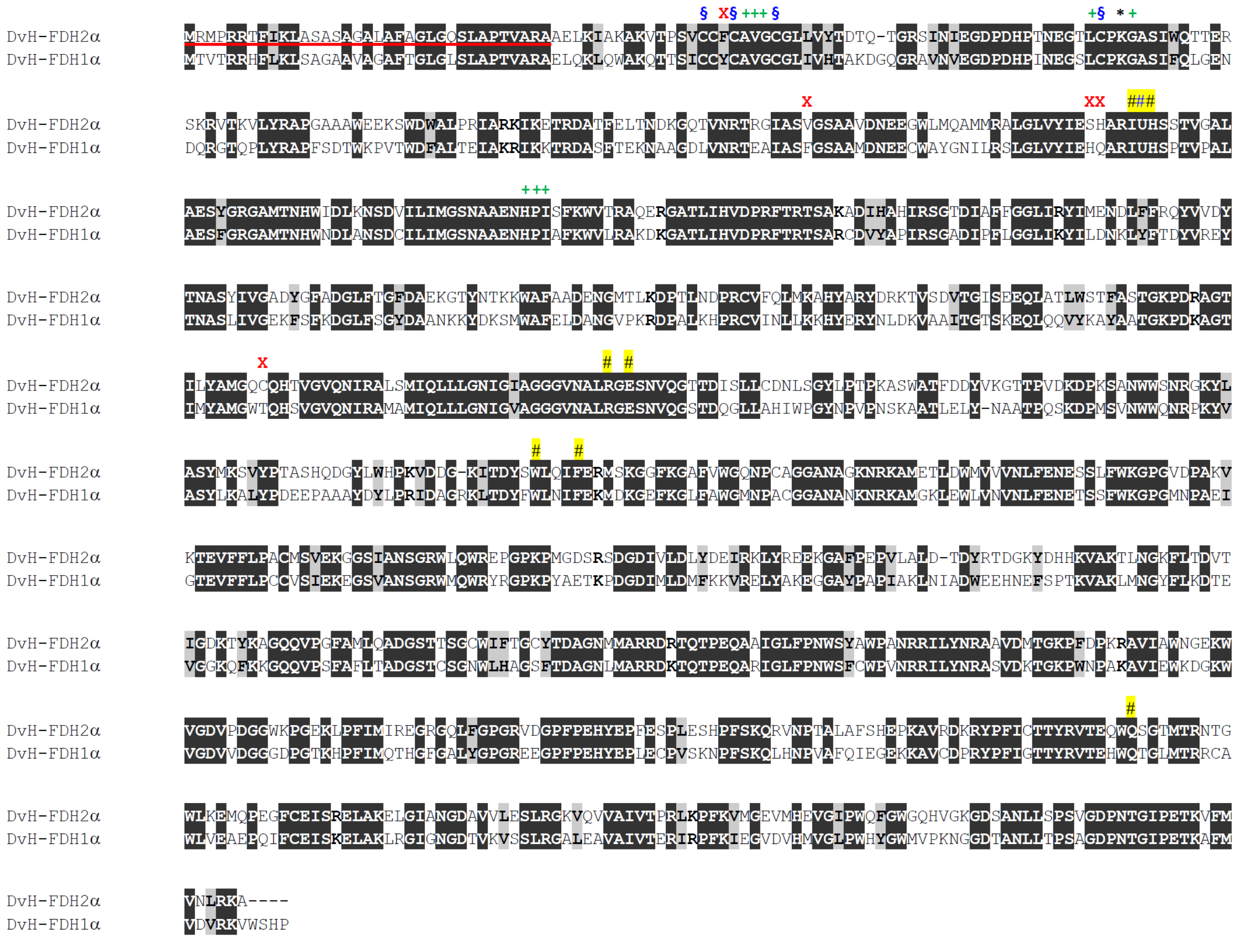

Figure S3. (A) Sequence alignment of the large subunit of DvH-FDH2 (DVU2482) with its DvH-FDH1 counterpart (DVU0587). Residues underlined in red represent the Tat (twin arginine translocation pathway) signal peptide (predicted by SignalP 6.0),[3] which is cleaved off by a peptidase (Tat/SPase I) following export to periplasmic space. Cys residues coordinating the three [4Fe-4S] clusters are identified by §. X identifies amino acids that are uniquely different between the two subunits. Sec is denoted by U.

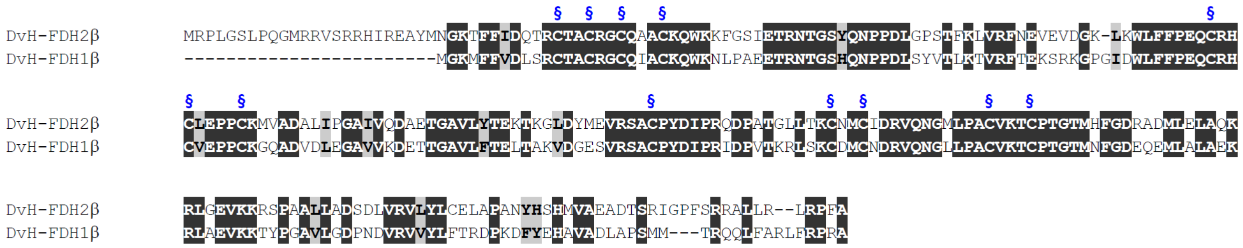

Figure S3. (B) Sequence alignment of the small subunit of DvH-FDH2 (DVU2481) with its DvH-FDH1 counterpart (DVU0588). Cys residues coordinating the three [4Fe-4S] clusters are identified by §.

Table S3: ICP-MS quantification of metal cofactors in DvH-FDH2[§]

| Metal | Predicted | Replicate 1<br>ng $mL^{-1}$ | [Metal]<br>μM | Replicate 2<br>ng $mL^{-1}$ | [Metal]<br>μM | Replicate 3<br>ng $mL^{-1}$ | [Metal]<br>μM[¶] | Observed |
|---|---|---|---|---|---|---|---|---|
| Fe | 16 | 82 ± 2.8 | 1.47 ± 0.05 | 42.8 | 0.76 | 35 ± 0.1 | 0.625 | 17 ± 1 |
| Mo | 1 | ND | ND | ND | ND | ND | ND | 0 |
| Se | 1 | 4.3 ± 0.1 | 0.055 ± 0.002 | 2.2 | 0.028 | 2.5 ± 0.1 | 0.032 | 0.7 ± 0.1 |
| W | 1 | 15.09 ± 0.02 | 0.083 ± 0.003 | 8 | 0.044 | 7.5 ± 0.1 | 0.041 | 1 ± 0.1 |

[§]Errors are standard deviations from triplicate measurements using protein samples derived from three independent preparations; ND, not detected

[¶]n=2, SEM < 0.001

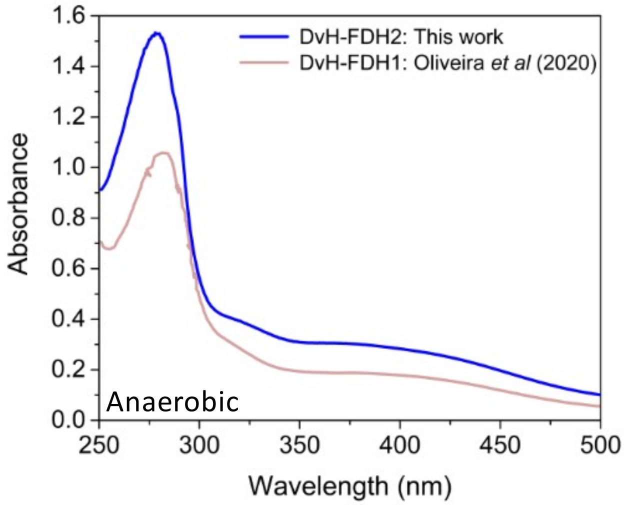

Figure S4. Comparison of the electronic spectra of DvH-FDH2 and DvH-FDH1.

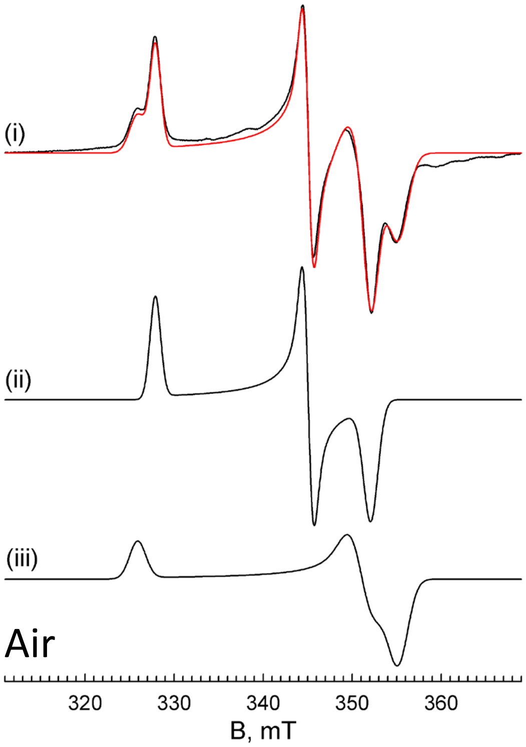

Figure S5. Simulation of Fe/S centers of formate-reduced Fdh2 prepared under aerobic conditions. (i) EPR spectrum (from Figure 4(iii)) and simulation (red trace). (ii), (iii) Scaled individual contributions to the simulation of the composite spectrum in (i). Simulation parameters are presented in Table S4.

Table S4: EPR simulation parameters for $W^{V}$ and reduced Fe/S centers of Fdh2

| | g tensors | | | | Tungsten coupling constants[a] | | | Relative contribution |
|---|---|---|---|---|---|---|---|---|
| **Center** | $g_1$ | $g_2$ | $g_3$ | $g_{ave}$ | $A_1$ | $A_2$ | $A_3$ | |
| $W^{V}1$[b] | 1.982 | 1.876 | 1.849 | 1.902 | 232 | 119 | 151 | 1.0 |
| $W^{V}2$[b] | 1.988 | 1.904 | 1.849 | 1.914 | 233[c] | 131 | 125 | 0.54 |
| Fe/S1[d] | 2.045 | 1.943 | 1.904 | 1.964 | | | | 1.0 |
| Fe/S2[d] | 2.058 | 1.910 | 1.888 | 1.952 | | | | 0.75 |

[a] In MHZ; in the absence of multi-frequency data, coupling constants are approximate [b] Spectrum collected at 108K
[c] Fixed during simulations [d] Spectrum collected at 15K

Table S5. Literature steady-state kinetics parameters of SRB-FDHs

| System | Organism | pH | T ( °C) | [BV] (mM) | [Enzyme] (nM) | $k_{cat}$ ($s^{-1}$) | $K_m$ (μM) | Reference |
|---|---|---|---|---|---|---|---|---|
| W-FDH1$^{WT}$ | Da | 8.0 | 37 | 7.5 | 35 | 241 | 10 | Mota (2011); Mota et al (2011) |
| W-FDH1$^{WT}$ | Dg | 8.0 | 37 | 7.5 | 35 | 174 | 51 | Mota (2011); Mota et al (2011) |
| W-FDH1$^{WT}$ | DvH | 7.6 | RT | 2 | 0.0124 | 3684 | 1 | Da Silva *et al* (2011) |
| W-FDH1$^{REC}$ | DvH | 7.6 | RT | 2 | 1.4 | 1100 | NR | Miller *et al* (2018) |
| W-FDH1$^{REC}$ | DvH | 7.6 | RT | 2 | 1.4 | 940 | NR | Szczesny *et al* (2019) |
| W-FDH1$^{WT}$ | DvH | 7.6 | RT | 2 | 1.4 | 1104 ± 62 | NR | Oliveira *et al* (2020) |
| W-FDH1$^{REC}$ | DvH | 7.6 | RT | 2 | 1.4 | 1310 ± 50 | 16.9 ± 2.8 | Oliveira *et al* (2020) |
| W-FDH1$^{REC}$ | DvH | 7.6 | RT | 2 | 1.4 | 1144 | NR | Alvarez-Malmagro *et al* (2021) |
| **??-FDH2$^{WT}$** | **DvH** | 7.6 | RT | 2 | 0.48 | 81 | 4 | Da Silva *et al* (2011) |
| Mo-FDH3$^{WT}$ | Dd | 7.6 | 37 | 7.5 | 35 | 357 ± 18 | 65 ± 8 | Rivas *et al* (2007) |
| Mo-FDH3$^{WT}$ | Dd | 8.0 | 37 | 7.5 | 35 | 347 | 64 | Mota (2011); Mota *et al* (2011) |
| Mo-FDH3$^{WT}$ | DvH | 7.6 | RT | 2 | 0.25 | 262 | 8 | Da Silva *et al* (2011) |
| Mo-FDH3$^{WT}$ | Dd | 8.0 | 22 | 5 | 1 | 543 | 57 | Maia *et al* (2016) |

SRB, sulfate-reducing bacteria; WT, wild-type natively-purified protein; REC, recombinant; W-, tungsten-containing; Mo-, molybdenum-containing; ??, metal status unknown; NR, not reported; likely to be similar to the value reported by Oliveira *et al* (2020); RT, room temperature; BV, benzyl viologen; DvH, *Desulfovibrio vulgaris* Hildenborough; Dd, *D. desulfuricans*; Dg, *D. gigas*; Da, *D. alaskensis*. Additional experimental details shared by Drs. Luisa Maia and Inês Pereira have been included here for the sake of completeness. The FDH probed in the present work is identified in blue.

$$E_{ox} + F \underset{0}{\overset{k_{+1}}{\rightleftharpoons}} E_{ox}F \underset{0}{\overset{k_{+2}}{\rightleftharpoons}} E_{red}CO_2 \underset{0}{\overset{10^4\ s^{-1}}{\rightleftharpoons}} E_{red} + CO_2$$

$$E_{red} + BV_{ox} \underset{0}{\overset{100\ \mu M^{-1}\ s^{-1}}{\rightleftharpoons}} E'_{ox} + BV_{red}$$

$$E'_{ox} + BV_{ox} \underset{0}{\overset{100\ \mu M^{-1}\ s^{-1}}{\rightleftharpoons}} E_{ox} + BV_{red}$$

**Scheme S1**: Minimal catalytic model used solely for estimating $k_{cat}/K_m$ ($k_{+1}$) and $k_{cat}$ ($k_{+2}$) from steady-state and full progress curves via dynamic simulation-based global fitting of BV data. This model takes into consideration both the $2e^-$ oxidation of formate and $1e^-$ reduction of BV, leading to a stoichiometry of $2BV^+$:1F. See Axley and Grahame[4] for additional details regarding the redox reactions. Also see Scheme 1.

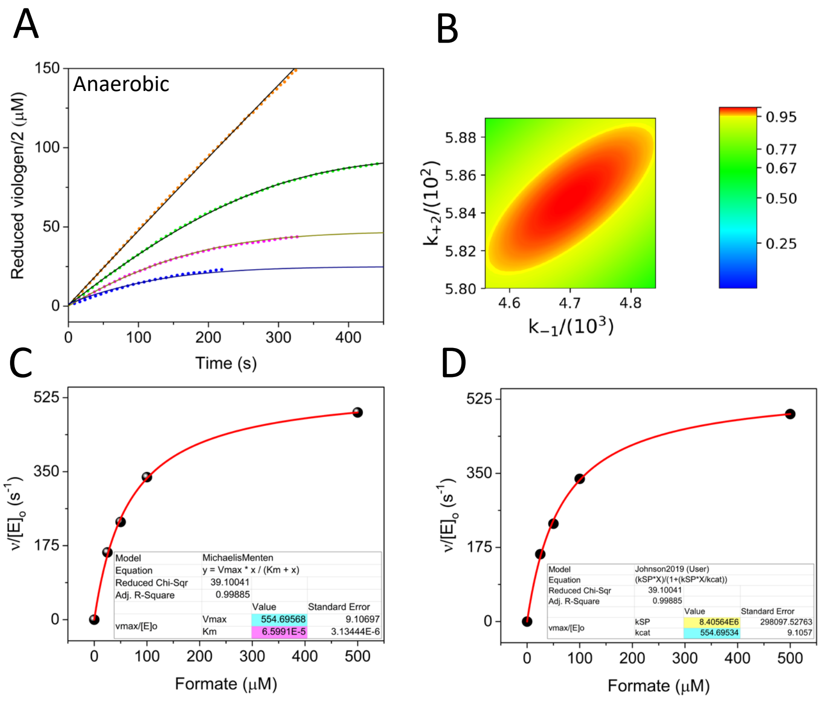

E

| Method | $k_{cat}$ ($s^{-1}$) | $K_m$ (μM) | $k_{cat}/K_m$ ($\mu M^{-1}\ s^{-1}$) |
|---|---|---|---|
| Equations¶ | 543 | 57 | 9.5 |
| Equations§ | 555 | 66 | 8.4 |
| Simulation# | 585 ± 4 | 47 ± 2.3 | 12.4 ± 0.6* |

¶Hanes plot – linear regression of Maia *et al* (2016)

§ICEKAT: 0-500 μM formate. Michaelis-Menten nonlinear regression (this work). Since the original traces were already normalized to take into account 2BV:1F stoichiometry (panel A), $k_{cat}$ values have not been divided by two.

#Kintek Explorer global fit (errors via FitSpace Explorer) (this work)

*Standard error values for $k_{cat}$ / $K_m$ (calculated from $k_{cat}$ and $K_m$) estimated according to Johnson (2019)

Figure S6. Validating Scheme S1 using source[5] Dd-FDH3 BV kinetics data. (A) Raw traces (points) and the associated global fits (solid lines). (B) Confidence contour analysis (for details on how to interpret this plot, see legend to Figure 5). (C) Classical Michaelis-Menten analysis. (D) Fitting the data in panel C to extract $k_{cat}$ amd $k_{cat}/K_m$ ($k_{SP}$) using the approach described by Johnson.[6] Initial velocities were obtained using ICEKAT.[7] (E) Comparison of kinetic parameters derived from three different approaches.

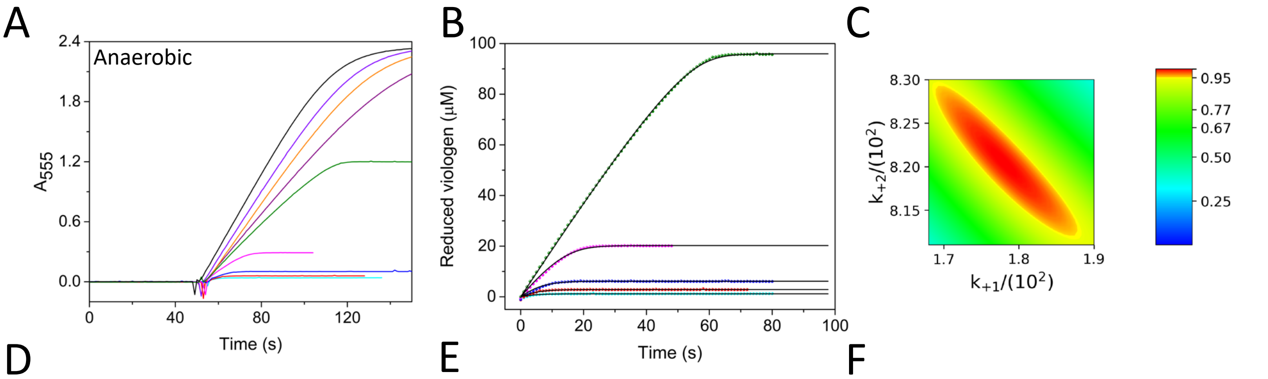

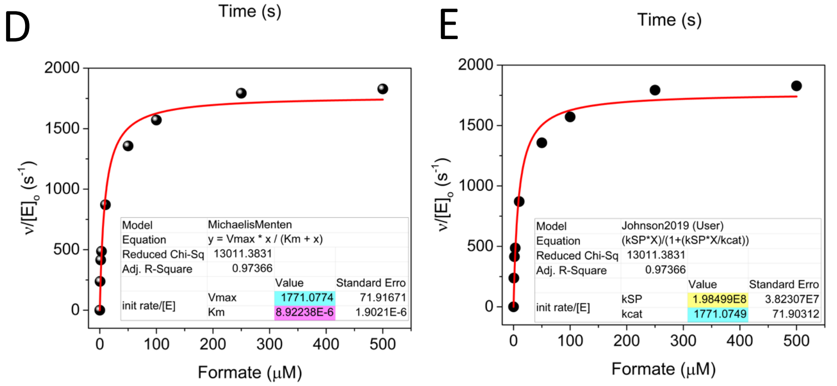

F

| Method | $k_{cat}$ ($s^{-1}$) | $K_m$ (µM) | $k_{cat}/K_m$ ($\mu M^{-1}$ $s^{-1}$) |
|---|---|---|---|
| Equations¶ | 1310 ± 50 | 16.9 ± 2.9 | 77.5 |
| Equations§ | 885* | 8.9 | 99.4 |
| Simulation# | 820 ± 9 | 4.6 ± 0.3 | 178 ± 10 |

¶Nonlinear regression of Oliveira *et al* (2020)
§ICEKAT: 0-500 µM formate (this work)
#Kintek Explorer: Full progress curve analysis (0.5-50 µM formate) (errors via FitSpace Explorer) (this work)
*Values obtained from panels D and E were divided by two to account for 2BV:1F stoichiometry

Figure S7. Ground truthing Scheme S1 with DvH-FDH1 source[8] data. (A) Raw kinetic traces of BV reduction. (B) Full progress curves (points) extracted from panel A for global fitting analysis. Fits are shown as solid lines. (C) Confidence contour analysis. (D) Nonlinear regression of Michaelis-Menten equation. (E) Fitting the initial velocity data according to Johnson.[6] (F) Summary of the results.

Table S6. Steady-state kinetics of DvH-FDH2[#]

| Enzyme | Electron acceptor | Reaction condition | $k_{cat}$ ($s^{-1}$) | $K_m$ (μM) | $k_{cat}/K_m$ ($\mu M^{-1}$ $s^{-1}$)[‡] | Replicates | Reference |
|---|---|---|---|---|---|---|---|
| $FDH2^{WT}$ | BV (2 mM) | Anaerobic | 81 | 4 | 20 | 1 | da Silva *et al* (2011) |
| $FDH2^{Rec}$ | BV (2 mM) | Anaerobic | 68 ± 5 | 3.5 ± 0.9 | 19.4 ± 5.2 | 3 | This work |
| $FDH2^{Rec}$ | BV (20 mM) | Anaerobic | 111 ± 16 | 7 ± 3 | 15.8 ± 7.1 | 3 | This work |
| $FDH2^{Rec}$ | PES/DCPIP | Anaerobic | 220 ± 5 | 5.5 ± 0.5 | 40 ± 4 | 5 | This work |
| $FDH2^{Rec}$ | PES/DCPIP | Air | 317 ± 14 | 7 ± 1 | 45.3 ± 6.3 | 10 | This work |

[#]Initial velocities were calculated using ICEKAT, utilizing the first 10 – 12 s of data.

[‡]Standard error values for $k_{cat}$ /$K_m$ estimated according to Johnson (2019)

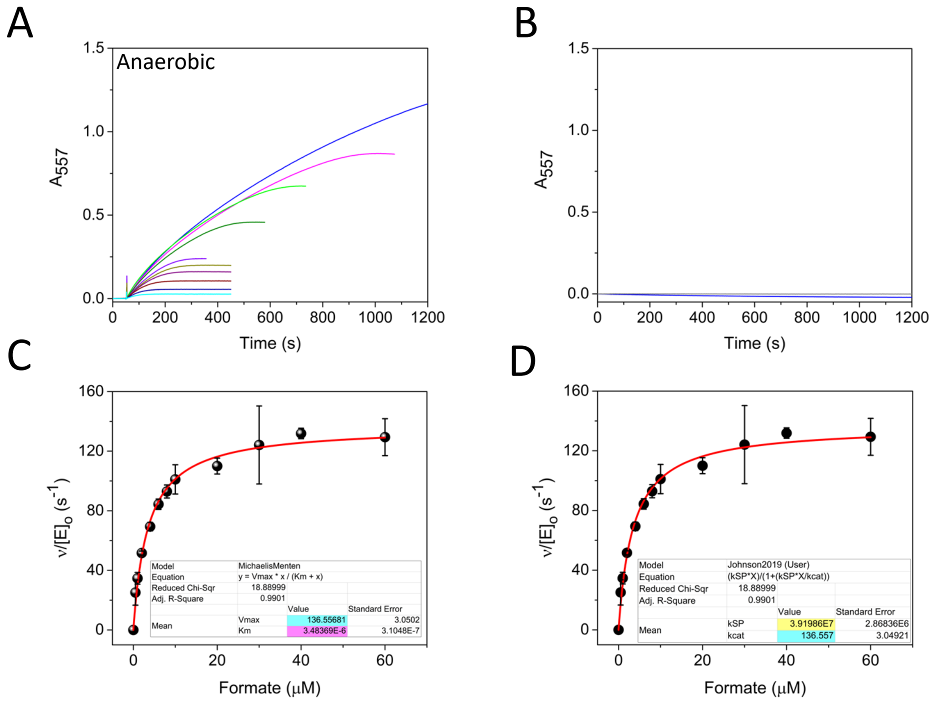

Figure S8. Reduction of 2 mM BV by DvH-FDH2 as a function of formate concentration (0 – 60 μM). (A) Raw kinetic traces. (B) No formate (grey) and no enzyme (blue) controls. (C) Nonlinear regression of Michaelis-Menten equation. (D) Fitting according to Johnson.[6] Error bars represent standard deviation of three independent measurements. Error bars represent standard deviation from three independent measurements.

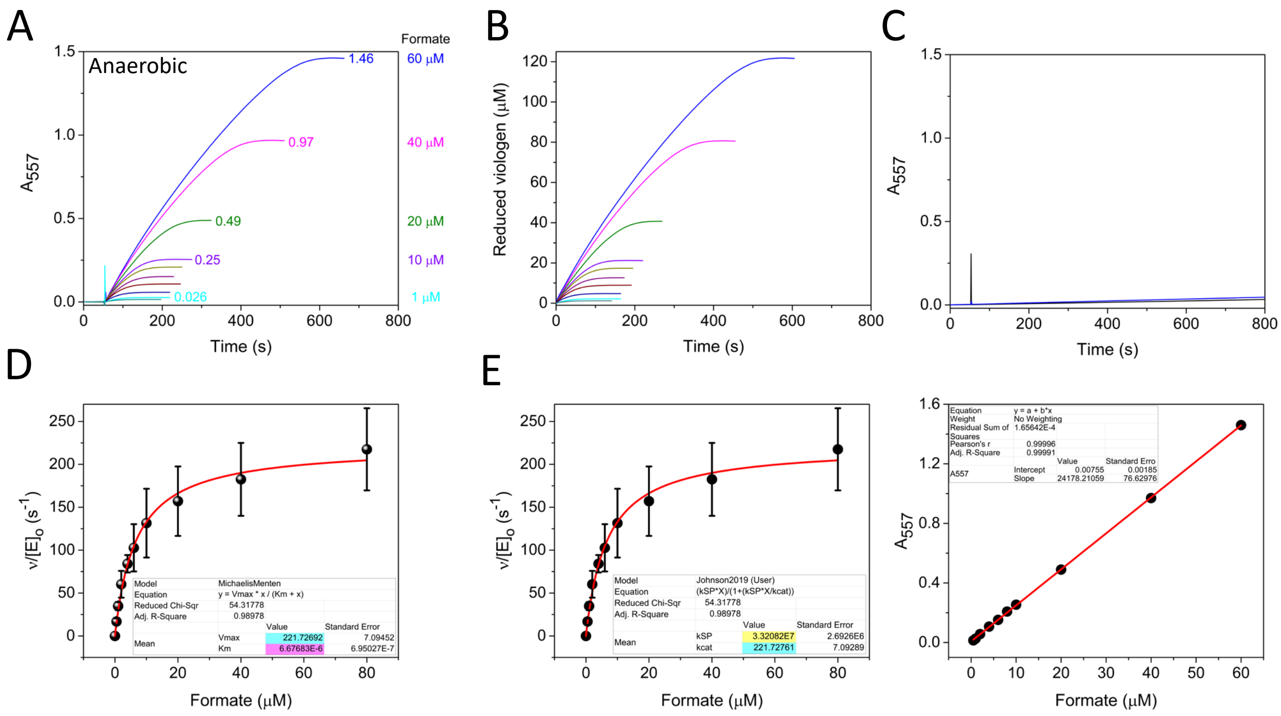

Figure S9. DvH-FDH2 catalyzed reduction of 20 mM BV as a function of formate concentration. (A) Raw kinetic traces are shown alongside endpoint values. Formate concentration is listed on the right Y axis. (B) Concentration normalized traces (points) and the global fits (solid lines going through the points). (C) No formate (grey) and no enzyme (blue) controls. (D) Nonlinear regression of Michaelis-Menten equation. (E) Nonlinear regression of Johnson's[6] equation. Error bars represent standard deviation (n=3). (F) Determination of BV extinction coefficient from full progress curve endpoints. Plotting the latter values as a function of [formate] results in a slope, which after correcting for $2BV^+$:1F stoichiometry, yields a value of 12,089 ± 38 $M^{-1}$ $cm^{-1}$. This is yet another way to validate stoichiometric reduction of BV by DvH-FDH2. Our approach to the determination of BV extinction coefficient is similar to that employed using hydrogenase as an electron donor.[9] Literature molar extinction values for BV range from 7.0 – 19.5 $mM^{-1}$ $cm^{-1}$ depending on the wavelength of measurement (546, 555, 578, 580 or 600 nm)[5, 8-20]. Because source kinetics data involving artificial electron acceptors is never reported in the literature, it is not possible to properly interpret/validate the kinetic parameters/reaction stoichiometry of a metallo-FDH. When combined with the fact that the reaction product is also seldom measured, the results turn out to be largely phenomenological and lack information content essential for making technological advances.

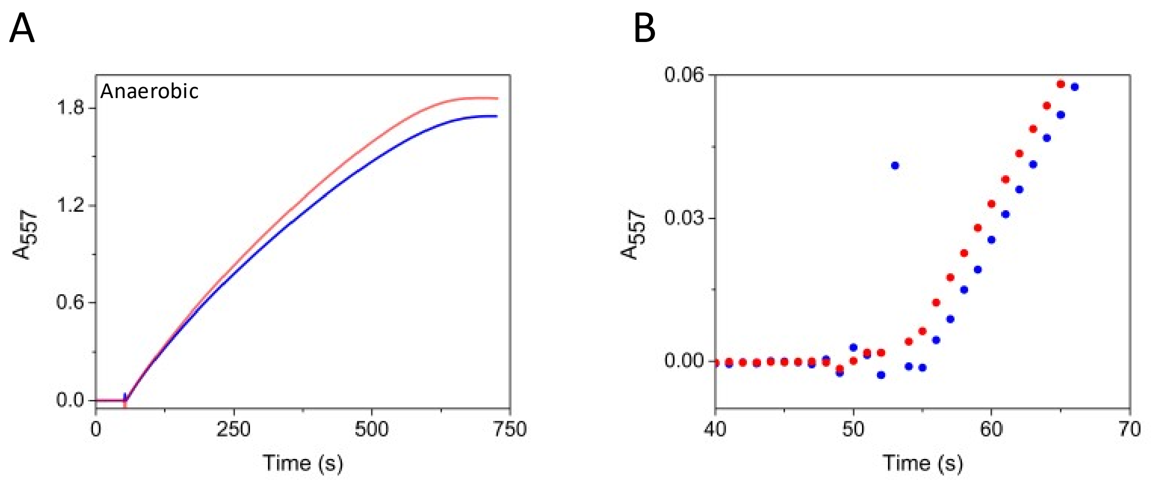

Figure S10. Glucose oxidase (GO) does not interfere with BV reduction by DvH-FDH2. (A) In the absence (blue trace) and presence (red trace) of 1 U GO. (B) Closeup view illustrating virtually identical slopes for the two traces between 55 – 65 s (region used for initial velocity estimation in ICEKAT).

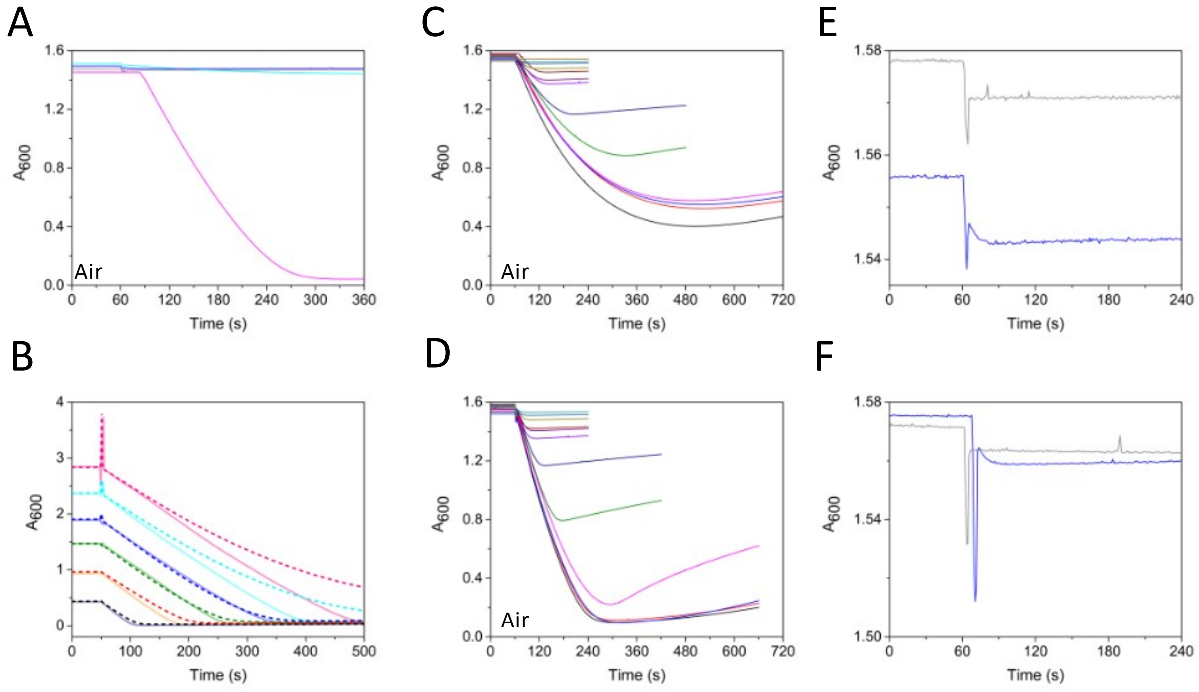

Figure S11. Optimization of experimental conditions for PES/DCPIP reduction. (A) DCPIP reduction occurs only when PES is present (pink trace). Otherwise, there is no signal change at 600 nm when FDH2 is combined with DCPIP alone (cyan). No DCPIP control is shown in grey. (B) Varying DCPIP while maintaining a fixed PES concentration. Solid and dashed lines represent measurements performed under aerobic and anaerobic conditions, respectively. (C) Varying PES concentration while keeping DCPIP fixed. (D) Optimized condition; (E,F), no formate (blue) and no enzyme (grey) controls.

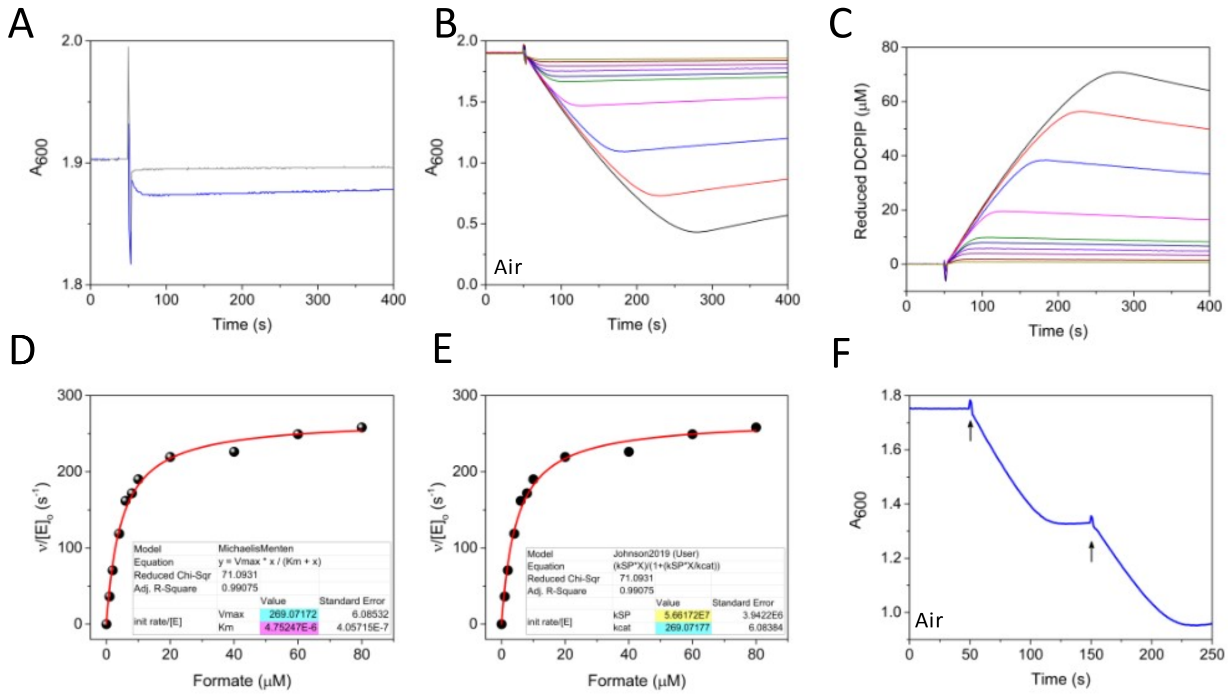

Figure S12. PES/DCPIP reduction by DvH-FDH2 in air. (A) no enzyme (grey) and no formate (blue) controls. (B) Raw kinetic traces as a function of varying formate concentration. DvH-FDH2, DCPIP, and PES levels were fixed. (C) Concentration-normalized and inverted (product increases as a function of time) traces. (D) Non-linear least squares fit to the classical Michaelis-Menten equation. (E) Fit to Johnson's equation[6] for extracting $k_{cat}$ and $k_{cat}/K_m$. Initial velocities were obtained via ICEKAT[7]. Fit parameters are included within the plots. (F) Demonstration that enzyme activity is not lost after plateauing. Addition of a second formate aliquot (up arrow) restores the original progress curve. When possible, this approach is superior to the Selwyn test,[21] which requires progress curve measurements at different [enzyme] but fixed [substrate]. Unfortunately, it lacks the ability to offer insights into the inactivation rates or the extent of enzyme inactivation.

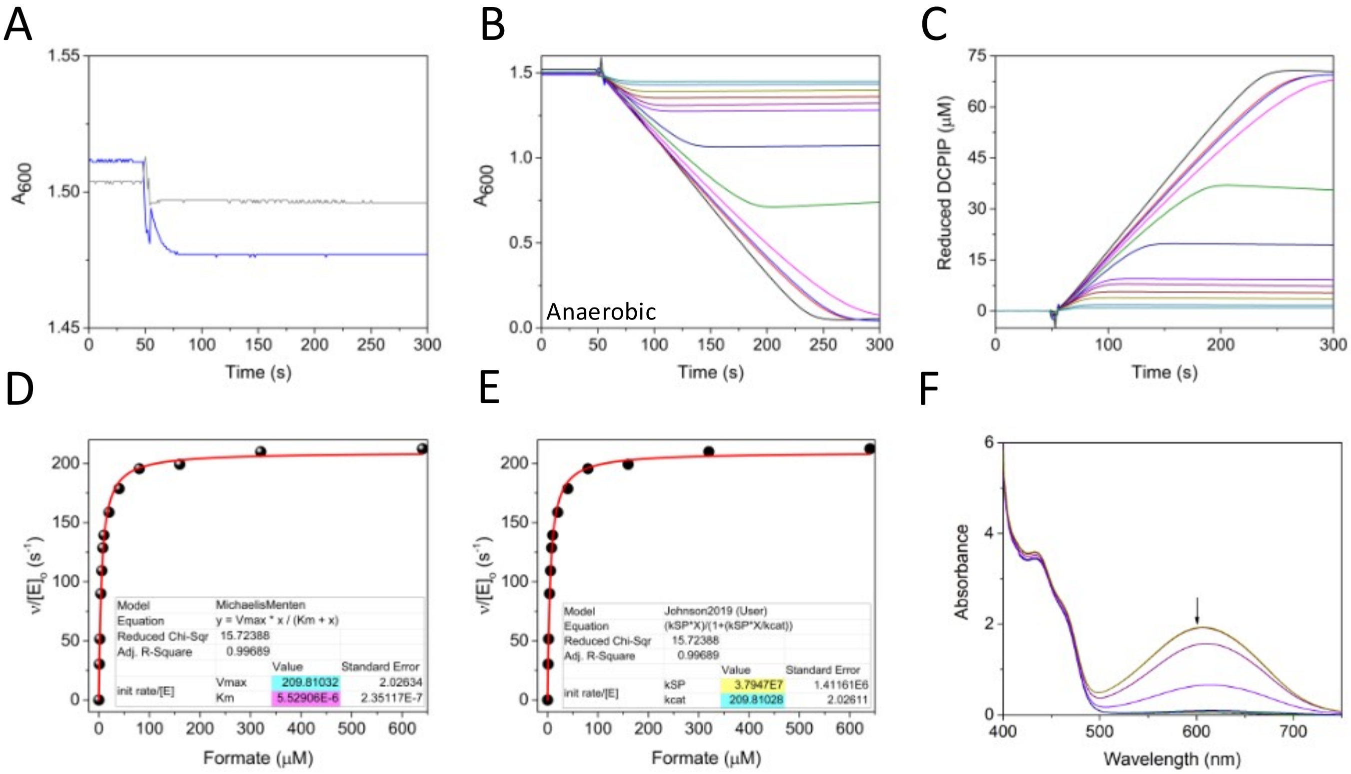

Figure S13. PES/DCPIP reduction by DvH-FDH2 under anaerobic conditions. (A) No enzyme (grey) and no formate (blue) controls. (B) Raw kinetic traces as a function of varying formate concentration. DvH-FDH2, DCPIP, and PES are fixed. (C) Concentration-normalized and inverted traces. (D) Non-linear least squares fit to the classical Michaelis-Menten equation. (E) Fit to Johnson's equation[6] for extracting $k_{cat}$ and $k_{cat}/K_m$. Initial velocities were obtained via ICEKAT[7]. Fit parameters are included within the plots. (F) Spectral changes associated with PES/DCPIP reduction. Absorbance at 600 nm (down arrow) decreases as DCPIP is reduced.

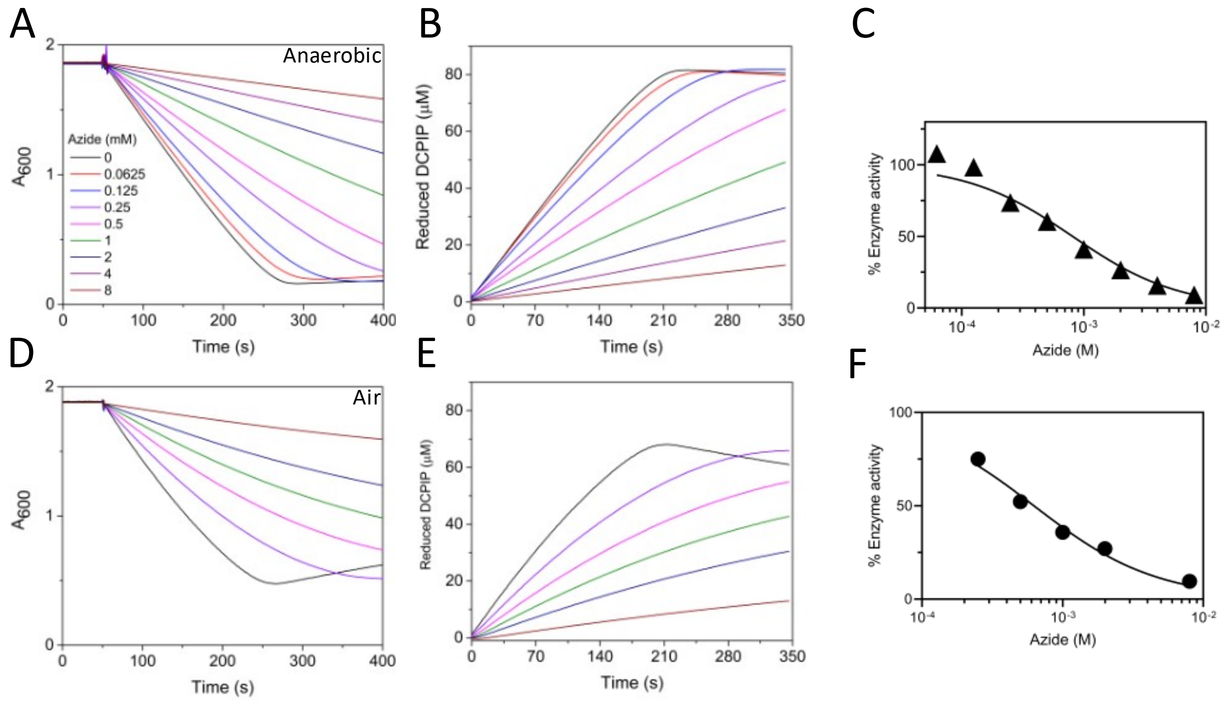

Figure S14. Effect of sodium azide on PES/DCPIP reduction by DvH-FDH2. Anaerobic and (A, B, C) aerobic (D,E,F) kinetics data are shown. A four-parameter sigmoidal function was used to derive the $IC_{50}$ values (C, F), which are in the range of 0.8 mM.

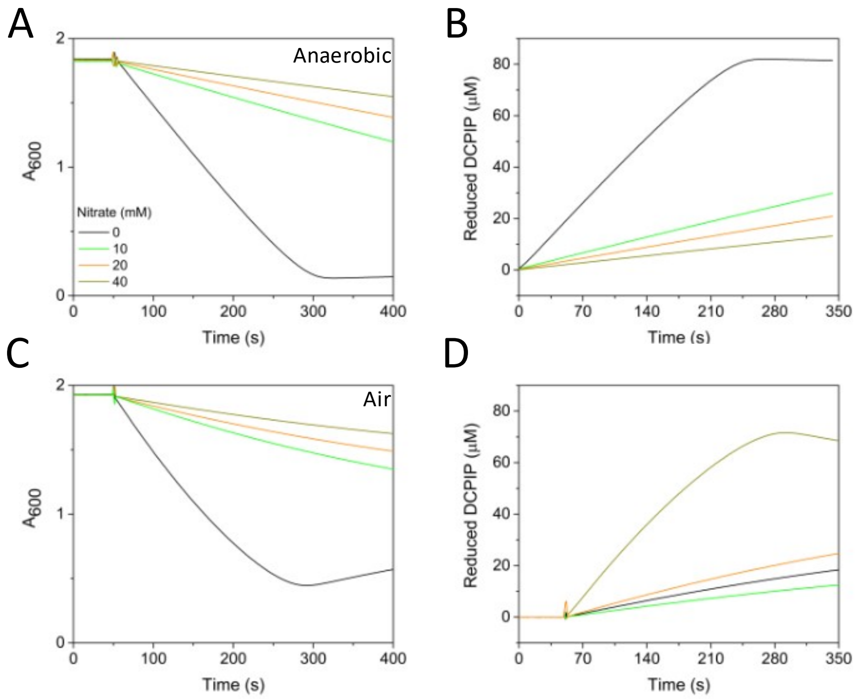

Figure S15. Effect of sodium nitrate on PES/DCPIP reduction by DvH-FDH2. Anaerobic (A, B) and aerobic (C,D) kinetics panels are shown.

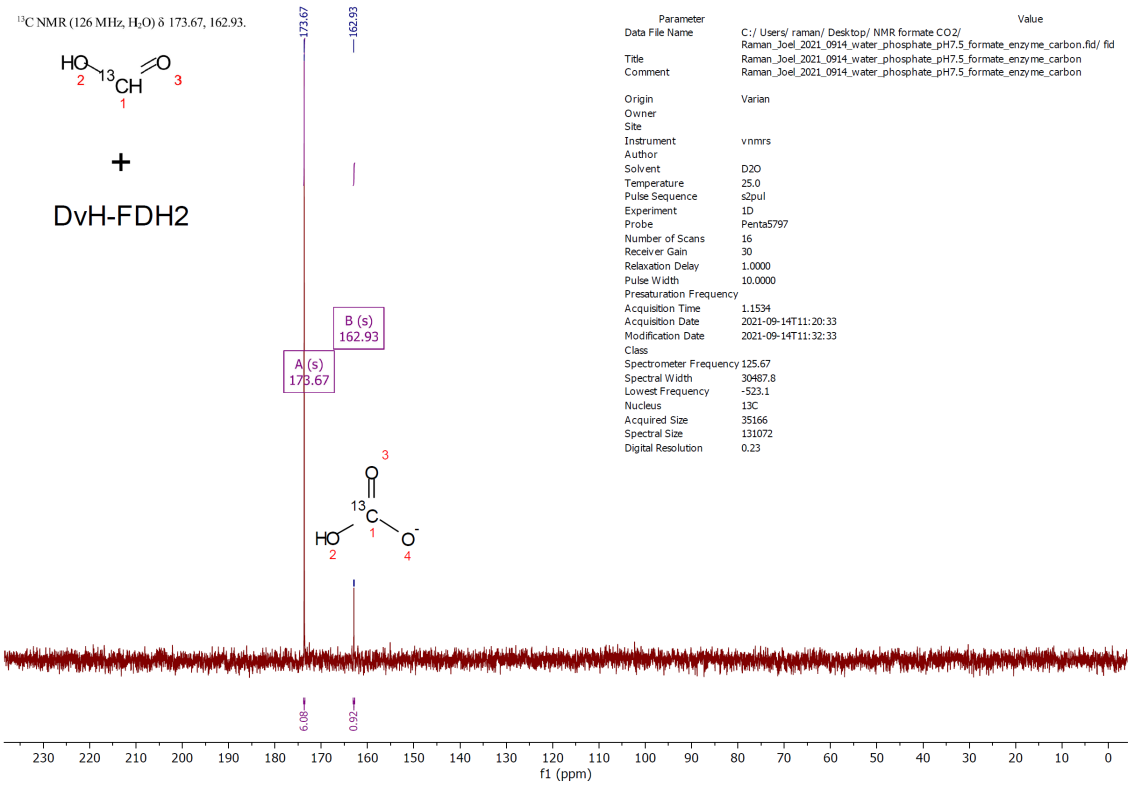

Figure S16. $^{13}$C NMR spectrum of $^{13}$C-formate + DvH-FDH2 at pH 7.5.

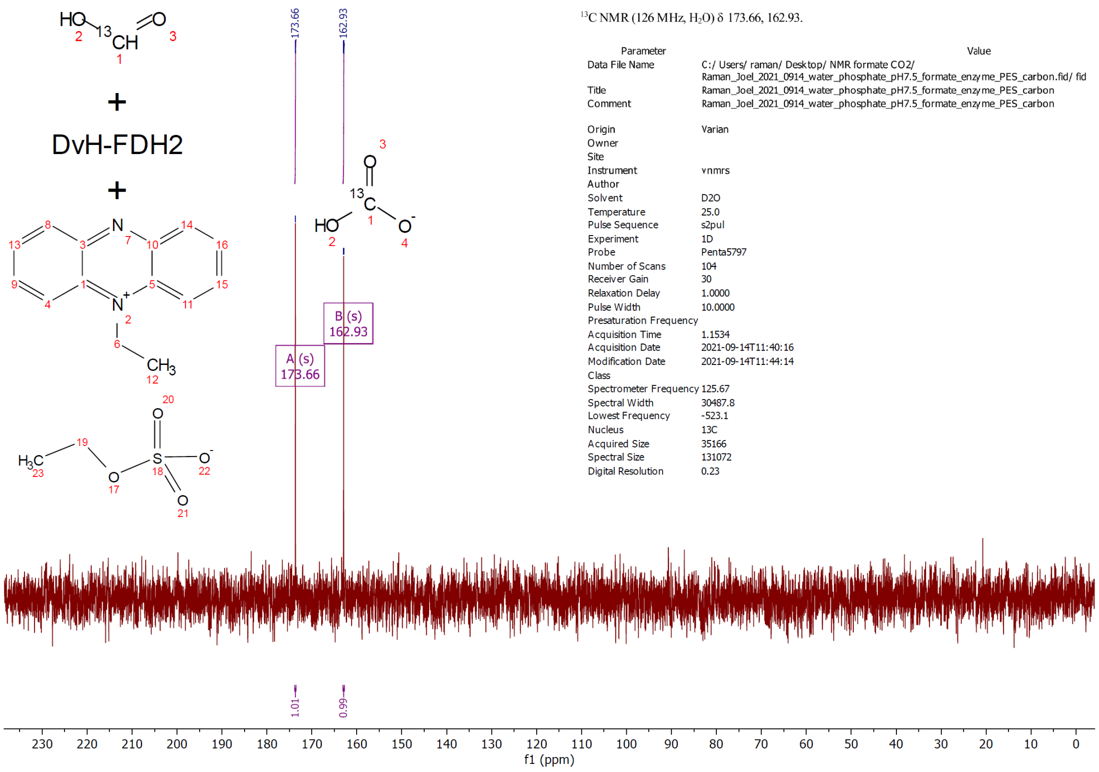

Figure S17. $^{13}$C NMR spectrum of $^{13}$C-formate + DvH-FDH2 + PES at pH 7.5.

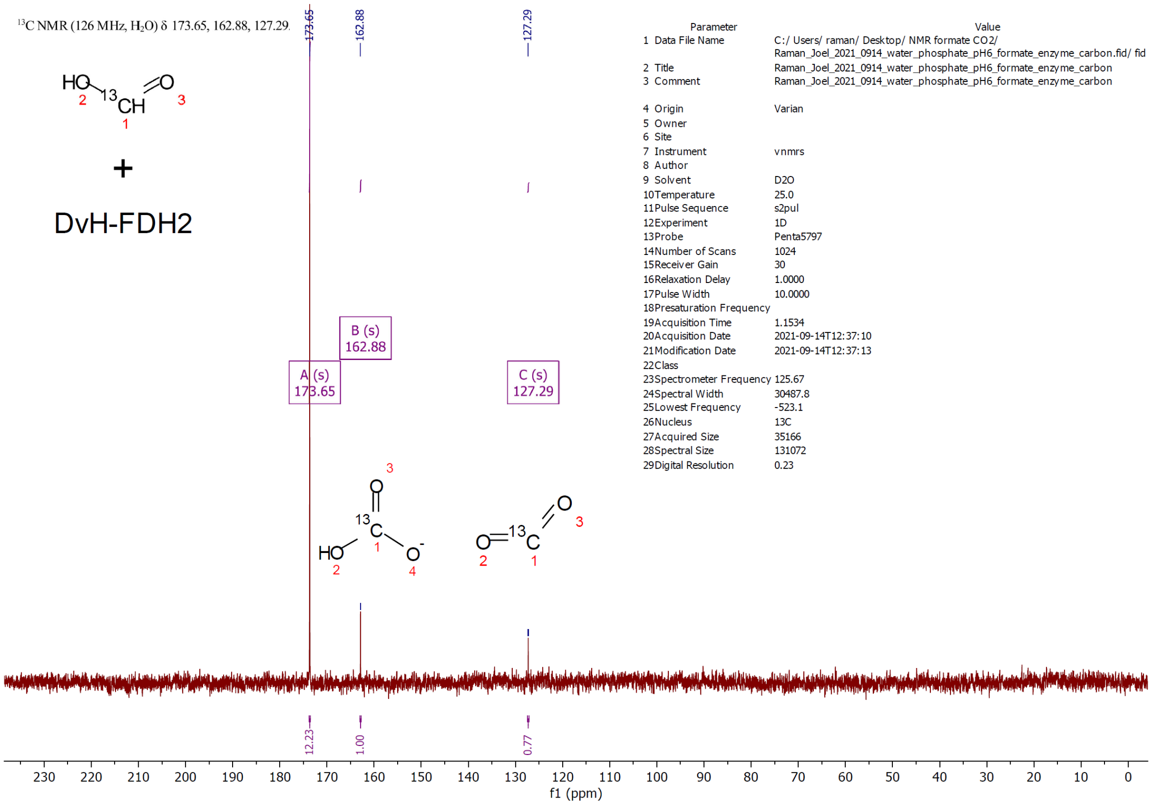

Figure S18. $^{13}$C NMR spectrum of $^{13}$C-formate + DvH-FDH2 at pH 6.0.

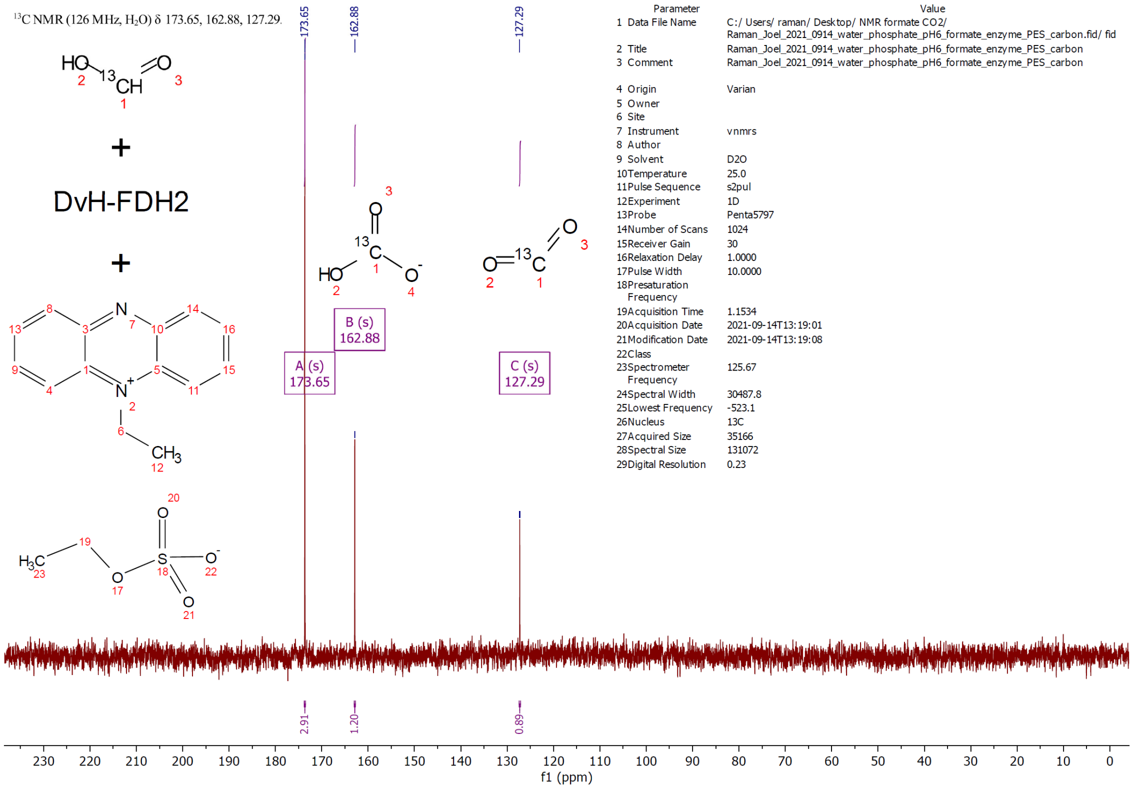

Figure S19. $^{13}C$ NMR spectrum of $^{13}C$-formate + DvH-FDH2 + PES at pH 6.

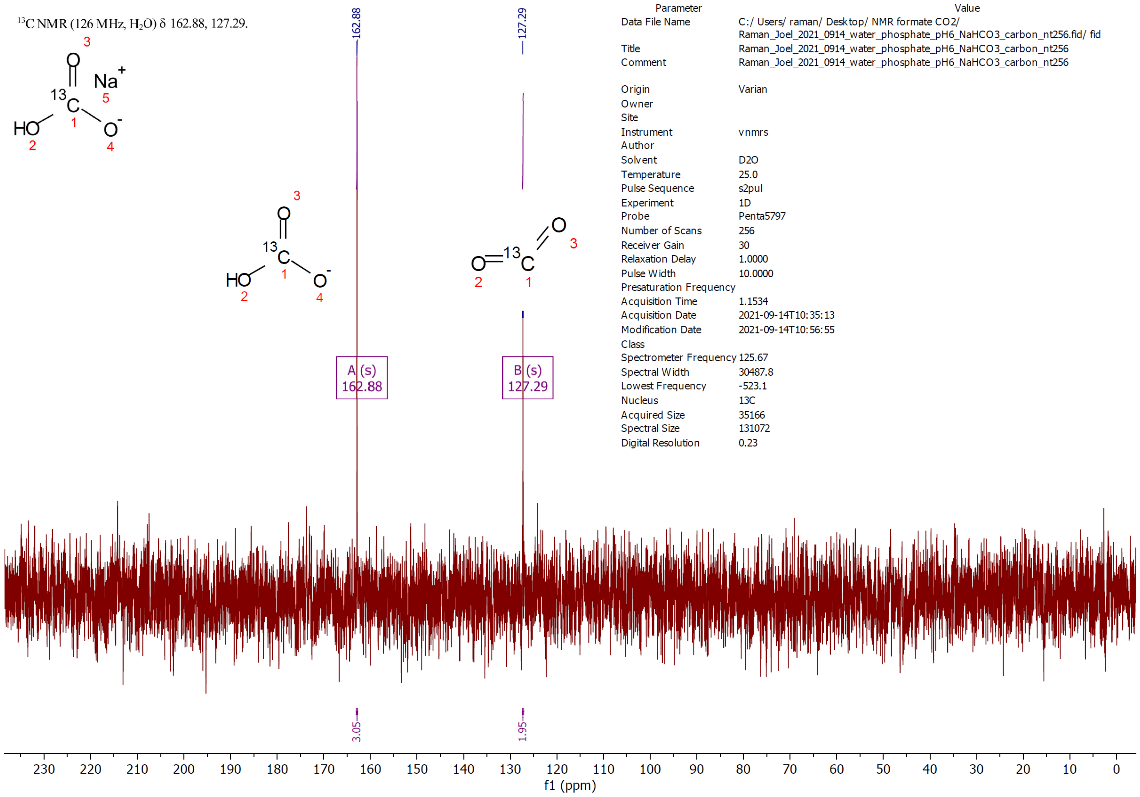

Figure S20. $^{13}$C NMR spectrum of isotopically enriched (99%) $^{13}$C-sodium bicarbonate at pH 6.

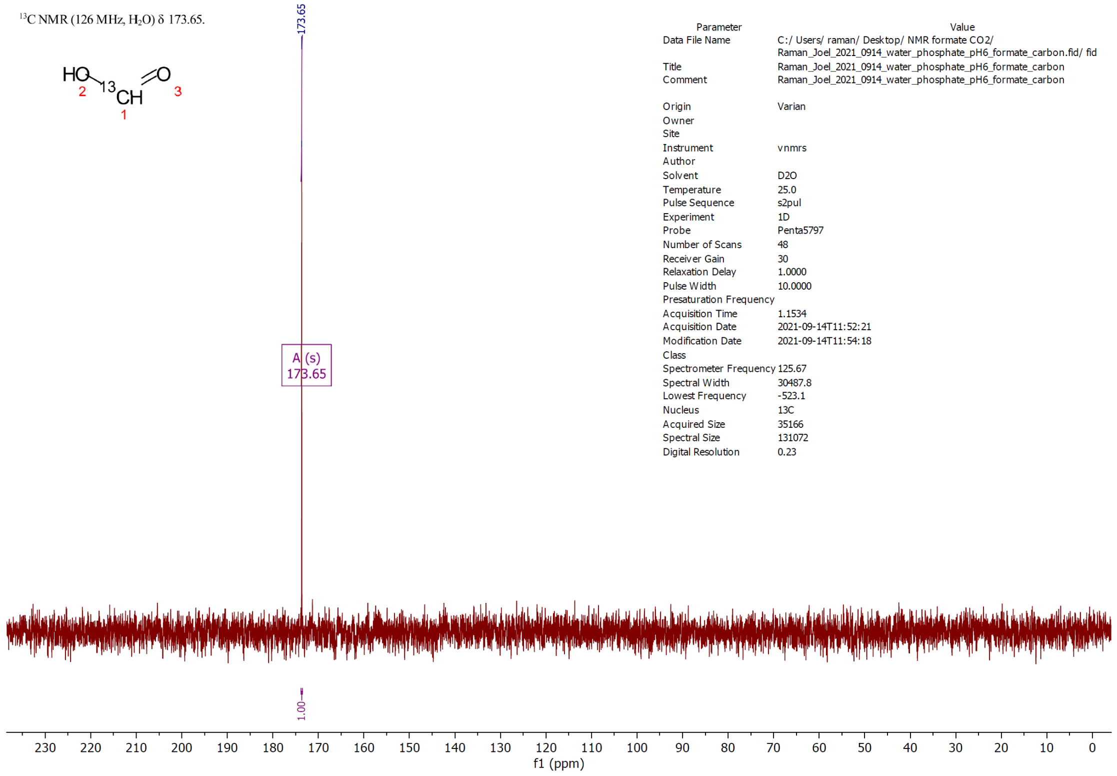

Figure S21. $^{13}$C NMR spectrum of isotopically enriched (99%) $^{13}$C-formate at pH 6.0.

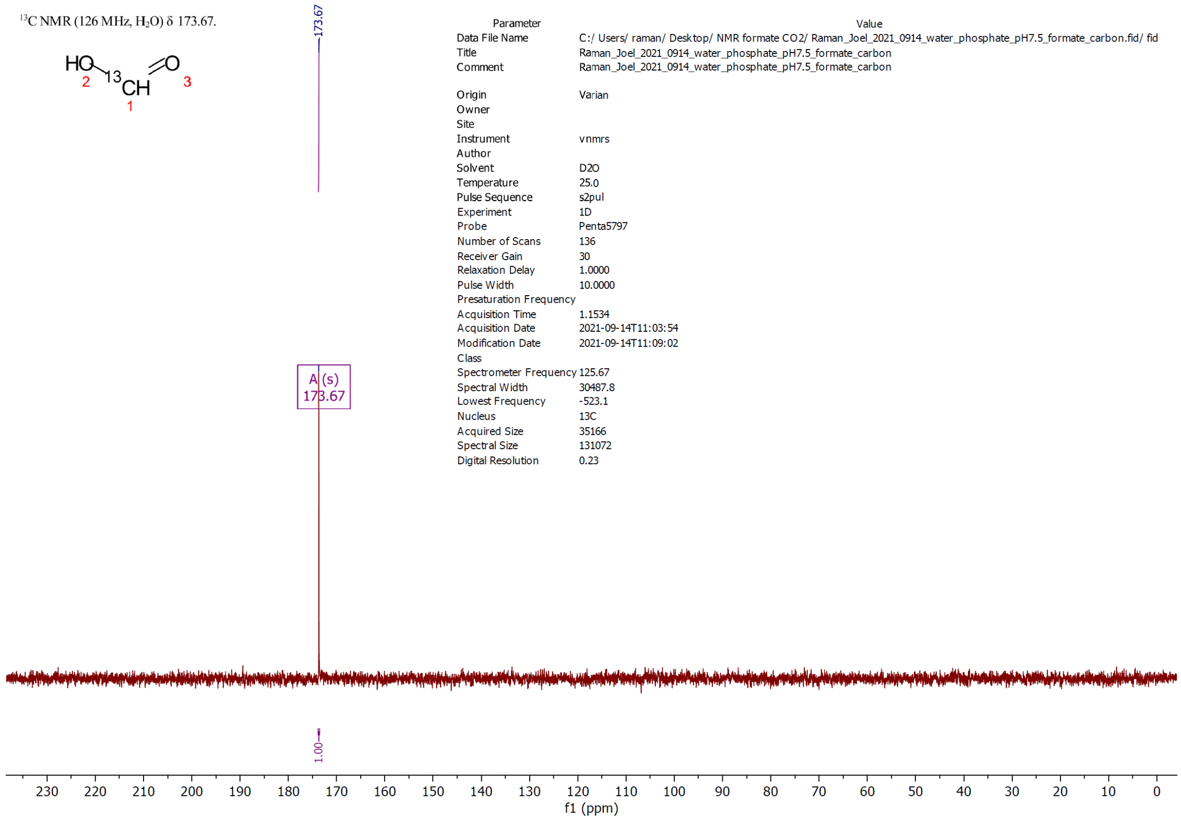

Figure S22. $^{13}C$ NMR spectrum of isotopically enriched (99%) $^{13}C$-formate at pH 7.5.

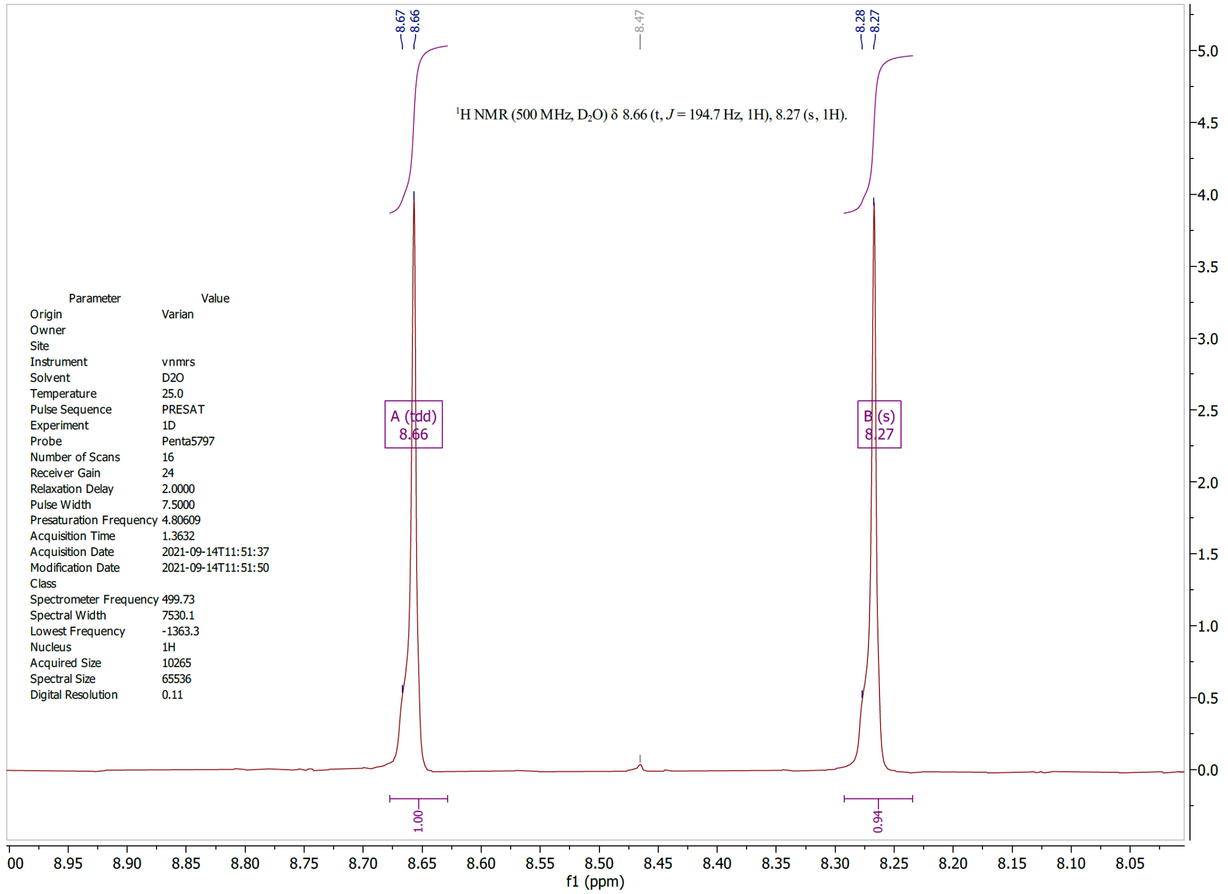

Figure S23. $^{1}H$ NMR spectrum of isotopically enriched (99%) $^{13}C$-formate at pH 6. The formyl singlet splits into a doublet in the $^{1}H$ spectrum due to the coupling of $^{1}H$-$^{13}C$ ($J$ ~ 195 Hz). Trace (1%) $^{12}C$-formate is visble as a singlet (8.47 ppm) sandwiched in between the doublet.

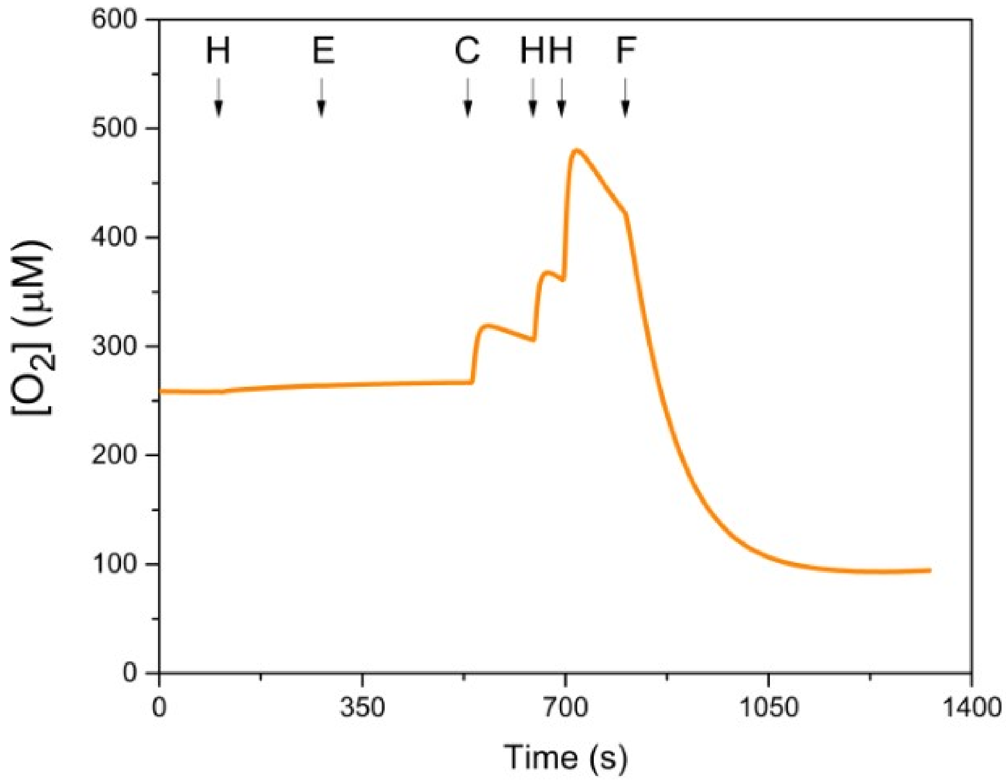

Figure S24. DvH-FDH2 lacks catalase activity. Upon incubation of $H_2O_2$ (H) and the enzyme (E), no $O_2$ evolution was observed. Subsequent addition of catalase (C), however, led to $O_2$ production, which was enhanced by two more $H_2O_2$ aliquots. Ultimately, the addition of formate (F) resulted in $O_2$ consumption. Despite experiencing an $O_2$ concentration of ca. 32%, the enzyme functioned normally.

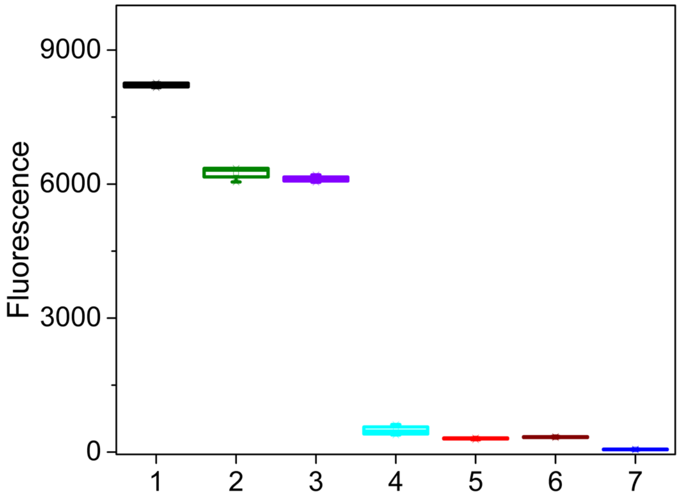

Figure S25. Amplex red assay at a fixed concentration (5 μM) of formate (F) (n=3). Abscissa refers to sample numbers. $H_2O_2$ standard (5 μM) (black), (2) FDH2 + F (green),. FDH2 + F + SOD (10 U/mL) (violet), FDH2 + F + catalase (100 U/mL) (cyan), heat denatured FDH2 + F (red), FDH2 alone (brown), and FDH2 + F sans HRP (blue). Unless specified otherwise, all samples contained buffer, DTPA, amplex red, HRP. 1.6 nM FDH2 was used. See Methods for additional details.

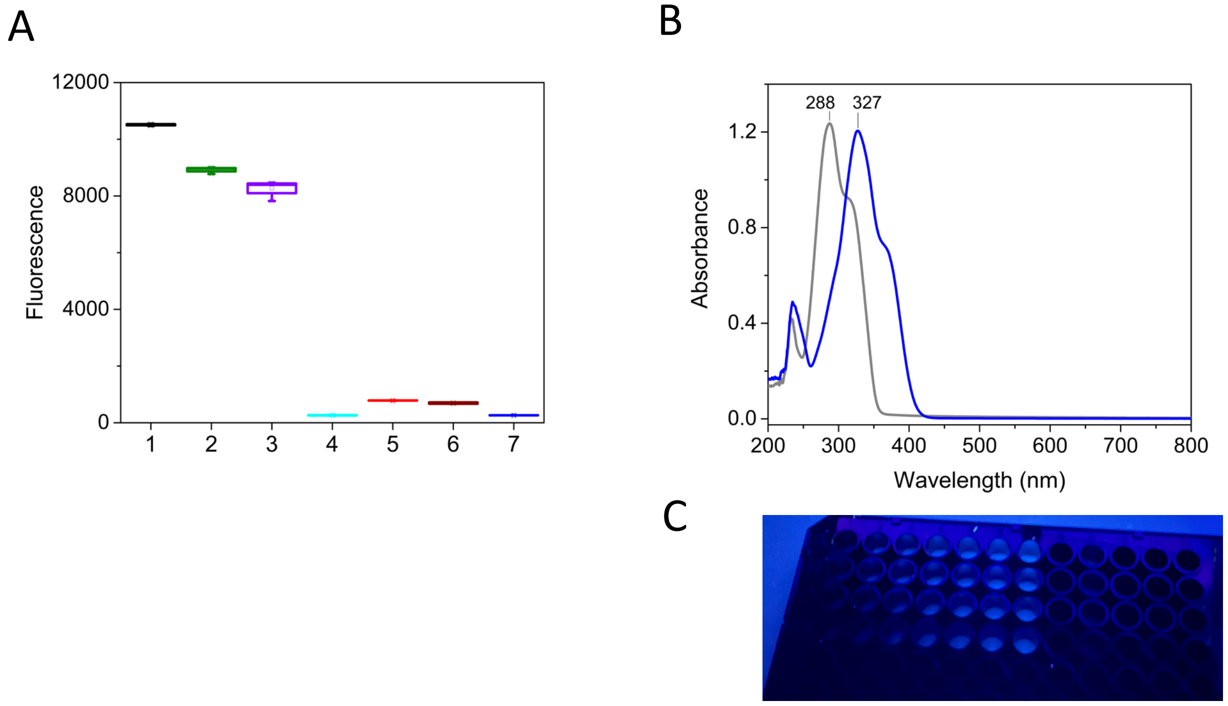

Figure S26. CBA assay at a fixed (10 μM) concentration of formate (F) (n=3). (A) Abscissa refers to sample numbers. $H_2O_2$ standard (10 μM) (black), (2) FDH2 + F (green),. FDH2 + F + SOD (10 U/mL) (violet), FDH2 + F + catalase (100 U/mL) (cyan), heat denatured FDH2 + F (red), FDH2 alone (brown), and FDH2 + F sans CBA (blue). 1.6 nM FDH2 was used. Unless specified otherwise, all samples contained buffer, DTPA, and CBA. (B) Electronic pectra of CBA (grey) and the product (7-hydroxy-coumarin, COH) (blue) derived from reacting CBA with 200 μM $H_2O_2$. (C) Representative image of a CBA assay plate under UV light.

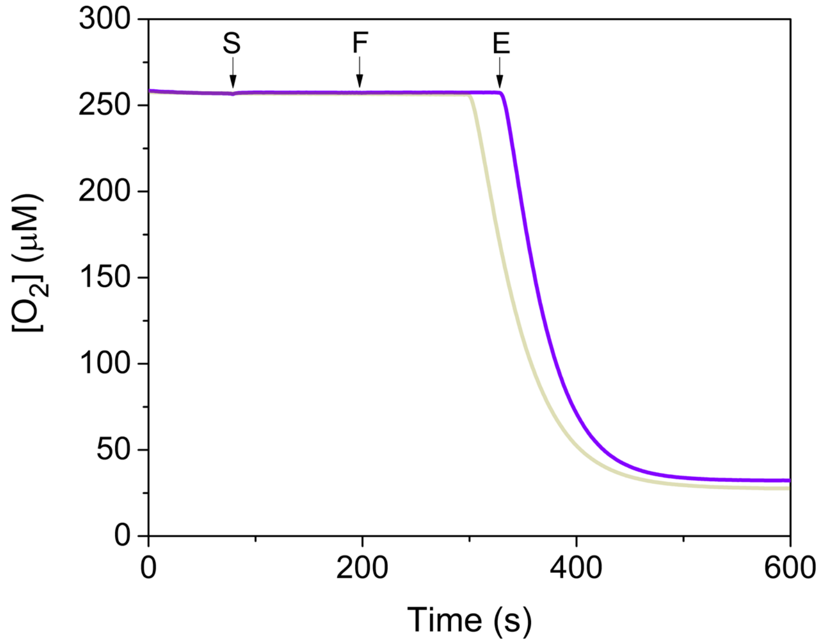

Figure S27. Rate of $O_2$ uptake by DvH-FDH2 remains largely unaffected by the presence of SOD. Purple and yellow traces were measured with and without SOD, respectively. E, enzyme, F, formate, and S, SOD.

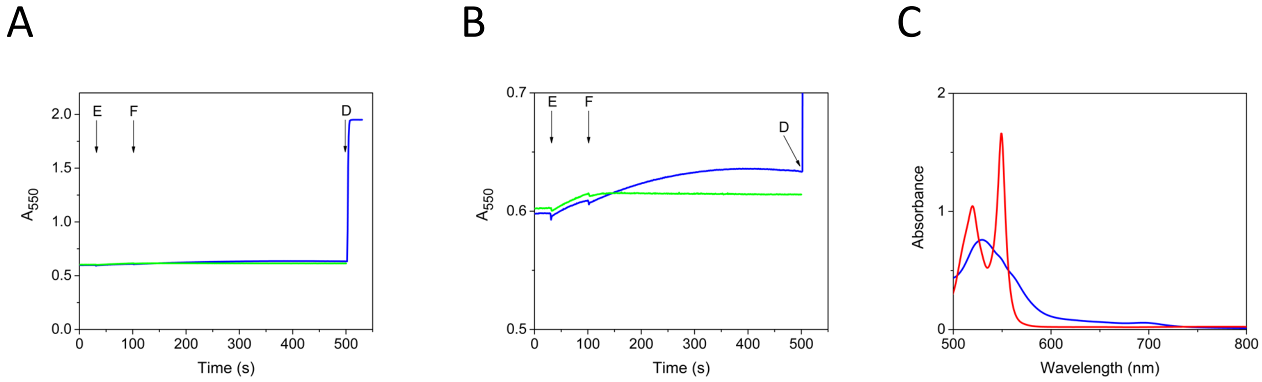

Figure S28. Partially acetylated equine cytochrome *c* is not significantly reduced during aerobic DvH-FDH2 catalysis. Points of addition of enzyme (E) and formate (F) are identified by arrows. Dithionite (D) addition at the end of the experiment is also shown as rference. (A) (B) Closeup view of panel A, (C) Electronic spectra of oxidized (blue) and reduced (red) acetylated cytochrome c. E + F + cytochrome *c* (blue), E + cytochrome c (no formate control) (green). $A_{550}$ refers to absorbance at 550 nm.

A B

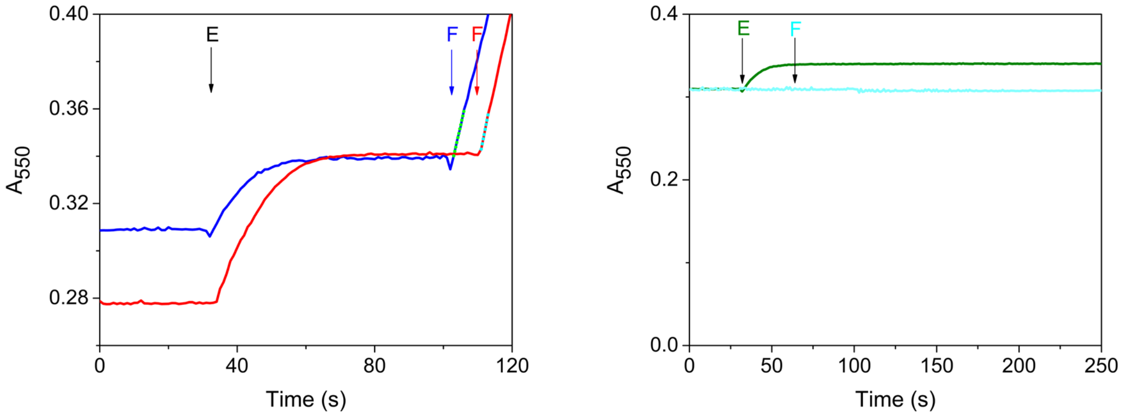

Figure 29. Kinetics of cytochrome c reduction by FDH2. (A) Enlarged view of Figure 7 depicting the early stages. Aerobic (blue) and anaerobic (red) conditions are shown. Green/cyan dots represent the region used in estimating the initial rates. (B) No formate (green) and no enzyme (cyan) controls.

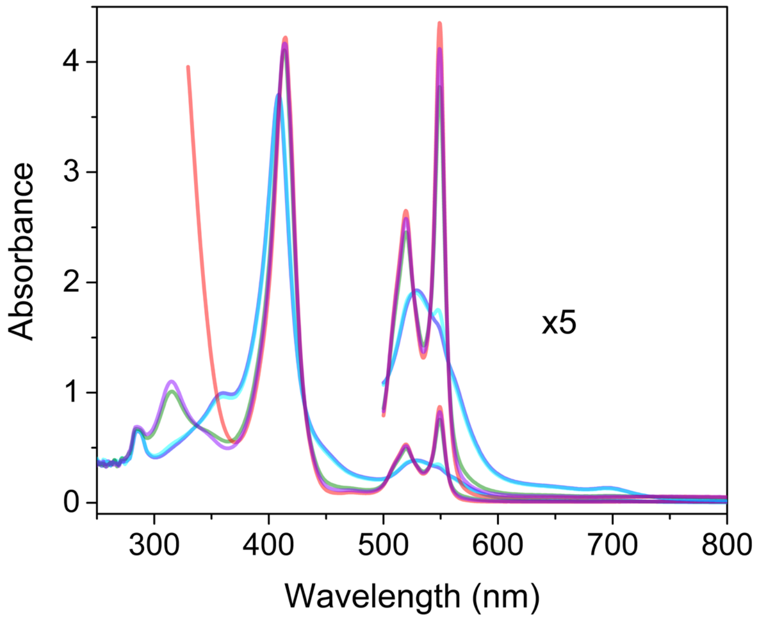

Figure S30. Direct reduction of equine cytochrome c during aerobic DvH-FDH2 catalysis. Electronic spectra of: Oxidized cytochrome c (blue), FDH2 + cytochrome c (cyan), FDH2 + cytochrome c + formate (green), FDH2 + cytochrome + 2x formate (violet), FDH2 + cytochrome c + 2x cytochrome c + dithionite (orange). Q-band and near-infrared region of the spectra have been magnified 5x to reveal the 695 nm band, which is a direct indicator of the functional integrity of cytochrome *c*. Photometric range of the Shimadzu UV-2600i spectrophotometer used in this measurement is ±5 absorbance units.

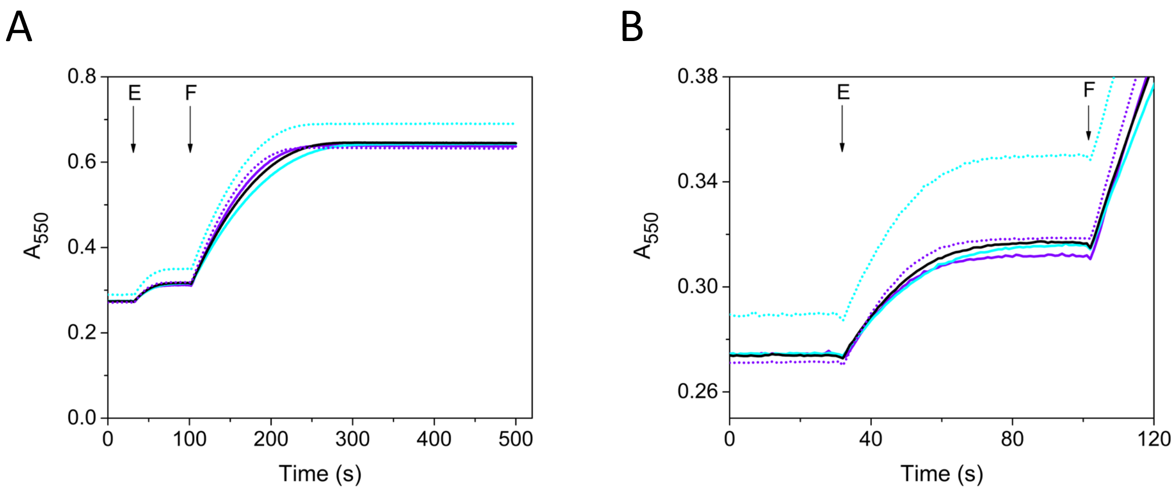

Figure S31. Effect of SOD and catalase on the aerobic kinetics of native equine cytochrome c reduction by FDH2. All reactions were performed in 50 mM Tris-HCl buffer pH 8 and 30 μM cytochrome c. E and F denote enzyme (1.6 nM) or formate (10 μM) additions. (A) +SOD (10 U/mL) (solid violet), + catalase (40 U/mL) (solid cyan), without SOD and catalase control (solid black), + SOD (100 U/mL) (dotted violet), + catalase (400 U/mL) (dotted cyan). (B) Enlarged view of panel A. There is a small absorbance change at 550 nm upon mixing FDH2 and oxidized cytochrome. This is also visible in Figures S29 and S30. We do not have an explanation for this effect.

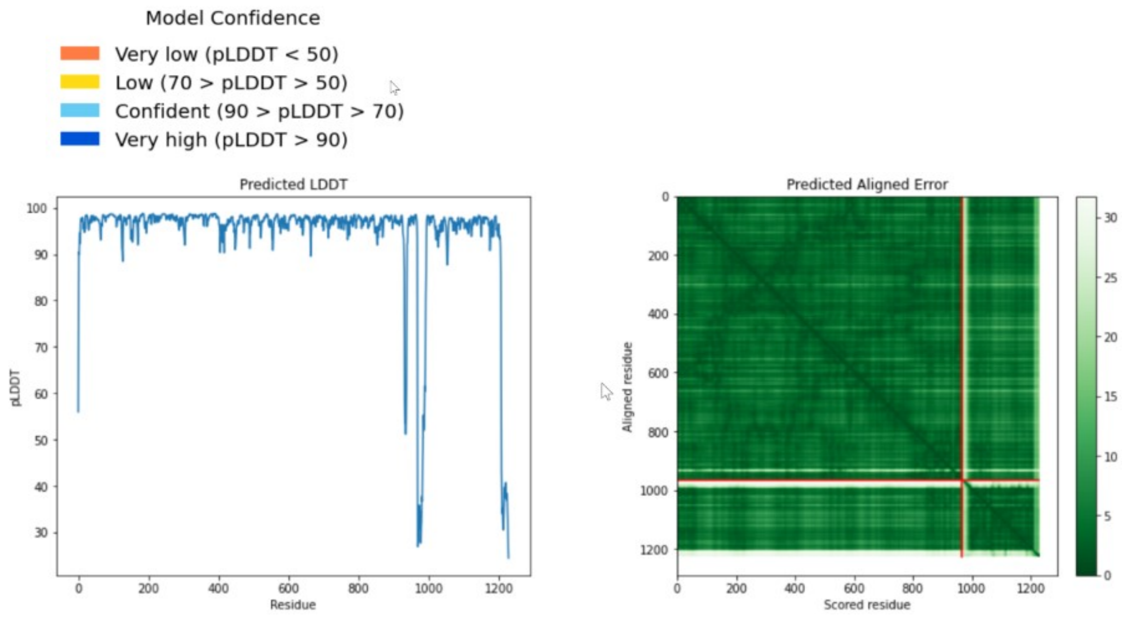

Figure S32. Confidence metrics associated with AlphaFold2.1 structure prediction. See text for details. The intersecting lines on the right panel is a consequence of protein boundaries introduced by the use of two independent sequences (large and small subunits of FDH2) for predicting the structure of FDH2 heterodimer. pLDDT spike near residue 1000 (left panel) is also a result of the same.

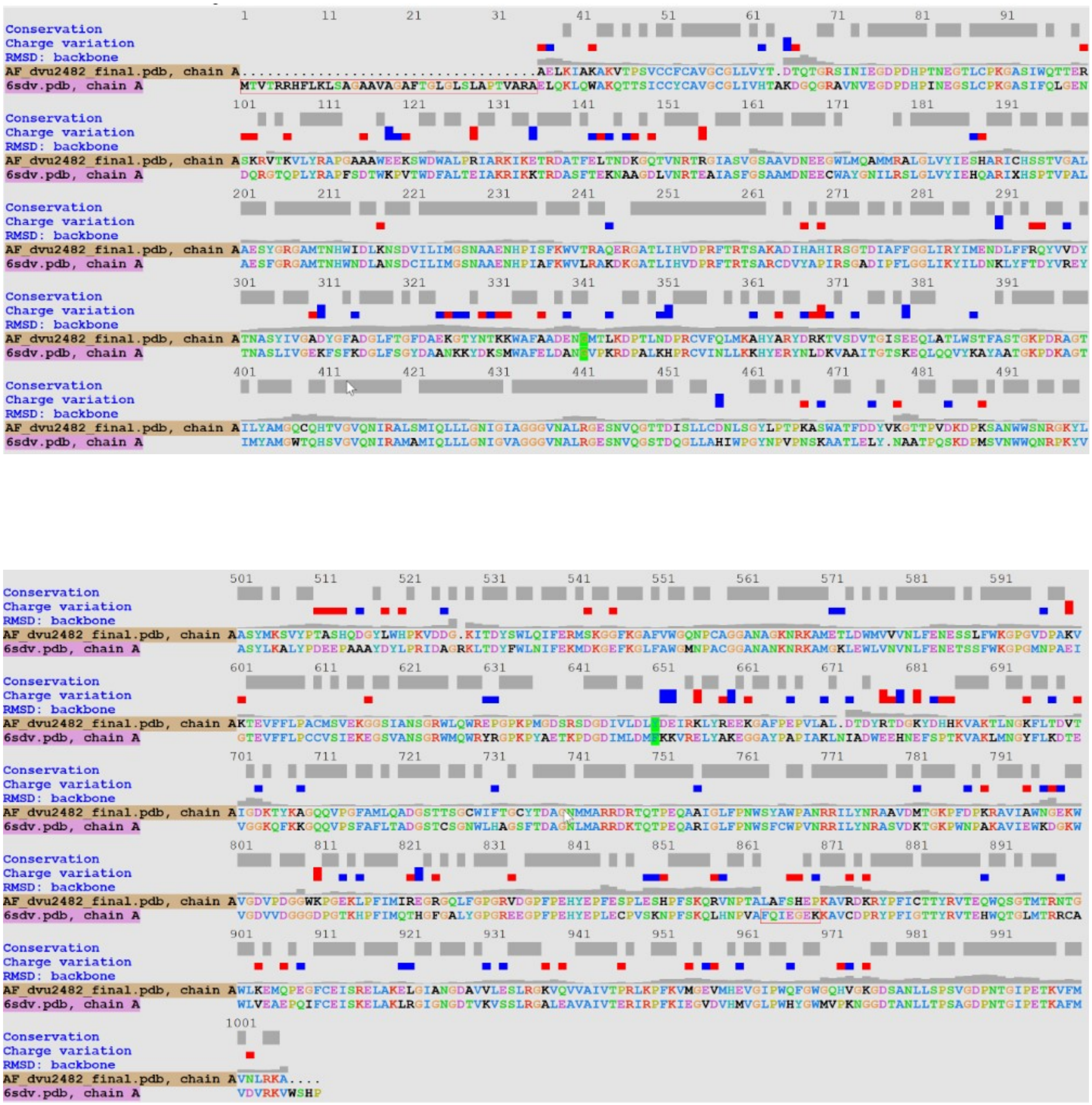

Figure 33. Backbone RMSD variations between the large subunits of FDH2 and FDH1 at the single residue level.

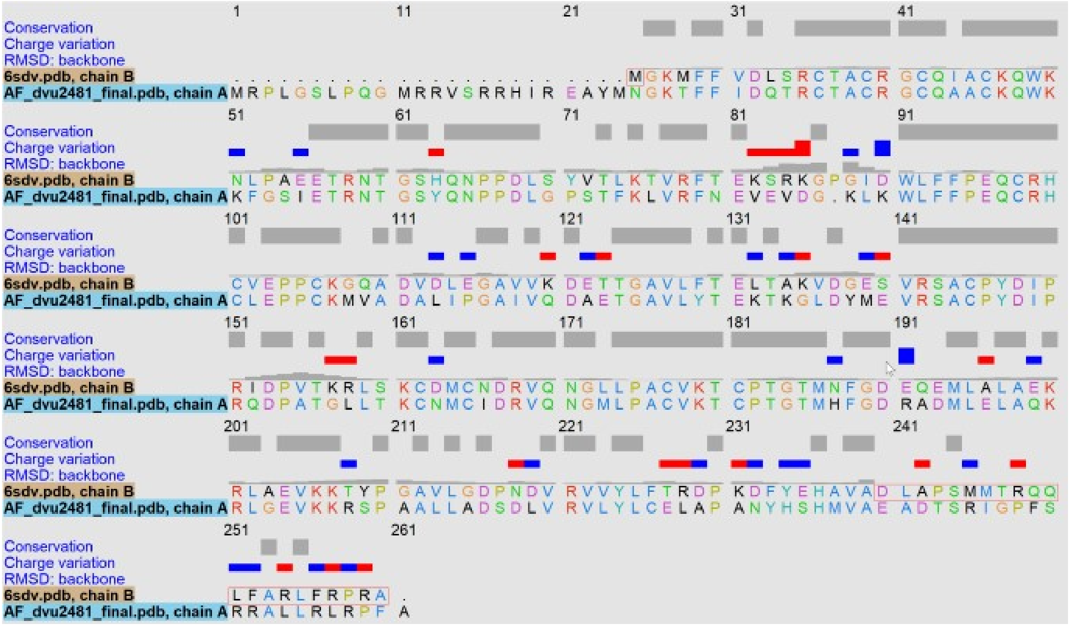

Figure 34. Backbone RMSD variations between the small subunits of FDH2 and FDH1 at the single residue level.

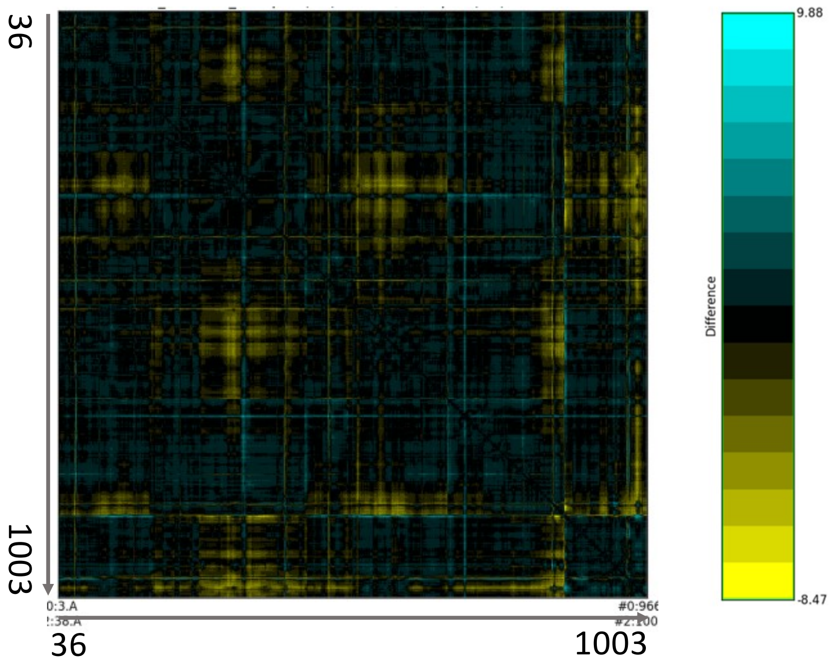

Figure 35. Difference residue-residue distance maps of FDH2:FDH1 large subunit pair. Zero (black), positive (cyan), and negative (yellow) differences are shown.

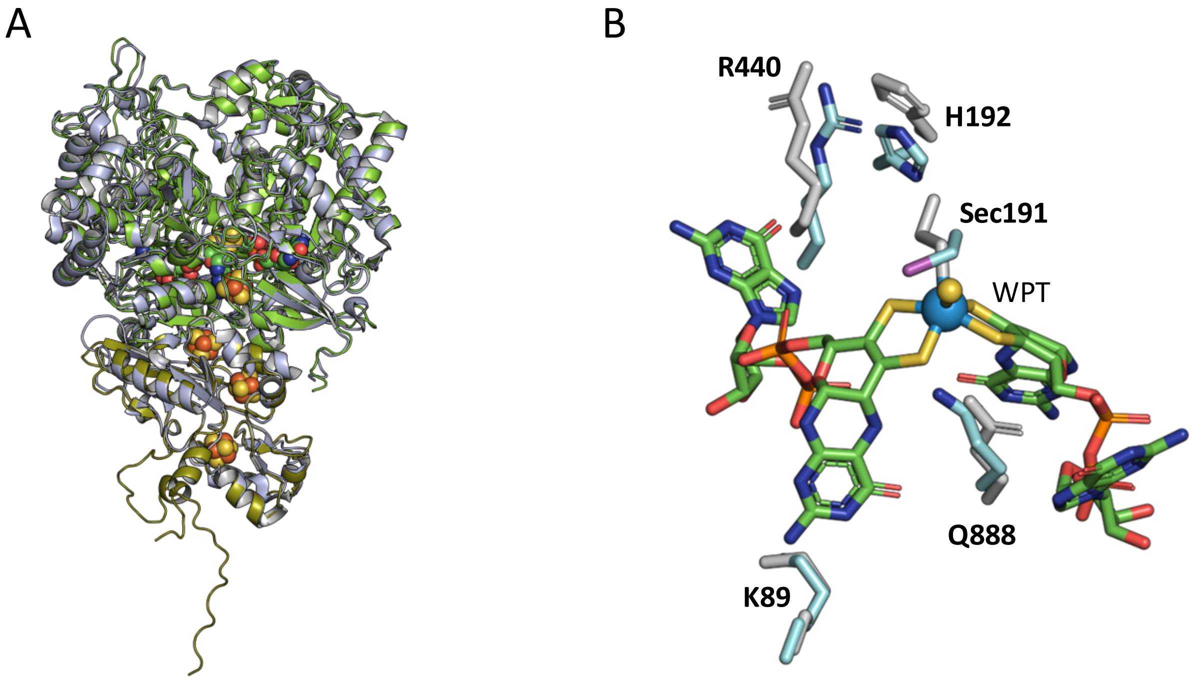

Figure 36. Structural comparison of FDH2 and FDH1 heterodimers. (A) Superposition of the two proteins. FDH1 is in grey. Large and small subunits of FDH2 are shown in green and olive, respectively. (B) Overlay of the invariant active site residues. A bond between Sec191 and W is not shown for clarity.

Protocol S1. Medium MOYLS4 for cultivating *Desulfovibrio* strains.

**Materials**

Serum bottles 100 mL (Wheaton)

Media bottles 500mL, 1000 mL, 2000mL (Gibco or Duran); caps modified in-house to include a 7 mm center bore.

Butyl rubber stoppers (Chemglass CLS-4209-14)

Neoprene stoppers size 6 (RPI-259100-6) cut to 2/3 original height

Magnetic stirrer

**Equipment**

Autoclave

Gas manifold

pH meter

**Reagents**

Magnesium chloride hexahydrate Sigma-Aldrich M9272 BCBT8684

Ammonium Chloride-Sigma A9434-1KG SLBS1591V

Calcium Chloride dihydrate- Acros 207780010 A014020501

Tris HCl –Fisher BP153-500 200711

Iron (III) Chloride Hexahydrate- Acros 423705000 lot A013817901

EDTA- Fisher BP120-1 lot 055880

Yeast Extract- Sigma-Aldrich 92144-5KG-F BCBQ9331V

Thioglycolate-Sigma T0632-25g STBH2638

Sodium DL-lactate 60% syrup-Sigma L1375-500mL SLBR4194V

Sodium Sulfate-Sigma-Aldrich 239313-2.5KG SLBT9903

Potassium phosphate dibasic-Fluka 60353 1167325

Sodium phosphate monobasic- Fluka 71505 1203314

Manganese Chloride tetrahydrate- Acros Organics 2058950000 A012429101

Cobalt Chloride hexahydrate Sigma C3169 70K3698

Zinc Chloride hexahydrate Sigma Z-4875 20K0264

Sodium molybdate dehydrate Sigma M1003 085K0098

Boric Acid Sigma B-7660 042K0150

Nickel Chloride ICN 155825 8649C

Copper Chloride dehydrate Sigma C-6917 121K0014

Sodium Selenate Sigma S8295-25G SLBD3716V

Sodium Tungstate dehydrate Aldrich 223336-5g MKBV3962V

Biotin Sigma-aldrich B4501-1g SLBS3069V

Folic Acid Sigma F7876-1g SLBN1618V

Pyridoxine HCl RPI P50240-10.0 10874462

Thiamine HCl Sigma T-4625 062K0103

Riboflavin Sigma R-4500 072K0887

Nicotinic Acid Sigma N-4126 052K0200

DL Pantothenic Acid Sigma P-2250 013K0583

4-Aminobenzoic Acid Sigma A9878-5G MKBZ3723V

Lipoic Acid Sigma T5625-500g SLBS2381V

Choline Chloride Fisher AC110290500 A0400838

Vitamin B12 Sigma V-2876 112K0646

Sodium hydroxide Fisher BP359-500 lot 201811

Sodium Resazurin –Aldrich 199303-1G MKBP2801V

**Solutions**

| Trace Metals Stock solution(1L) | (g) |
|---|---|
| Manganese Chloride tetrahydrate | 0.5 |
| Cobalt chloride hexahydrate | 0.3 |
| Zinc Chloride | 0.2 |
| Sodium molybdate dihydrate | 0.05 |
| Boric Acid | 0.02 |
| Nickel chloride | 0.09 |
| Copper chloride dihydrate | 0.002 |
| Sodium selenate | 0.006 |
| Sodium tungstate dihydrate | 0.008 |

| Vitamins Stock solution (10x) | (g) |
|---|---|
| Biotin | 0.02 |
| Folic Acid | 0.02 |
| Pyridoxine HCl | 0.1 |
| Thiamine HCl | 0.05 |
| Riboflavin | 0.05 |
| Nicotinic Acid | 0.05 |
| DL Pantothenic Acid | 0.05 |
| 4-Aminobenzoic Acid | 0.05 |
| Lipoic Acid | 0.05 |
| Choline Chloride | 2 |
| Vitamin B12 | 0.01 |

pH 7.0 with KOH. Filter sterilize and store at -20°C.

Vitamins solution working stock (1x)

10 mL of 10x filter sterilized into 90 mL of anaerobic ($N_2$ sparged) water. Stored at 4 C and in the dark.

Sodium resazurin 0.1%

0.1 g

100 mL water

0.5 M EDTA

93 g EDTA

pH to 8.0 with NaOH

volume to 500mL with MilliQ water

FeCl/EDTA solution 125mM/250mM- 4.8 mL-

0.162 g Iron chloride Hexahydate

2.4 mL water

2.4 mL 0.5 M EDTA

| Potassium/Sodium Phosphate 1M | per L |
|---|---|
| Pottasium Phosphate dibasic | 87g |
| Sodium phosphate monobasic | 78g |

MOYLS4 Medium 1L

To ~800 mL MiliQ water add:

1.6 g Magnesium Chloride hexahydrate

1.06 g Ammonium Chloride

0.088 g Calcium Chloride dihydrate

2 mL Potassium/Sodium Phosphate solution

6 mL Mo-Trace elements

4.73 g Tris HCl (or 15 mL 2M –pH 7.2)

1 g yeast extract

11.2 mL sodium lactate 60%

4.26 g Sodium sulfate

0.16 mL sodium resazurin 0.1%

0.48 mL Iron chloride/EDTA 125mM/250mM (**see Methods**)

Bring to 1 L and pH to 7.2 with 4M sodium hydroxide.

Bubble under $N_2$ gas with stirring for 1 hr

Add 0.14 g Sodium Thioglycolate

Cap bottle with number 6 neoprene stopper and medium bottle cap with center bore hole

Or distribute under $N_2$ to serum bottles (50 mL per 100 mL bottle) and cap with butyl rubber stopper and aluminum crimp seal.

Autoclave liquid cycle. Cool to room temperature.

To finish medium add 1x vitamin stock at 0.5 mL per 50 mL, just prior to inoculation by sterile anaerobic transfer. Add desired antibiotics at this time as well.

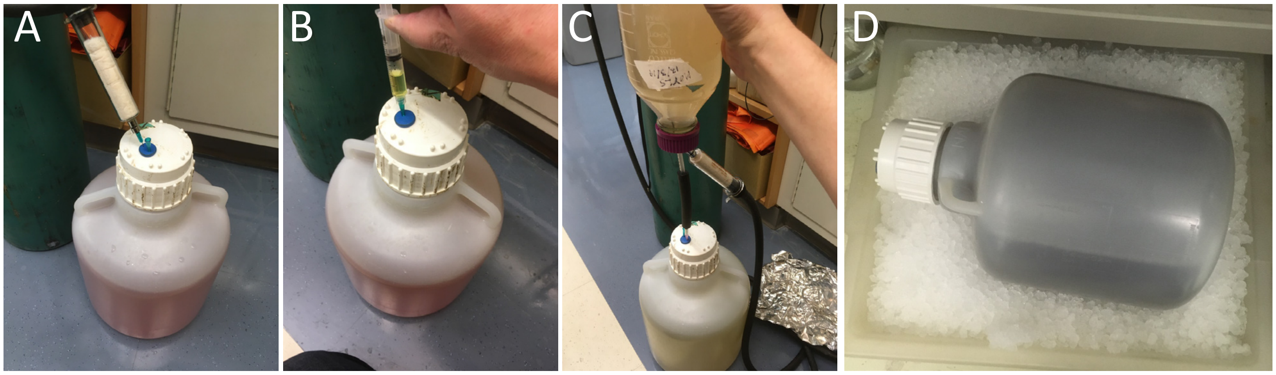

Figure S37. Experimental workflow for facile medium scale cultivation of DvH. To a 10 L carboy containing pre-warmed sterile MOYLS4 media, the following were added in sequence using a sterile syringe: vitamins, iron chloride/EDTA solution, and spectinomycin. The carboy lid was closed tightly. Carboy was gassed with $N_2$ through stopper to remove air in the headspace; vented with a 23 gauge needle (A). Injected sodium sulfide by sterile anaerobic syringe transfer (B). Mix by rolling or shaking the carboxy and incubate at 37 C until resazurin turned colorless. Connected carboy to a 500 mL culture bottle using a transfer line fitted with 18 gauge needle. Transferred 250 – 500 mL of active culture to carboy while venting with a 23 gauge needle (C). Incubated at 37 C. Monitored growth via $OD_{550}$ until it plateaus and subsequently chilled the carboy on ice (D).

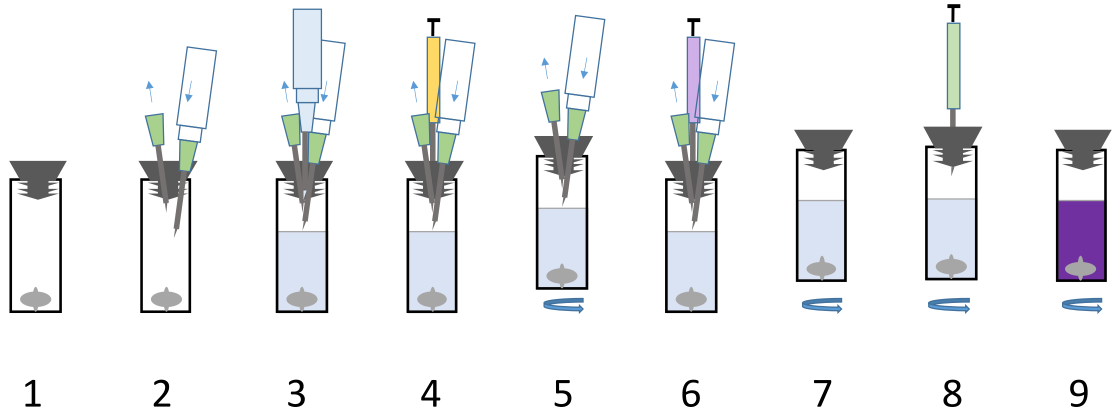

Figure S38. Experimental workflow for BV assay. Setup cuvette (1), gas with argon (2), fill with reaction mix (3), inject GO + catalase (4), gas with argon while mixing for 20 minutes (5), transfer to spectrophotometer and continue to stir (7), initiate a kinetics run by injecting formate (8), continue data collection until end (9).

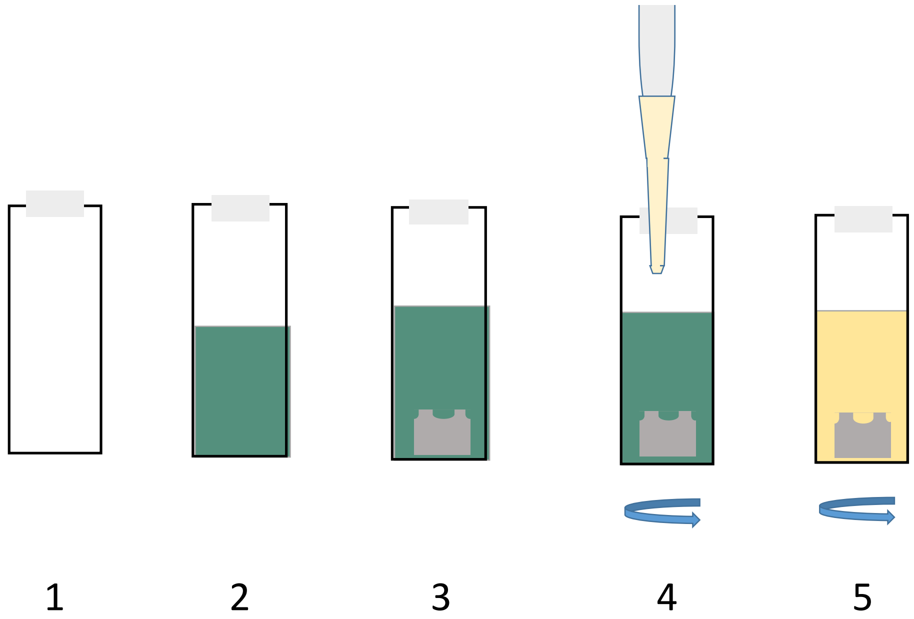

Figure S39. Experimental workflow for aerobic PES/DCPIP assay. Prepare cuvette (1), fill with reaction mix (2), add stirrer (3), initiate kinetics by the addition of DvH-FDH2 while the contents of the cuvette are being mixed(4), and continue data collection till end (5).

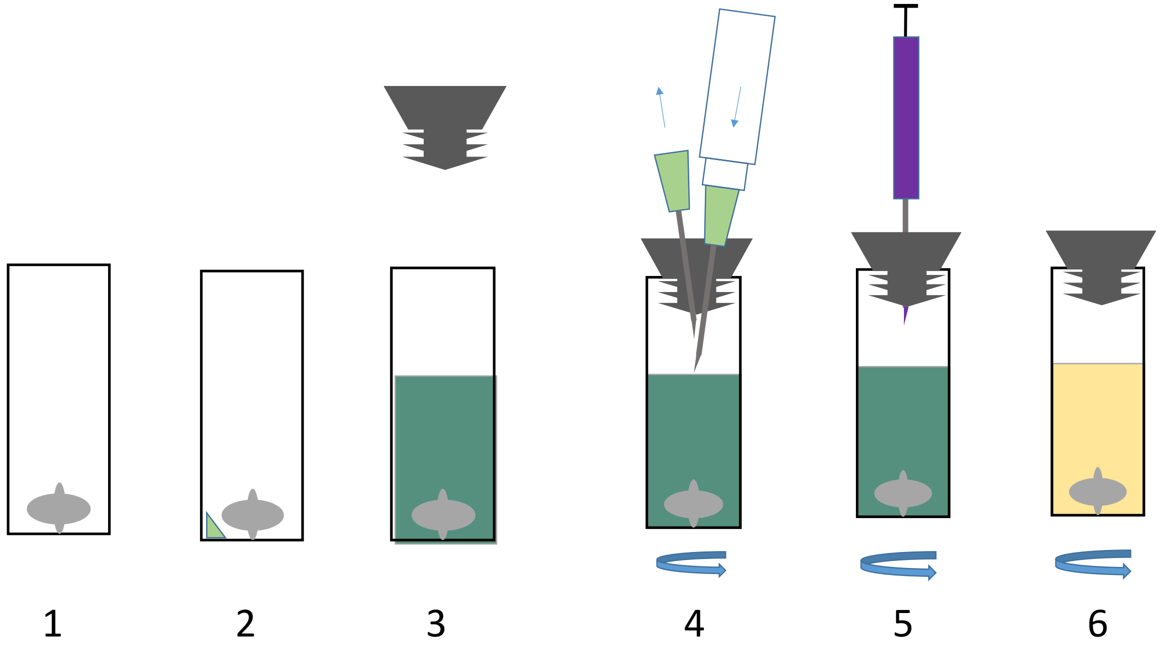

Figure S40: Experimental workflow for anaerobic PES/DCPIP assay. Setup cuvette (1,2). add reaction mix (3), gas with argon (4), start the reaction by injecting DvH-FDH2 (5), and collect data until the reaction goes to completion (6).

### Supplement Citations

(1) Keller, K. L.; Bender, K. S.; Wall, J. D., Development of a markerless genetic exchange system for Desulfovibrio vulgaris Hildenborough and its use in generating a strain with increased transformation efficiency. *Appl Environ Microbiol* **2009,** *75*, 7682-91.

(2) Parks, J. M.; Johs, A.; Podar, M.; Bridou, R.; Hurt, R. A., Jr.; Smith, S. D.; Tomanicek, S. J.; Qian, Y.; Brown, S. D.; Brandt, C. C.; Palumbo, A. V.; Smith, J. C.; Wall, J. D.; Elias, D. A.; Liang, L., The genetic basis for bacterial mercury methylation. *Science* **2013,** *339*, 1332-5.

(3) Teufel, F.; Almagro Armenteros, J. J.; Johansen, A. R.; Gislason, M. H.; Pihl, S. I.; Tsirigos, K. D.; Winther, O.; Brunak, S.; von Heijne, G.; Nielsen, H., SignalP 6.0 predicts all five types of signal peptides using protein language models. *Nat Biotechnol* **2022**.

(4) Axley, M. J.; Grahame, D. A., Kinetics for formate dehydrogenase of Escherichia coli formate-hydrogenlyase. *J Biol Chem* **1991,** *266*, 13731-6.

(5) Maia, L. B.; Fonseca, L.; Moura, I.; Moura, J. J., Reduction of Carbon Dioxide by a Molybdenum-Containing Formate Dehydrogenase: A Kinetic and Mechanistic Study. *J Am Chem Soc* **2016,** *138*, 8834-46.

(6) Johnson, K. A., New standards for collecting and fitting steady state kinetic data. *Beilstein J Org Chem* **2019,** *15*, 16-29.

(7) Olp, M. D.; Kalous, K. S.; Smith, B. C., ICEKAT: an interactive online tool for calculating initial rates from continuous enzyme kinetic traces. *BMC Bioinformatics* **2020,** *21*, 186.

(8) Oliveira, A. R.; Mota, C.; Mourato, C.; Domingos, R. M.; Santos, M. F. A.; Gesto, D.; Guigliarelli, B.; Santos-Silva, T.; Romão, M. J.; Cardoso Pereira, I. A., Toward the Mechanistic Understanding of Enzymatic CO2 Reduction. *ACS Catalysis* **2020,** *10*, 3844-3856.

(9) Lissolo, T.; Pulvin, S.; Thomas, D., Reactivation of the hydrogenase from Desulfovibrio gigas by hydrogen. Influence of redox potential. *J Biol Chem* **1984,** *259*, 11725-9.

(10) Tagawa, K.; Arnon, D. I., Oxidation-reduction potentials and stoichiometry of electron transfer in ferredoxins. *Biochim Biophys Acta* **1968,** *153*, 602-13.

(11) Jones, R. W., The topography of the membrane-bound hydrogenase of Escherichia coli explored by non-physiological electron acceptors [proceedings]. *Biochem Soc Trans* **1979,** *7*, 724-5.

(12) McKellar, R. C.; Sprott, G. D., Solubilization and properties of a particulate hydrogenase from Methanobacterium strain G2R. *J Bacteriol* **1979,** *139*, 231-8.

(13) Graf, E.-G.; Thauer, R. K., Hydrogenase from Methanobacterium thermoautotrophicum, a nickel-containing enzyme. *FEBS Lett* **1981,** *136*, 163-169.

(14) Arp, D. J.; Burris, R. H., Kinetic mechanism of the hydrogen-oxidizing hydrogenase from soybean nodule bacteroids. *Biochemistry* **1981,** *20*, 2234-40.

(15) Axley, M. J.; Grahame, D. A.; Stadtman, T. C., Escherichia coli formate-hydrogen lyase. Purification and properties of the selenium-dependent formate dehydrogenase component. *J Biol Chem* **1990,** *265*, 18213-8.

(16) Rosner, B. M.; Schink, B., Purification and characterization of acetylene hydratase of Pelobacter acetylenicus, a tungsten iron-sulfur protein. *J Bacteriol* **1995,** *177*, 5767-72.

(17) Mukund, S.; Adams, M. W., Glyceraldehyde-3-phosphate ferredoxin oxidoreductase, a novel tungsten-containing enzyme with a potential glycolytic role in the hyperthermophilic archaeon Pyrococcus furiosus. *J Biol Chem* **1995,** *270*, 8389-92.

(18) Costa, C.; Teixeira, M.; LeGall, J.; Moura, J. J. G.; Moura, I., Formate dehydrogenase from Desulfovibrio desulfuricans ATCC 27774: isolation and spectroscopic characterization of

the active sites (heme, iron-sulfur centers and molybdenum). *JBIC Journal of Biological Inorganic Chemistry* **1997,** *2*, 198-208.
(19) Gross, R.; Simon, J.; Kroger, A., Periplasmic methacrylate reductase activity in Wolinella succinogenes. *Arch Microbiol* **2001,** *176*, 310-3.
(20) de Bok, F. A.; Roze, E. H.; Stams, A. J., Hydrogenases and formate dehydrogenases of Syntrophobacter fumaroxidans. *Antonie Van Leeuwenhoek* **2002,** *81*, 283-91.
(21) Selwyn, M. J., A simple test for inactivation of an enzyme during assay. *Biochim Biophys Acta* **1965,** *105*, 193-5.
